# Mitochondrial Retrograde Signaling in *Arabidopsis thaliana*: heterogenous, spatial and polarised aspects

**DOI:** 10.64898/2026.02.08.704520

**Authors:** Yanqiao Zhu, Minxuan Li, Jia Hao Li, Rijad Sarić, Feiran Han, Xinxin Zhou, Xiachao Wang, Yuhua Liu, Chi Zhang, Huixia Shou, Ghazanfar Abbas Khan, Mathew G. Lewsey, James Whelan

**Affiliations:** State Key Laboratory of Plant Environmental Resilience, College of Life Science, Zhejiang University, Hangzhou, Zhejiang 310058, P.R. China; The Provincial International Science and Technology Cooperation Base on Engineering Biology, International Campus of Zhejiang University, Haining, Zhejiang 314400, China; La Trobe Institute for Sustainable Agriculture and Food, AgriBio, La Trobe University, Bundoora, VIC 3086, Australia; Australian Research Council Industrial Transformation Research Hub for Protected Cropping, La Trobe University, Bundoora, VIC 3086, Australia; LC-Bio Technology (Hangzhou) Co., Ltd., Hangzhou, 310000, China; School of Life and Environmental Sciences & Centre for Sustainable Bioproducts, Deakin University, Waurn Ponds, VIC 3216, Australia; La Trobe Institute for Molecular Sciences, La Trobe University, Bundoora, VIC 3086, Australia; Australian Research Council Centre of Excellence in Plants for Space, AgriBio, La Trobe University, Bundoora, VIC 3086, Australia

**Keywords:** Mitochondria retrograde signaling, Single nuclei and Spatial RNA-seq, Cell and cluster specificity, Polarity, Transcription factors

## Abstract

Organelle retrograde signaling studies have exclusively relied on bulk tissues, obscuring cell-type specificity, spatial organization, and heterogeneity within cell types. Here, we use single-nucleus and spatial RNA sequencing to define the cellular and spatial architecture of mitochondrial stress signaling in *Arabidopsis thaliana*. Inhibition of the mitochondrial electron transport chain by antimycin A and myxothiazol rapidly redistributed nuclei among transcriptional states within epidermal, leaf pavement, and mesophyll lineages, revealing that mitochondrial dysfunction reshapes cell identity trajectories rather than eliciting a uniform response. Stress-associated clusters were already present at low frequency in untreated samples, but the numbers of cells increased following stress treatments, indicating that stress increases the proportion of nuclei occupying pre-existing, primed transcriptional states. Gene-level analyses revealed distinct temporal dynamics and variable cell-state penetrance of the canonical mitochondrial stress markers and identified broadly responsive genes absent from earlier marker sets. Spatial transcriptomics resolved tissue-scale responses, spanning from pan-tissue induction to cell-type- and cluster-restricted activation, and uncovered pronounced adaxial–abaxial polarity in gene expression.

Cell identity and spatial inferences were supported by extensive experimental validation using a custom 465-probe 10x Xenium panel and 52 promoter-GFP reporter lines. Together, these data provide a high-resolution framework for organelle-to-nucleus signaling and a resource of cell-type markers and spatial maps to dissect mitochondrial stress signaling and its integration with developmental and environmental pathways.

## INTRODUCTION

Mitochondria are not only the powerhouses of plant cells but also critical hubs for stress signaling. These organelles actively communicate their functional status to the nucleus through retrograde signaling pathways (Meng et al., 2019). In concert with anterograde (nucleus-to-organelle) controls, mitochondrial retrograde signaling coordinates organelle function with whole-plant needs (Ng et al., 2014). When mitochondrial function is perturbed, such as by electron transport chain disruption, intracellular signals like reactive oxygen species (ROS) accumulate and trigger changes in nuclear gene expression (Meng et al., 2019). This mitochondrial retrograde response (MRR) constitutes an adaptive program to maintain cellular homeostasis under stress.

Over the past two decades, key components of the plant MRR pathway have been elucidated. In *Arabidopsis thaliana*, a NAC-domain transcription factor, ANAC017, has emerged as a master regulator of mitochondrial stress signaling (Ng et al., 2013a). ANAC017 is an endoplasmic reticulum (ER) associated protein that becomes activated upon mitochondrial dysfunction, via proteolytic cleavage, allowing its amino-terminal fragment to translocate to the nucleus (Eysholdt-Derzsó et al., 2023). Once in the nucleus, ANAC017 binds to target promoters and reprograms gene expression. One direct target is ALTERNATIVE OXIDASE 1A (*AOX1a*), that helps maintain electron flow and reduce ROS accumulation when the main respiratory chain is impaired (Meng et al., 2019; Millar et al., 2011). *AOX1a* is widely used as a hallmark of the MRR, as its robust transcriptional induction signifies an active retrograde response (Vanlerberghe and McIntosh, 1997). A variety of NAC transcription factors, including ANAC013, operate downstream of ANAC017 to coordinate a wider transcriptional program termed the mitochondrial dysfunction response (MDR), which includes genes responsive to mitochondrial impairment regardless of protein localization (De Clercq et al., 2013). Many MDR genes overlap with the mitochondrial stress response (MSR), a defined set of nuclear genes encoding mitochondrialZItargeted proteins upregulated during organelle stress (Van Aken et al., 2009). Additional transcription factors including ABI4 (ABA INSENSITIVE 4) and members of the WRKY, MYB, bZIP, and ERF families function as regulatory hubs that integrate mitochondrial signals into wider stress and developmental networks (Giraud et al., 2009; Li et al., 2025; Pedrotti et al., 2018; Racca et al., 2022; Van Aken et al., 2013; Vanderauwera et al., 2012; Xie et al., 2023). The MRR is also closely linked with hormonal pathways and signals originating from other organelles, forming a complex regulatory landscape (Broad et al., 2024; He et al., 2023; He et al., 2022).

Despite major advances in identifying components of MRR, our understanding remains constrained by the limits of bulk transcriptomic approaches. Most studies have relied on whole-seedling analyses and loss-of-function mutants to infer regulatory roles (De Clercq et al., 2013; Ng et al., 2013a). While these approaches identified core regulators and gene sets, they obscure cellular heterogeneity by averaging gene expression across complex tissues (Li et al., 2025). As a result, the fundamental question of whether there is a uniform MRR across all cells or, instead, if responses are tissue and/or cell specific, remains unanswered. Does the MRR operate uniformly across all cells, or is it activated more strongly in specific tissues or cells? Preliminary evidence from other stress responses suggests the latter. For instance, under mild drought, opposing transcriptional changes occur in mesophyll and epidermal cells, a pattern undetectable in bulk data (Sheng et al., 2025). Similarly, laser capture microdissection of Arabidopsis leaves treated with antimycin A revealed that mitochondrial stress responses differ across tissue layers, with epidermal cells showing enhanced expression of specific MRR genes compared to mesophyll (Berkowitz et al., 2021). These findings suggest that mitochondrial signalling may follow similarly complex, cell type–specific trajectories.

Single-cell and single-nucleus RNA sequencing (sc/snRNA-seq) provide the resolution needed to dissect transcriptional responses at cellular scale, enabling the identification of rare or specialized cell populations involved in organellar signaling (Guo et al., 2025; Lee et al., 2025; Wang et al., 2025b). However, dissociative single-cell methods inherently disrupt tissue architecture, preventing direct assignment of gene expression patterns to their native spatial context. Spatial transcriptomics overcomes this limitation by preserving RNA location *in situ*, enabling direct visualization of where genes are expressed within intact organs (Peirats-Llobet et al., 2023). This spatial resolution reveals expression gradients, tissue-layer specificity, and transcriptional hotspots that would otherwise be undetectable. While a few recent studies have applied these technologies to stress contexts such as drought and pathogen challenge (Sheng et al., 2025; Tenorio Berrío et al., 2025; Wang et al., 2025a), no study has yet mapped organelle retrograde signalling at single-cell or spatial resolution.

Here, we integrate snRNA-seq and spatial transcriptomics to define the cellular and spatial architecture of the MRR in *Arabidopsis thaliana* seedlings treated with mitochondrial electron transport inhibitors antimycin A and myxothiazol. By resolving transcriptional responses across over 600,000 nuclei and visualizing spatial gene expression at subcellular resolution, we identify the cell types and tissue domains that orchestrate the retrograde response. This work reveals substantial heterogeneity in MRR activity, uncovers putative sensor or primed cells, and establishes a spatially organized framework for organelle-to-nucleus signaling in multicellular plant tissues.

## Results

### Establishing a cell-wide view of mitochondrial dysfunction

To define the cellular landscape of the mitochondrial dysfunction response, we profiled Arabidopsis shoot tissue by snRNA-seq across a time course following inhibition of the mitochondrial electron transport chain. Seedlings were treated with antimycin A or myxothiazol for 1, 3, 6, and 12 h, with time-matched mock controls to control for diurnal effects and spray-induced transcriptional changes (Figure 1A) (Van Aken et al., 2016; Xu et al., 2019). Unsupervised clustering across all samples resolved 28 transcriptionally distinct clusters (Figure 1B, Supplemental Table 2) (Chen et al., 2021; Jin et al., 2022). Initial annotation based on computational marker resources identified four epidermal clusters, three leaf pavement cell clusters, six mesophyll clusters, two bundle sheath clusters, and single clusters corresponding to phloem, phloem parenchyma, companion cells, leaf guard cells, and cell cycle states (G2/M and S phase), together with seven clusters that could not be assigned using computational approaches alone (Figure 1B, Supplemental Table 2).

**Figure 1.**
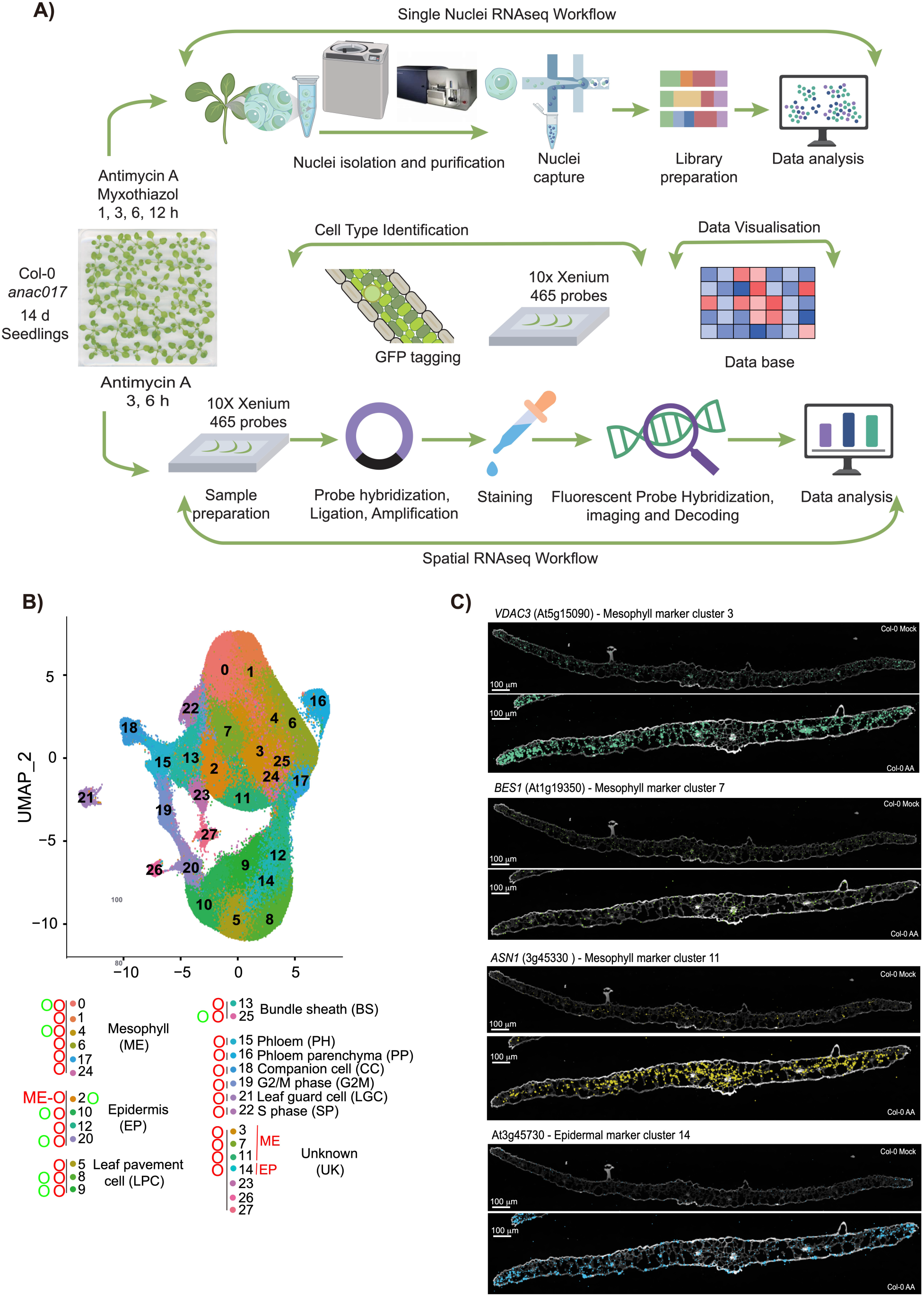
Experimental design, cell type clustering and marker gene expression. **(A)** Overview of the experimental workflow. Fourteen-day-old *Arabidopsis thaliana* Col-0 and *anac017* seedlings were treated with antimycin A or myxothiazol for 1, 3, 6 or 12 h for single-nucleus RNA sequencing (snRNA-seq), or with antimycin A for 3 and 6 h for spatial RNA-seq (spRNA-seq). For snRNA-seq, nuclei were isolated and purified by flow cytometry, captured on the 10x Genomics platform, used for library preparation and sequenced, followed by computational analysis. Cell type identities were refined using a combination of public marker resources, a custom 10x Xenium panel containing 465 probes, and 52 promoter GFP lines. Processed datasets were deposited in an interactive database for cluster- and cell-type–level visualisation (https://arabidopsismrsdb.com/). For spRNA-seq, sections from treated seedlings were processed for 10x Xenium analysis: sample preparation, probe hybridisation, ligation and amplification, tissue staining, fluorescent probe hybridisation and imaging, decoding and downstream data analysis. **(B)** UMAP projection of >600,000 nuclei from Col-0 and *anac017* across treatments and time points, coloured by unsupervised cluster identity (0–27). Major cell types are assigned based on marker expression and experimental validation: mesophyll (ME), epidermis (EP), leaf pavement cells (LPC), bundle sheath (BS), phloem (PH), phloem parenchyma (PP), companion cells (CC), leaf guard cells (LGC), G2/M phase (G2M), S phase (SP) and unknown (UK). Cell type assignments were confirmed using 10x Xenium spatial localisation (red circles) and promoter–GFP lines (green circles); clusters originally annotated as unknown are labelled UK in black text, and, with experimentally supported identities indicated in red. **(C)** Examples of the spatial patterns obtained with predicted marker genes. Shown here are examples of marker genes for clusters that were designated as unknown using the various plant cell marker databases. *VDAC3* (*Voltage Dependent Anion Channel 3*) (At5g15090) for cluster 3, *BES1* (*BRI1* (*BRASSINOSTEROID INSENSITIVE 1*) *EMS SUPPRESSOR 1)* (At19350) for cluster 7, and *ASN1* (*GLUTAMINE-DEPENDENT ASPARAGINE SYNTHASE 1*) (At3g37340) for cluster 11, a gene at locus At3g45730 that encodes a protein of unknown function for cluster 14.

Although Arabidopsis is the most extensively profiled plant for single-cell and single-nucleus transcriptomics, curated marker gene sets and available atlases provided limited coverage with which to define the cell types represented by the clusters recovered here. Experimentally verified literature markers identified the cell types of only ∼50% of clusters (Supplemental Table 1), often with just one high-ranking marker among the top predicted genes. We therefore undertook a comprehensive validation of cluster identities using two complementary experimental strategies. First, we designed a 10x Xenium panel of 465 probes incorporating the top markers from each cluster and from key subclusters (Figure 1A, Supplemental Table 1). Second, we generated 52 promoter–GFP reporter lines to test predicted cell-type enrichment *in planta* (Supplemental Table 2). Together, these approaches confirmed multiple mesophyll clusters (clusters 0, 1, 4, 6, 17, and 24) and enabled experimental re-assignment of three previously unknown clusters (3, 7, and 11) as mesophyll-associated states (Figure 1C Supplemental Figures 1–3; Supplemental Table 2). Examples of enrichment of transcript abundance in mesophyll under control and antimycin A treated conditions is evident in the cross-sections of Arabidopsis leaves for each of these three clusters, *VDAC3* (*Voltage Dependent Anion Channel 3*) (At5g15090) for cluster 3, *BES1* (*BRI1* (*BRASSINOSTEROID INSENSITIVE 1*) *EMS SUPPRESSOR 1)* (At19350) for cluster 7, and *ASN1* (*GLUTAMINE-DEPENDENT ASPARAGINE SYNTHASE 1*) (At3g37340) for cluster 11. Notably, this re-designation was also supported by subclustering analysis, which showed that clusters 3, 7 and 11 each resolve into multiple subclusters whose marker profiles are dominated by mesophyll-associated genes (Supplemental Figure 4; Supplemental Table 2). Cluster 14 was assigned to epidermis (Figure 1C, Supplemental Figures 1–3; Supplemental Table 2), with a gene at locus At3g45730 that encodes a protein of unknown function that shows a strong epidermal enrichment in control conditions, but loses specificity with antimycin A treatment (Figure 1C). Three additional low-abundance unassigned clusters (23, 26, 27) remained unresolved, consistent with limited representation for robust experimental confirmation. Among epidermal and leaf pavement lineages, clusters 5, 8, 9, 10, 12, and 20 were supported by experimental validation, whereas cluster 2 showed mixed epidermal and mesophyll signatures. Specifically an *Expansin*-like gene (At3g45970) and a *myo-inositol oxygenase* (At2g19800) showed mesophyll cell localisation in the 10x Xenium data, whereas *SLAH3* (At5g24030) and a gene encoding a cell wall protein (At4g08950) displayed localisation leaf pavement cells (Supplemental Table 2).

In total, we profiled over 600,000 nuclei from shoots of 14-day-old (stage 1.02) seedlings (Boyes et al., 2001) (Supplemental Table 3), with median UMIs per nucleus ranging from 989 to 3,209 and median genes detected per nucleus from 767 to 1,995 (Supplemental Table 3). This sampling depth is comparable to, or greater than, recent atlas-scale and stress-focused single-cell studies in Arabidopsis and other crops (Guo et al., 2025; Lee et al., 2025; Marand et al., 2021; Sun et al., 2024; Wang et al., 2025b; Wang et al., 2021; Yan et al., 2025; Zhang et al., 2025a). To facilitate community use, we provide a web resource for time-resolved visualization of cell-type–level responses (https://arabidopsismrsdb.com/).

### Mitochondrial dysfunction changes the transcriptional identity of cells

Inhibition of mitochondrial electron transport with antimycin A or myxothiazol caused rapid, time-dependent redistribution of nuclei across clusters relative to time-matched mock controls (Figure 2A; Supplemental Figure 5). Changes were detectable within 1 h, most prominently after antimycin A treatment (Supplemental Figures 5 and 6). The response was maximal at 3 h, when 11 of 28 clusters changed significantly in abundance under both inhibitors, with additional inhibitor-specific shifts observed for a small number of clusters (Figure 2A; Supplemental Figures 5 and 6). After 3 h, the number and magnitude of abundance changes generally did not continue to increase, consistent with the established peak of mitochondrial dysfunction responses around this time in bulk transcript studies (Ng et al., 2013a). We have therefore used of the 3 h time point for detailed cell-state analyses while retaining the full-time course in the supplement (Supplemental Figures 5 and 6).

**Figure 2.**
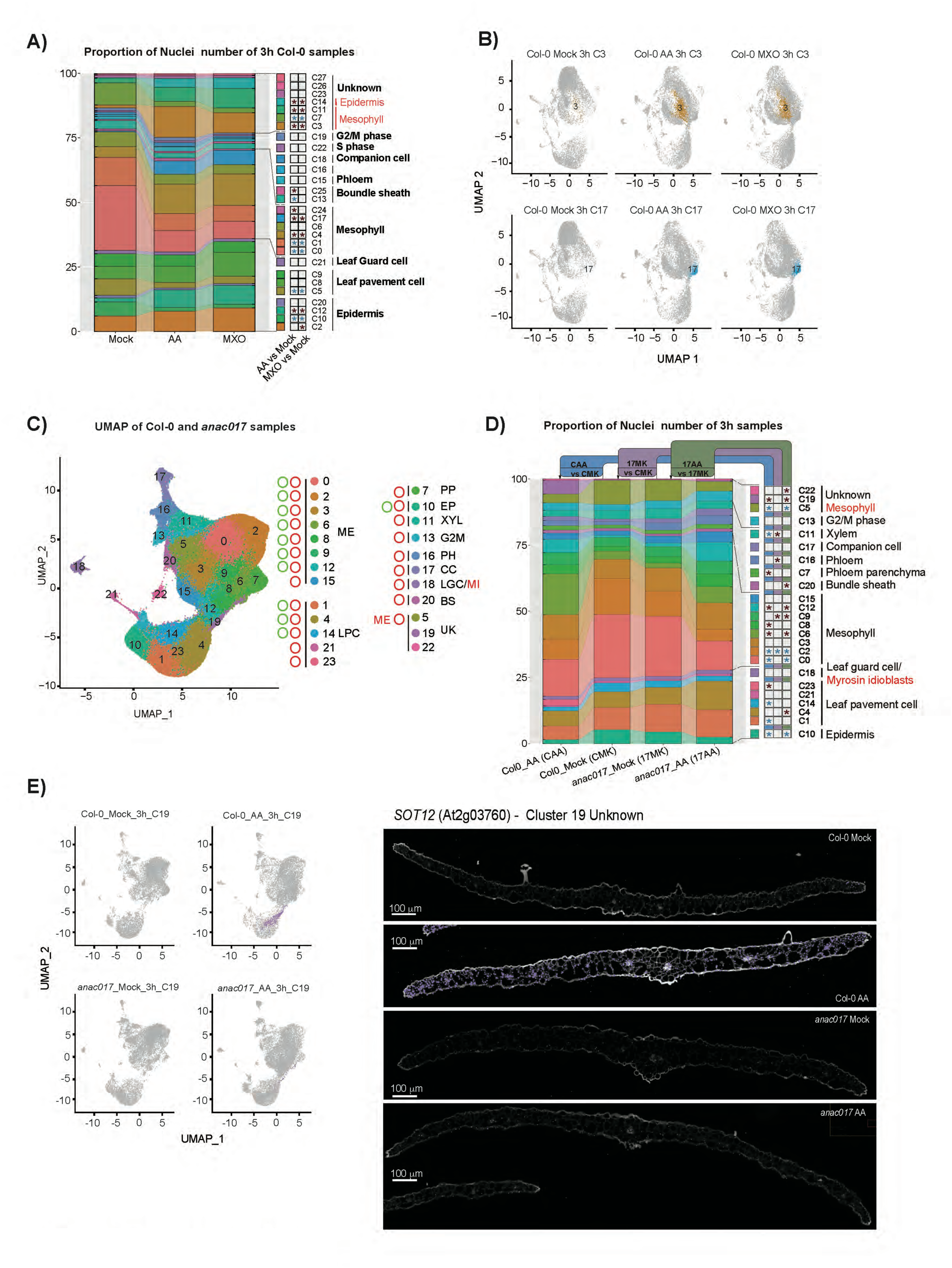
Mitochondrial dysfunction reshapes cell-type composition and reveals ANAC017-dependent stress clusters. **(A)** Proportion of nuclei assigned to each cluster in Col-0 at 3 h under mock, antimycin A (AA) and myxothiazol (MXO) treatment. Stacked bars show the relative contribution of each cell type annotated cluster (colour-coded at right; see Figure 1B, C) to the total nuclei population for each condition, with clusters ordered by cell type. Red asterisks mark clusters with a significant increase and blue asterisks clusters with a significant decrease in nuclei number relative to the corresponding mock control. **(B)** UMAP projections highlighting clusters 3 (C3) and 17 (C17) in Col-0 at 3 h. Each panel shows nuclei from a single condition (mock, AA, MXO), with the cluster of interest coloured (orange for C3, blue for C17) and all other nuclei in grey. Both clusters, annotated as mesophyll, with C3 re-assigned based on marker validation and both expand upon AA and MXO treatment, illustrating treatment-induced shifts in mesophyll cell states. **(C)** Integrated UMAP of snRNA-seq data from Col-0 and *anac017* at 3 h, coloured by cluster identity (0–23). Unsupervised clustering was performed with Seurat v4.1.1, and initial annotation used Single Cell Plant DataBase (Ryu et al., 2021). Major cell types are indicated by coloured labels: mesophyll (ME), epidermis (EP), leaf pavement cells (LPC), bundle sheath (BS), phloem (PH), phloem parenchyma (PP), companion cells (CC), xylem (XYL), leaf guard cells (LGC), G2/M phase (G2M), S phase (SP) and unknown (UK). Cell-type assignments were confirmed using 10x Xenium spatial localisation (red circles) and promoter–GFP lines (green circles); clusters originally annotated as unknown are retained as UK for reproducibility, with experimentally supported identities indicated in red. **(D)** Proportion of nuclei in each cluster for Col-0 and *anac017* at 3 h under mock or AA treatment. Stacked bars summarise three comparisons: (i) Col-0 AA versus Col-0 mock (blue, CAA vs CMK), (ii) *anac017* mock versus Col-0 mock (purple, 17MK vs CMK) and (iii) *anac017* AA versus *anac017* mock (green, 17AA vs 17MK). Red and blue asterisks indicate significant increases and decreases in nuclei number, respectively. Corresponding analyses at 1 and 6 h are shown in Supplemental Figure 7. **(E)** UMAP projections highlighting cluster 19 (C19) in Col-0 and *anac017* at 3 h. Each panel shows C19 nuclei (purple) overlaid on all other nuclei (grey) for mock and AA treatments. C19 expands strongly in Col-0 but shows a much smaller increase in *anac017*, illustrating an ANAC017-dependent mesophyll stress state. *SOT12* (*SULFOTRANSFERASE 1*) (At2g03760) is an example of a gene from this cluster that is induced with antimycin A in an ANAC017 dependent manner.

The strongest abundance changes at 3 h occurred within mesophyll-associated states. Mesophyll clusters 0 and 1 declined significantly under both inhibitors (Figure 2A; Supplemental Figures 5–6). These reductions were accompanied by increases in other mesophyll-designated clusters, including cluster 4 and the re-assigned mesophyll clusters 3 and 11, indicating a shift in transcriptional identity within the mesophyll lineage rather than a uniform response across all mesophyll states (Figure 2A; Figure 1B; Supplemental Figures 5–6; Supplemental Table 3).

Epidermal and leaf pavement lineages also showed divergent behaviours across related clusters. Leaf pavement cluster□5 decreased significantly at 3□h and 12□h and also showed a nonlZIsignificant decrease at 6□h, while an epidermal cluster 10 decreased at 3 h and 6 h (Figure 2A; Supplemental Figures 5 and 6). In contrast, epidermal clusters 12 and 14 increased at 3 h and remained elevated at 6 h and 12 h (Figure 2A; Supplemental Figures 5 and 6). Cell cycle–associated clusters (G2/M (cluster 19) and S phase (cluster 22)) displayed a transient shift at 1 h, with little evidence of sustained abundance changes thereafter (Figure 2A; Supplemental Figures 5 and 6). UMAP visualizations illustrate these shifts at the level of individual clusters. At 3 h, clusters 3 (re-designated mesophyll) and 17 (mesophyll) expanded strongly under both inhibitors (Figure 2B), and similar expansion was evident at later time points for cluster 3 and for the epidermal cluster 14 (Supplemental Figure 6). Collectively, these patterns show that mitochondrial dysfunction does not elicit a uniform response within a given cell type. Instead, it redistributes nuclei among discrete transcriptional states, revealing heterogeneity both between and within canonical cell identities. Notably, clusters that expanded most strongly after treatment were not exclusive to the stressed samples but contained a small number of nuclei in untreated controls. This suggests that a subset of cells already occupies, or readily accesses, a stress-associated transcriptional programme in the basal state, consistent with the presence of putative primed cells that can rapidly increase in prevalence upon mitochondrial perturbation.

### ANAC017 reshapes cell type specific responses to mitochondrial dysfunction

The NAC transcription factor ANAC017 is a central regulator of mitochondrial stress signalling that controls both positive and negative downstream regulators (He et al., 2022; Zhu et al., 2023). To determine how ANAC017 influences cellular responses to mitochondrial inhibition, we compared Col-0 and *anac017* mutant seedlings treated with antimycin A for 1, 3 and 6 h using snRNA-seq (Figure 2 C, D; Supplemental Figure 7). Joint analysis of both genotypes resolved 24 clusters spanning eight mesophyll clusters, five leaf pavement cell clusters, and single clusters corresponding to phloem parenchyma, companion cells, guard cells, bundle sheath, phloem, epidermis, xylem, and G2/M phase, together with three initially undefined clusters (Figure 2C). Spatial transcriptomics and promoter–GFP validation supported re-assignment of two of these undefined clusters (C5 and C19) as mesophyll-associated states (Figure 2C, D; Supplemental Table 4). Cluster 18 was predicted to be leaf guard cell and has been additionally annotated as Myrosin idioblasts as they share marker genes (Shirakawa et al., 2025).

Under mock conditions, cluster composition was broadly similar between genotypes, with a limited number of differences at 3 h, including reduced representation of one mesophyll cluster (C2) and increased representation of another (C9) in *anac017*, together with modest shifts in phloem (C16) and xylem (C11) (Figure 2D; Supplemental Table 5). No significant genotype-dependent differences were detected at 1 h in mock samples, and only a small number emerged at 6 h (Supplemental Figure 7; Supplemental Table 5).

Antimycin A treatment induced both shared and genotype-dependent redistribution across clusters. At 3 h, both genotypes showed decreased abundance of mesophyll-associated clusters C0, C2 and C5 and the epidermal cluster C10, together with increased abundance of mesophyll-associated clusters C6 and C12 (Figure 2D; Supplemental Figure 7; Supplemental Table 5). Beyond these shared shifts, several treatment-responsive clusters diverged strongly between genotypes. For example, a leaf pavement cluster (C1) decreased in Col-0 but not in *anac017*, while another leaf pavement cluster (C4) increased in *anac017* across multiple time points but showed little change or a decrease in Col-0 (Figure 2D; Supplemental Figure 7; Supplemental Table 5). Differences were also evident in mesophyll cluster C8 across the time course, and in the transient induction of the G2/M cluster (C13) at 1 h in Col-0, which was absent in *anac017* (Supplemental Figure 7; Supplemental Table 5). The strongest ANAC017 dependence was evident in two stress-associated clusters, C23 (leaf pavement) and C19 (unknown), which expanded sharply in Col-0 but showed minimal or strongly attenuated expansion in *anac017*, despite being present at comparable low levels in untreated samples (Figure 2D, E; Supplemental Figure 7; Supplemental Table 5). An examples of a gene for this cluster is *SOT12* (*SULFOTRANSFERASE 1*) (At2g03760) that has been shown to be a direct target of ANAC017 (Zhu et al., 2023) (Figure 2E).

Across the time course, the overall pattern of remodeling also differed. By 6 h, a larger number of clusters changed significantly in *anac017* than in Col-0, with several mesophyll and leaf pavement clusters showing higher representation in the mutant (Supplemental Figure 7; Supplemental Table 5). This is consistent with a delayed or compensatory progression of transcriptional reprogramming when a central regulator is absent, such that the wild type reaches a stabilized state earlier while the mutant continues to redistribute across stress-associated cell states.

### Transcriptional responses to mitochondrial inhibition are cell-state specific and ANAC017-dependent

We next examined how individual genes and clusters respond to antimycin A and myxothiazol treatment. Firstly, we performed independent reclustering analysis for each time point for Col-0 samples. This analysis identified 21, 24, 22, and 24 distinct cell clusters in the 1, 3, 6, and 12 h samples, respectively (Figure 3A, Supplemental Table 3). In Col-0, classical markers of mitochondrial dysfunction such as *RAV2* (*Related to ABI3/VP1 2*) and *AOX1a* were rapidly induced within 1 h, whereas genes including *UPOX* (*Up-regulated by oxidative stress*, AT2G21640) and *LETm* (*Leucine Zipper-EF-hand-containing Transmembrane protein*, AT3G59820) showed slower induction (Figure 3A, panel I). By 3 h, genes encoding LEA5 (Late embryogenesis abundant like protein 5, AT4G02380), GSTU9 (Glutathione S-transferase tau 9, AT5G62480) and VDAC3 (Voltage-dependent anion channel 3, AT5G15090) were broadly induced across nearly all clusters (Figure 3A, panel II). These genes are not usually included in canonical mitochondrial stress marker sets, likely because earlier microarray platforms lacked specific probes. In contrast, genes such as *NDB2* (*NAD(P)H dehydrogenase B2*, AT4G05020) and *ACO1* (*Aconitase 1*, AT4G35830) were induced in only about half of the clusters (Figure 3A, panels I and II), indicating more restricted, cell type dependent responses. Genes encoding components of the mitochondrial protein import machinery showed later induction (Figure 3A, panels III and IV), consistent with a response that peaks between 3 and 6 h and then relaxes toward a new steady state.

**Figure 3.**
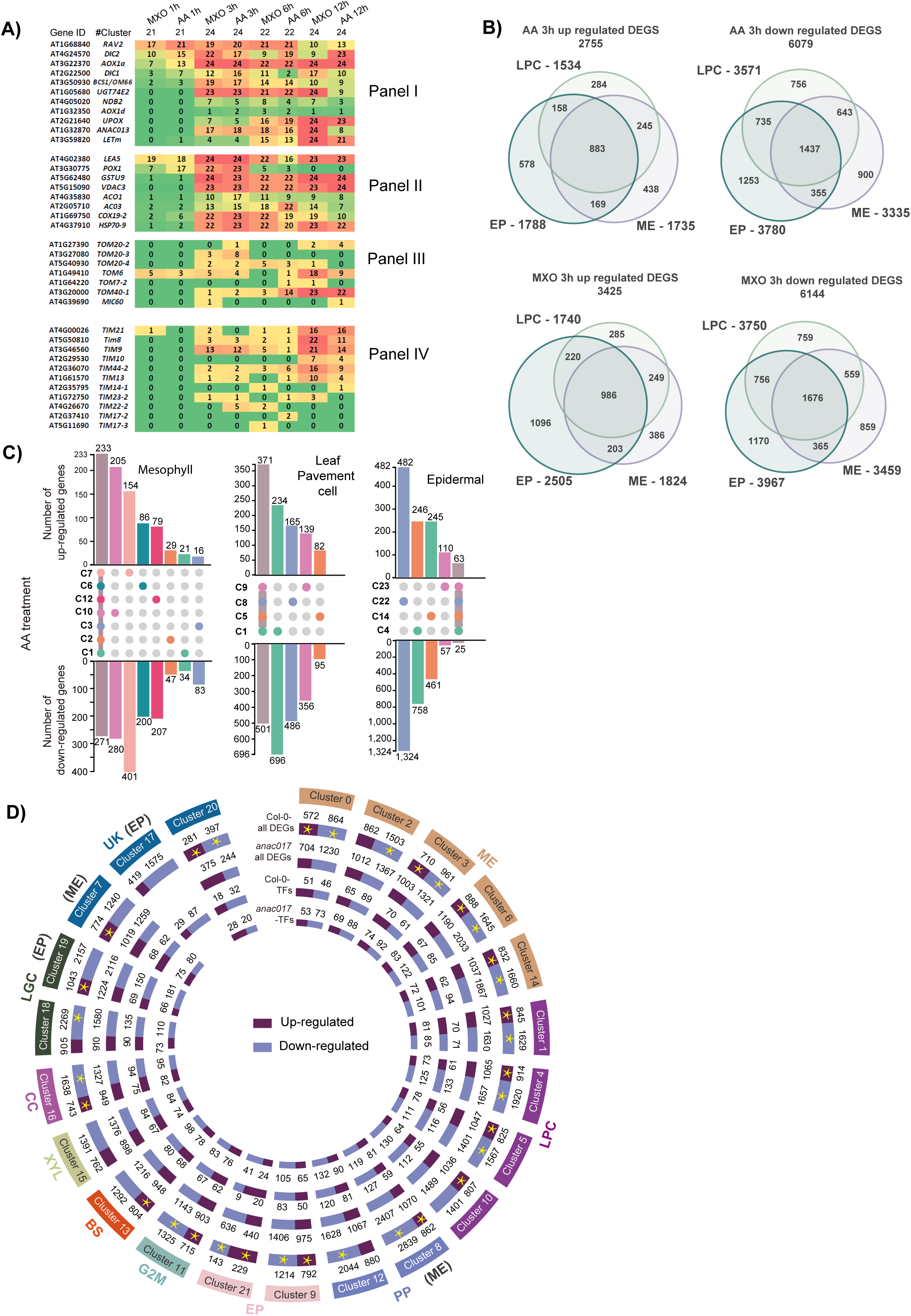
A gene-level view of the mitochondrial stress response at cellular resolution. **(A)** Response of genes encoding mitochondrial proteins to antimycin A (AA) and myxothiazol (MXO) in Col-0 at 1, 3, 6 and 12 h after treatment. The heatmap shows, for each gene (rows), the number of clusters in which it is differentially expressed at each time point (columns). Panels I - nuclear genes encoding mitochondrial proteins widely used as markers of mitochondrial dysfunction. II) nuclear genes encoding mitochondrial proteins identified as broadly responsive but not traditionally used as markers of mitochondrial dysfunction, III and IV) genes encoding components of the mitochondrial protein import machinery, that differ in both response kinetics and the extent of cluster coverage. **(B)** Numbers of genes that increase or decrease in transcript abundance after 3 h of AA or MXO treatment and their overlap between epidermal (EP), leaf pavement cell (LPC) and mesophyll (ME) clusters. Venn diagrams separate upregulated and downregulated differentially expressed genes (DEGs) for each treatment. **(C)** Common and unique DEGs within epidermal, leaf pavement cell and mesophyll clusters after 3 h AA treatment. Bar plots indicate the number of up- and downregulated genes per cluster, illustrating that many DEGs are cluster specific rather than shared across all clusters of a given cell type. **(D)** Total number of DEGs and DEG-encoded transcription factors per cluster in Col-0 and *anac017* after 3 h AA treatment. Concentric rings represent Col-0 and *anac017*, with segments corresponding to individual clusters whose numbers and cell-type identities are indicated in the outer ring. Yellow asterisks mark clusters in which the proportion of genes that change in transcript abundance differs significantly between Col-0 and *anac017* (p < 0.001, based on all detected genes). The number of transcription factor DEGs per cluster is shown but the numbers are insufficient for a proportionality test. See Supplemental Figure 11 for all time points. Major cell types are indicated by coloured labels: mesophyll (ME), leaf pavement cells (LPC), epidermis (EP), phloem parenchyma (PP), G2/M phase (G2M), bundle sheath (BS), xylem (XYL), companion cells (CC), leaf guard cells (LGC) and unknown (UK). Clusters re-designated using experimental based markers are indicated in brackets.

At the cluster level, the number of differentially expressed genes (DEGs) per cluster ranged from as low as 50 to several thousand, but this needs to be cautiously interpreted due to the differences in cell number per cluster that also varied from thousands to as little as a few hundred (Supplemental Figure 9; Supplemental Tables 3, 6, 7). Overlap analysis between epidermal, leaf pavement and mesophyll clusters at 3 h showed that each cell type harboured several hundred to more than a thousand unique DEGs (Figure 3B). Within each cell type, shared DEGs in different clusters represented only a small fraction of the total, with many cluster-specific changes. For example, an epidermal cluster (C22) exhibited more than 1,000 unique DEGs (Figure 3C). Similar patterns were observed at 1, 6 and 12 h, with differences primarily reflecting response kinetics (Supplemental Figure 9 A-D). At 1 h, epidermal and leaf pavement clusters showed less overlap with mesophyll, which may relate to topical application of the inhibitors (Supplemental Figure 9 A).

Comparing DEGs between Col-0 and *anac017* after antimycin A treatment highlighted the regulatory role of ANAC017. At 1 h, most clusters had many more DEGs in Col-0 than in *anac017*, with the exception of clusters 18 (mesophyll) and 20 (leaf pavement cell), which contained relatively few nuclei (∼600 in total) (Supplemental Figure 10A). By 3 h, DEG numbers became more similar between the genotypes and were higher in *anac017* for some clusters (Supplemental Figure 10B). By 6 h, the pattern had reversed: most clusters showed many more DEGs in *anac017* than in Col-0 (Supplemental Figure 10C). This pattern is consistent with previous bulk RNA sequencing studies showing that the response to antimycin A treatment in Col-0 peaks at approximately 3 h, at which point a new transcriptional homeostasis is established (He et al., 2022; Ng et al., 2013b). By contrast, in the absence of ANAC017, this stabilization is delayed, and transcriptomic changes continue to accumulate over time, likely reflecting slower or incomplete engagement of compensatory regulatory pathways.

We quantified these differences by testing whether the proportion of significantly changed genes differed between genotypes in each cluster. Using a threshold of p < 0.001, most clusters showed a significantly different number of DEGs between Col-0 and *anac017* at 3 h (Figure 3D). A similar trend was observed at 1 and 6 h, where most clusters again differed significantly between genotypes (Supplemental Figure 11; Supplemental Table Statistics). Overlap analysis of DEG identities at 3 h showed that many genes were specific to either Col-0 or *anac017* in most clusters (Supplemental Figure 10). Thus, the response to antimycin A differs between Col-0 and *anac017* in the kinetics of induction, the fraction of genes that change in each cluster and the identities of those genes, consistent with ANAC017 acting as a high-level regulator of the mitochondrial stress response.

### Spatial transcriptomics reveals patterned and selective activation of mitochondrial stress programs

A range of spatial transcriptomics platforms has been applied in plants, spanning low-throughput PHYTOMap (Nobori et al., 2023) and medium-throughput MERSCOPE (Lee et al., 2025), through to higher-throughput 10x Visium and Stereo-seq, which provide sub-tissue but not cellular resolution (Peirats-Llobet et al., 2023; Zhang et al., 2025a). Visium typically operates at ∼50 μm resolution, whereas Stereo-seq resolution depends on binning and can range from ∼2 μm to ∼100 μm across bin sizes of 3 – 140 (Chen et al., 2022), with most plant applications using approximately 20–100 μm bins (Guo et al., 2025; Xia et al., 2022). We therefore chose to use 10x Xenium to enable single cell localization at medium throughput using a custom 465-probe panel. We analysed 174 sections in total, with 19 to 25 sections per time point at 3 h and 6 h for each genotype (Col-0 and *anac017*), generated across two independent experiments (Supplemental Figure 3). To avoid selection bias, we present the same representative leaf cross-section for each gene across treatments. All sections and probe signals are available via Zenodo (https://zenodo.org – <u>see data availability</u>), together with instructions for exploration in 10x Xenium Explorer, and examples of cell type markers in all sections are also available in Supplemental Figure 3).

To provide a matched reference for spatial analyses, we clustered the 3 h snRNA-seq dataset from Col-0 and *anac017*, which resolved 22 clusters. Two clusters showed mixed marker signatures and could not be assigned confidently, and one cluster contained very few nuclei (Figure 4A; Supplemental Table 4). Spatial RNA profiling at 3 h resolved 19 clusters that aligned with the major tissue domains of the leaf, including multiple epidermal and mesophyll clusters, a vascular cluster, guard cells, and an S-phase cluster (Figure 4A).

**Figure 4.**
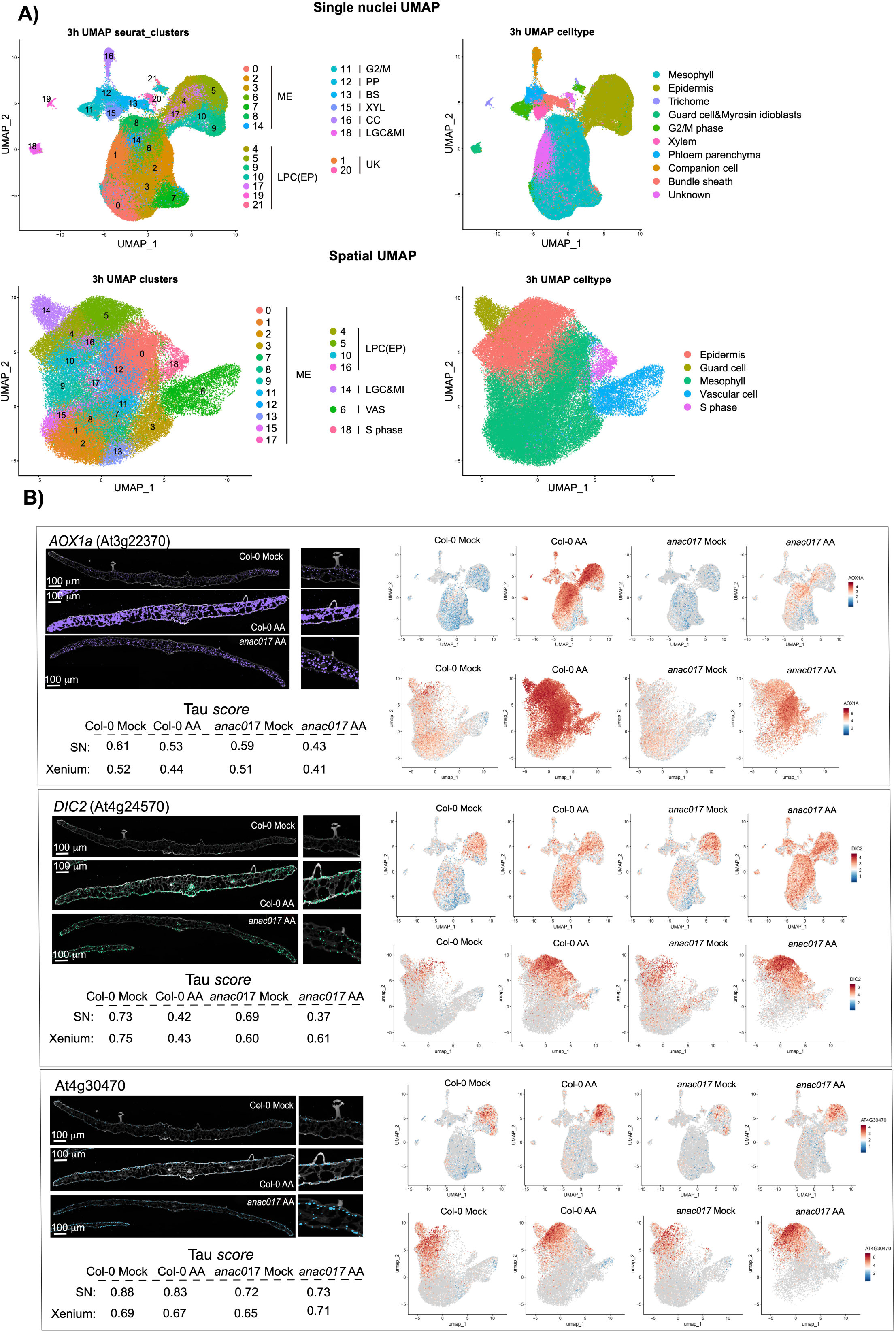
Spatial context of mitochondrial dysfunction. **(A)** UMAPs of single-nucleus RNA-seq (top, snRNA-seq) and spatial RNA-seq (bottom, spRNA-seq) from Col-0 and *anac017* seedlings after mock or antimycin A (AA) treatment. For each modality, the left panel shows individual clusters (snRNA-seq: 22 clusters, 0–21; spRNA-seq: 19 clusters, 0–18), and the right panel groups clusters into major cell types (epidermis, mesophyll, vascular, leaf guard cell, S-phase and unknown). Cell types were annotated using published single-cell databases (black labels), 10x Xenium marker expression (red) and promoter–GFP lines (green) (Supplemental Figure 8). Predicted and experimentally confirmed markers are listed in Supplemental Table 4. Representative tissue sections for marker genes are shown in Supplemental Figure 1 and 3. **(B)** Spatial expression patterns of genes that report perturbation of mitochondrial function. For each gene, the left images show 10x Xenium tissue sections for Col-0 control (Col-0 mock) and AA-treated samples at 3 h (one representative section of 10-12 per time point; 20-24 sections total per condition). The right panels show UMAP feature plots for snRNA-seq (top row) and spatial RNA-seq (bottom row) for Col-0 mock, Col-0 AA, *anac017* mock and *anac017* AA, coloured by scaled expression. Tau scores beneath each panel summarise expression specificity (1 = highly specific, 0 = broad). *AOX1a* = *Alternative oxidase 1a*, *DIC2* = *Dicarboxylate carrier 2*, SN = single nuclei.

Spatial mapping of established mitochondrial dysfunction marker genes (De Clercq et al., 2013; Schwarzländer et al., 2012; Van Aken et al., 2009) revealed distinct classes of response. *AOX1a* showed broad induction across essentially all clusters and tissue regions after antimycin A treatment, consistent with its role as a widely responsive marker of mitochondrial perturbation (Figure 4B; Supplemental Table 7). This induction was strongly attenuated in *anac017*, matching the reduced responsiveness observed in the single-nucleus dataset (Figure 4B; Supplemental Table 7). In contrast, *DIC2* (*Dicarboxylate Carrier 2*, At4g24570) exhibited an epidermis-enriched pattern in control tissue but expanded into mesophyll and regions adjacent to vascular bundles after treatment (Figure 4B; Supplemental Table 7). Notably, this expansion was selective, with limited induction in several mesophyll-associated spatial clusters, indicating that mitochondrial dysfunction does not uniformly propagate even within a single tissue layer (Figure 4B). Similarly, *Glutathione S-transferase F6* (*GSTF6*, At1g02930) and *Glutathione S-transferase F7* (*GSTF7*, At1g02920), which have experimental and proteomic support for mitochondrial localisation among other reported localisations (Fuchs et al., 2020; Hooper et al., 2017), showed epidermal and trichome-associated signals under control conditions, followed by induction in epidermis and subsets of mesophyll rather than uniform activation across all clusters (Supplemental Figure 12; Supplemental Table 7). A third pattern was illustrated by At4g30470, which maintained epidermal specificity before and after treatment, indicating that induction can occur without loss of spatial restriction (Figure 4B; Supplemental Table 7).

The spatial data shows that mitochondrial inhibition generates a structured response landscape. Some transcripts are induced broadly across cell types and tissue domains, whereas others expand from pre-existing tissue-enriched patterns into selected clusters or remain spatially restricted despite strong induction.

### Transcription factor spatial specificity is remodelled by mitochondrial dysfunction

To elucidate how mitochondrial dysfunction reshapes regulatory programs across cell types, we quantified cell type-enhanced expression for 1119 genes encoding transcription factors using a *tau* specificity threshold of ≥0.85, where values closer to 1 indicate strong cell type specificity and values closer to 0 indicate broad expression. Across the four datasets (Col-0 mock, Col-0 antimycin A, *anac017* mock, *anac017* antimycin A) representing the 3□h time point, 404 transcription factor genes exceeded this threshold in at least one condition (Figure 5A; Supplemental Table 8). We then used the *tau* score for each condition (represented by coloured dots in Figure 5A) together with the standard deviation of *tau* across conditions to distinguish transcription factors whose expression remained consistently cell type–enhanced from those whose specificity was remodeled by treatment or by loss of ANAC017 (Figure 5A).

**Figure 5.**
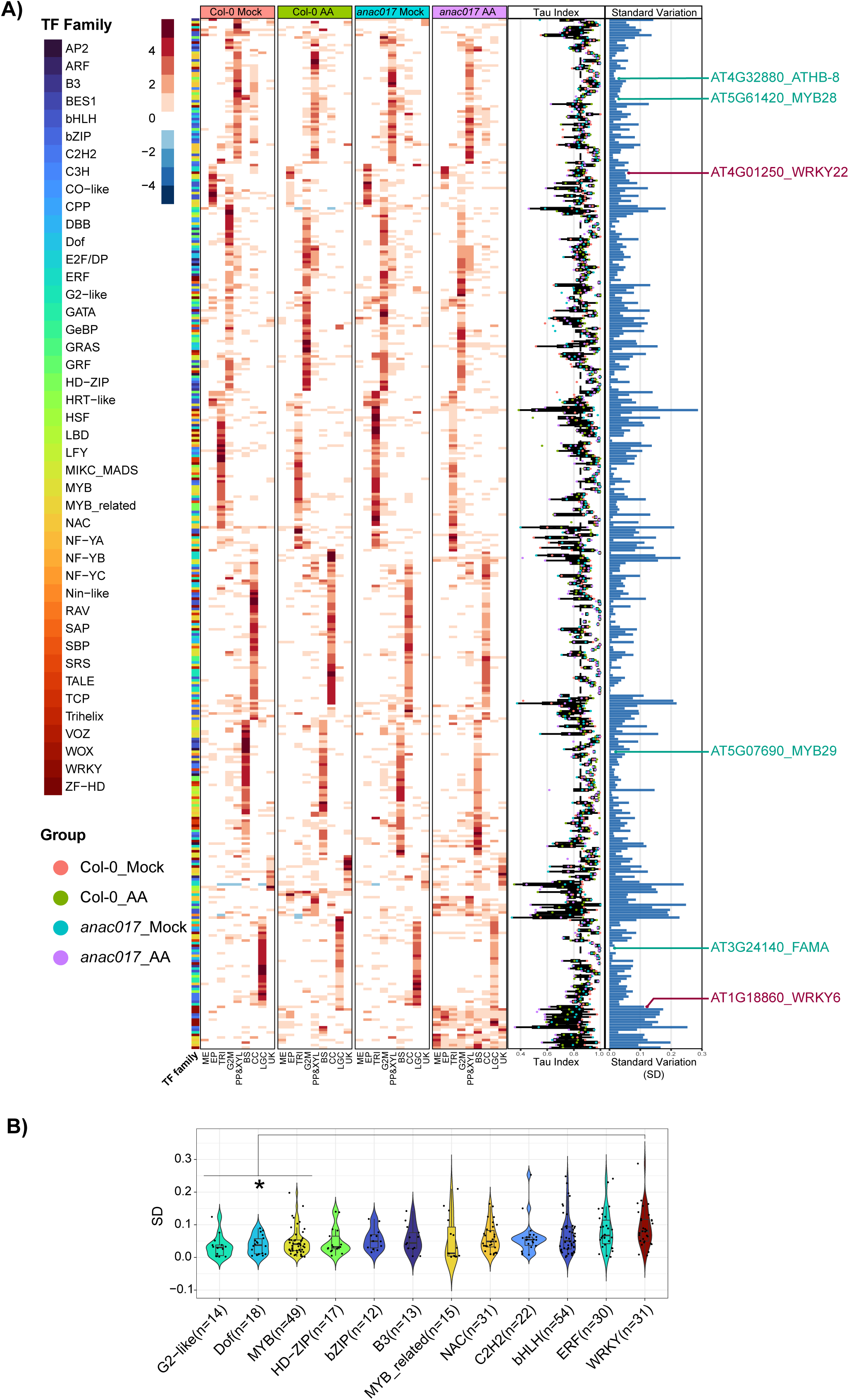
Cell type–specific and stress-responsive transcription factor expression. **(A)** Cell type specificity of transcription factor (TF) genes at 3 h. A total of 1,119 TF-encoding genes were assessed using the *tau* index, where 1 indicates expression confined to a single cluster and 0 indicates broad expression. Genes with *tau* ≥ 0.85 in at least one condition (404 TFs) are shown. The heatmap displays tau scores and expression levels across clusters for Col-0 mock, Col-0 antimycin A (AA), *anac017* mock and *anac017* AA. TF families are indicated by the colour bar on the left. The bar plot to the right shows the standard deviation of *tau* across the four conditions for each TF, highlighting factors whose cell type specificity shifts most strongly with AA treatment or loss of ANAC017. Selected TFs examined in detail in later figures are labelled. mesophyll (ME), epidermis (EP), trichome (TRI), bundle sheath (BS), phloem parenchyma (PP), xylem (XYL), companion cells (CC), leaf guard cells (LGC), G2/M phase (G2M) and unknown (UK). **B)** *Tau* index standard deviation (SD) of transcription factor-coding genes across different transcription factor families. Each dot represents the *tau* index SD of an individual transcription factor-coding gene in (A). The SD reflects variation in the *tau* index across the four groups: Col-0 mock, Col-0 AA, *anac017* mock, and *anac017* AA. Only transcription factor families containing more than 10 gene members are displayed. Statistical comparisons of Tau index SD distributions among transcription factor families were performed using a two-tailed Student’s t-test (*, p < 0.05).

Several transcription factors retained strong and stable cell type enhancement, indicating that mitochondrial dysfunction can preserve developmental or lineage-associated regulatory boundaries. FAMA (basic helix-loop-helix (bHLH) At3g24140), a key stomatal regulator (Ohashi-Ito and Bergmann, 2006; Smit and Bergmann, 2023), remained enriched in stomatal lineages, and this restriction was maintained after antimycin A treatment in both Col-0 and *anac017* (Figure 5A; Supplemental Figure 13). Similarly, HOMEOBOX PROTEIN 8 (HB-8; At4g32880) remained vascular enriched (Zhang et al., 2025b), and MYB DOMAIN PROTEIN 28 (MYB28; At5g61420) and MYB DOMAIN PROTEIN 29 (MYB29; At5g07690) maintained vascular bundle localization after treatment (Supplemental Figure 13). The persistence of vascular restriction for MYB29, previously linked to repression of the mitochondrial stress response (Zhang et al., 2017b), highlights that negative regulatory inputs can remain spatially confined even when canonical mitochondrial dysfunction markers are broadly induced.

In contrast, other transcription factor families showed substantial condition-dependent shifts in specificity. The WRKY family displayed the greatest variation in *tau* across the four datasets (Figure 5B), consistent with its established involvement in mitochondrial stress signalling (Evans et al., 2024; Van Aken et al., 2013; Zhang et al., 2017a). Spatial profiling illustrated distinct modes of remodeling among induced WRKY transcription factors (Figure 6). *WRKY6* gained mesophyll expression after antimycin A treatment while retaining epidermal enrichment that was biased toward the lower epidermis, and induction remained restricted to a subset of epidermal and mesophyll clusters. *WRKY11* shifted from epidermal-enhanced expression under control conditions to broader expression including mesophyll after treatment, while still remaining restricted within mesophyll to specific clusters and showing limited dependence on ANAC017. By contrast, WRKY22 increased in abundance but largely preserved epidermal-enhanced expression.

**Figure 6.**
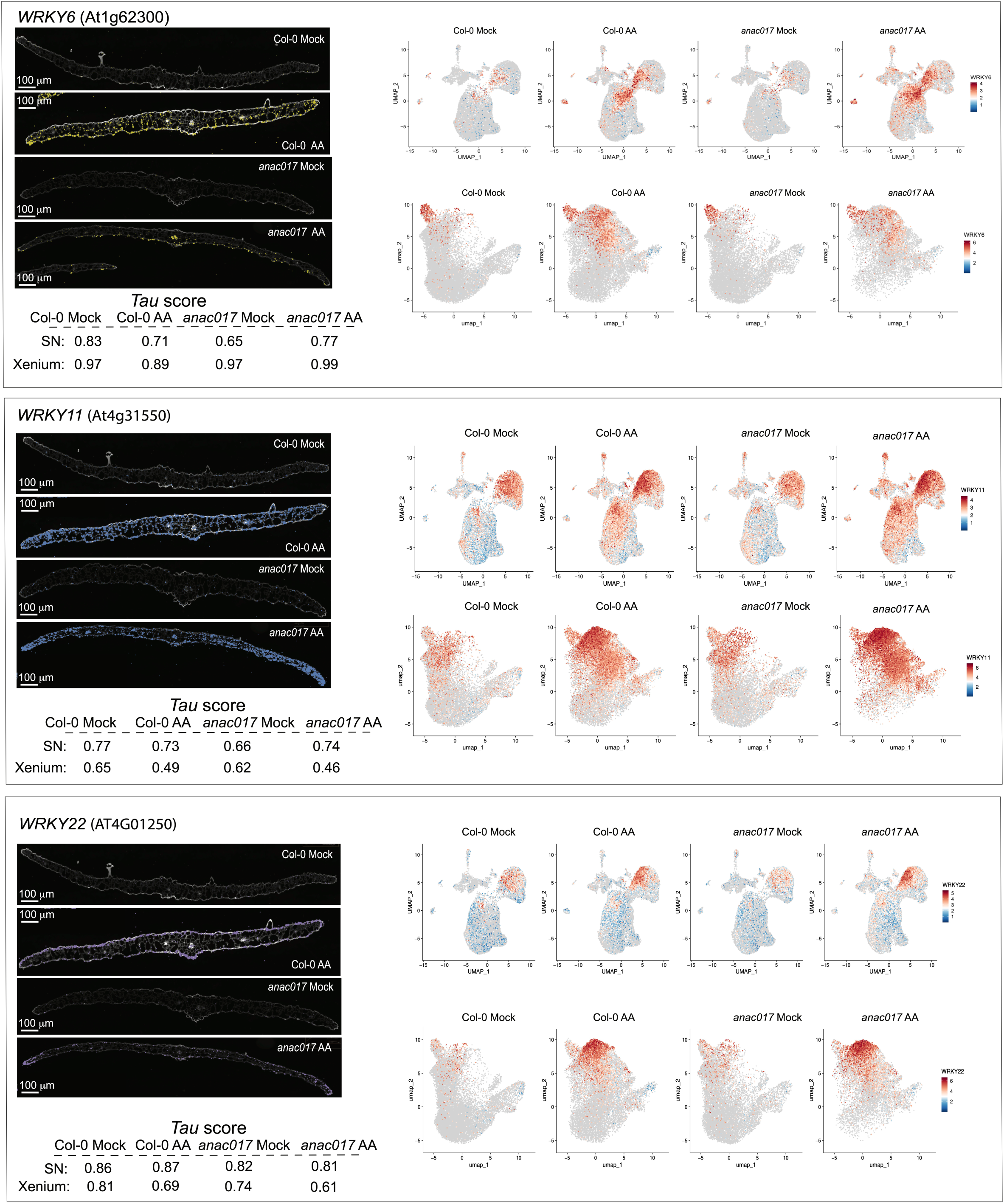
Spatial and cell type–specific induction of WRKY transcription factors by mitochondrial dysfunction. **(A)** *WRKY6* (At1g62300). Left, 10x Xenium spatial transcriptomics images of leaf cross sections from Col-0 control (Col-0 mock), Col-0 antimycin A (Col-0 AA), *anac017* mock and *anac017* AA, with *WRKY6* transcripts pseudo-coloured and overlaid on Fluorescent brightener 28 (FB28) stained leaf sections that stains cell walls. Right, UMAP feature plots from single-nucleus RNA-seq (SN; top row) and spatial RNA-seq (10x Xenium; bottom row) for the same four conditions, coloured by scaled expression. Under control conditions *WRKY6* is largely restricted to epidermal and guard cells, and AA treatment induces expression in a subset of mesophyll clusters with a bias toward the abaxial epidermis. *Tau* scores beneath the panels summarise cell type specificity (1 = highly specific, 0 = broad) for Col-0 mock, Col-0 AA, *anac017* mock and *anac017* AA in SN and 10x Xenium datasets. **(B)** *WRKY11* (At4g31550). Spatial sections and UMAP feature plots as in **(A)**. *WRKY11* shows epidermis-enriched expression under control conditions. AA treatment leads to strong induction in mesophyll clusters in both Col-0 and *anac017*, although expression remains confined to a subset of mesophyll clusters, indicating a selective expansion rather than a uniform response. *Tau* scores indicate a marked shift in cell type specificity upon treatment. **(C)** *WRKY22* (At4g01250). Spatial sections and UMAP feature plots as in (A). WRKY22 displays pronounced epidermal-enriched expression in control leaves and largely maintains this spatial pattern after AA treatment, with only limited induction in internal tissues. *Tau* scores show high specificity under all conditions.

Together, these patterns show that mitochondrial dysfunction can either preserve, relax, or reconfigure transcription factor spatial restriction, generating regulator landscapes that differ not only between cell types but also between clusters within the same cell type.

### Mitochondrial dysfunction reveals spatial domains and leaf polarity in stress regulation

MYB96 (MYB domain protein 96, At5g62470), which promotes cuticular wax synthesis and mediates ABA responses (Lee and Seo, 2019; Lee et al., 2016), retained strong epidermal-enriched expression under antimycin A treatment (Figure 7). *ERF19* (*Ethylene response factor 19*, At1g22810) was induced primarily in an epidermal cluster in snRNA-seq and in the corresponding epidermal cluster in spatial data, with additional signal in some S-phase cells (Figure 7). Leaf cross sections showed that ERF19 induction was stronger in the adaxial (upper) epidermis than in the abaxial (lower) epidermis, except near the central vein (Figure 7). YAB5 (YABBY5, At2g26580), a YABBY family transcription factor with abaxial polarity in leaves (Kojima et al., 2011; Siegfried et al., 1999), maintained abaxial-enriched expression under antimycin A treatment and served as a robust marker for the lower leaf surface (Figure 7). *MYB15* (At3g23250), which regulates lignification and basal immunity (Chezem et al., 2017), and *MYB51* (*MYB domain protein 51*, At1g18570), which controls indolic glucosinolate synthesis (Gigolashvili et al., 2007), both showed abaxially biased induction under antimycin A treatment in UMAP projections and spatial sections (Figure 7). *MSRB6* (At4g04840), encoding methionine sulfoxide reductase B6 and induced by photooxidative stress (Vieira Dos Santos et al., 2005), showed exclusive adaxial epidermal expression under control conditions. After antimycin A treatment, expression extended to both adaxial and abaxial epidermis (Figure 7). *LTP1* (At2g38540), a lipid transfer protein involved in ethylene signalling (Wang et al., 2016), was expressed in epidermis, guard cells and some mesophyll under control conditions, with a bias towards the adaxial epidermis in control conditions. Antimycin A treatment shifted *LTP1* to stronger expression in the abaxial epidermis (Figure 7). *ASB1* (At1g24807), encoding anthranilate synthase subunit beta and controlling a rate-limiting step in tryptophan and hence auxin biosynthesis (Abu-Zaitoon et al., 2023; Stepanova et al., 2005), was expressed in mesophyll and some epidermal cells in control samples. With antimycin A treatment, *ASB1* was more strongly induced on the abaxial side of the leaf (Figure 7), in contrast to the adaxial-biased induction of *ERF19*.

**Figure 7.**
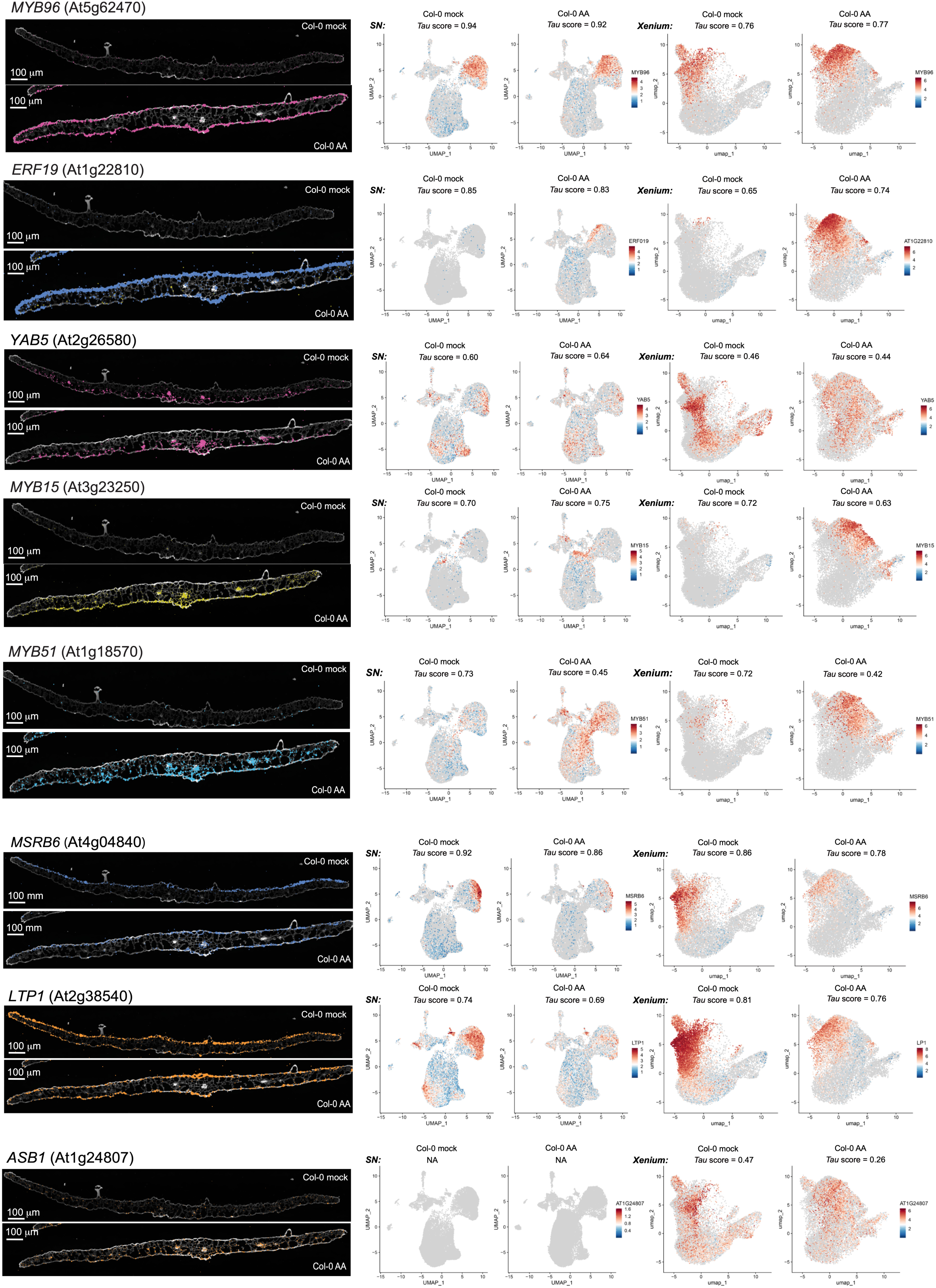
Treatment with antimycin A changes the polarity of gene expression. **(A–H)** Spatial transcriptomics and single nucleus RNA sequencing of polarity-biased regulators in Col-0 under control (mock) and antimycin A (AA) treatment. For each gene, the left panels show 10x Xenium spatial transcriptomics images of leaf cross sections from Col-0 mock and 3 h AA-treated seedlings, with transcripts overlaid on FB28 leaf sections that stains cell walls. The right panels show uniform manifold approximation and projection (UMAP) feature plots for snRNA-seq (SN) and spatial RNA sequencing (10x Xenium) for Col-0 mock and Col-0 AA, with colour indicating scaled expression. *Tau* values above each panel report cell type specificity (1 = highly specific, 0 = broad). Cluster and cell-type identities are as defined in Figure 5A. (A) MYB DOMAIN PROTEIN 96 (MYB96, At5g62470). (B) ETHYLENE RESPONSE FACTOR 19 (ERF19, At1g22810). (C) YABBY5 (YAB5, At2g26580). (D) MYB DOMAIN PROTEIN 15 (MYB15, At3g23250). (E) MYB DOMAIN PROTEIN 51 (MYB51, At1g18570). (F) METHIONINE SULFOXIDE REDUCTASE B6 (MSRB6, At4g04840). (G) LIPID TRANSFER PROTEIN 1 (LTP1, At2g38540). (H) ANTHRANILATE SYNTHASE BETA SUBUNIT 1 (ASB1, At1g24807).

Finally, we examined other stress-related regulators that displayed distinctive spatial or polarity patterns upon antimycin A treatment. HIS4 (Histone H4, At2g28740), encoding histone H4, was expressed at low levels across most cell types under control conditions. After antimycin A treatment, HIS4 expression became more spatially restricted and co-localised with the S-phase marker HTA6 (Histone H2A 6, At5g59870, Figure 8), suggesting a tighter association with proliferative cells during mitochondrial stress (Figure 8; Supplemental Figure 1).

**Figure 8.**
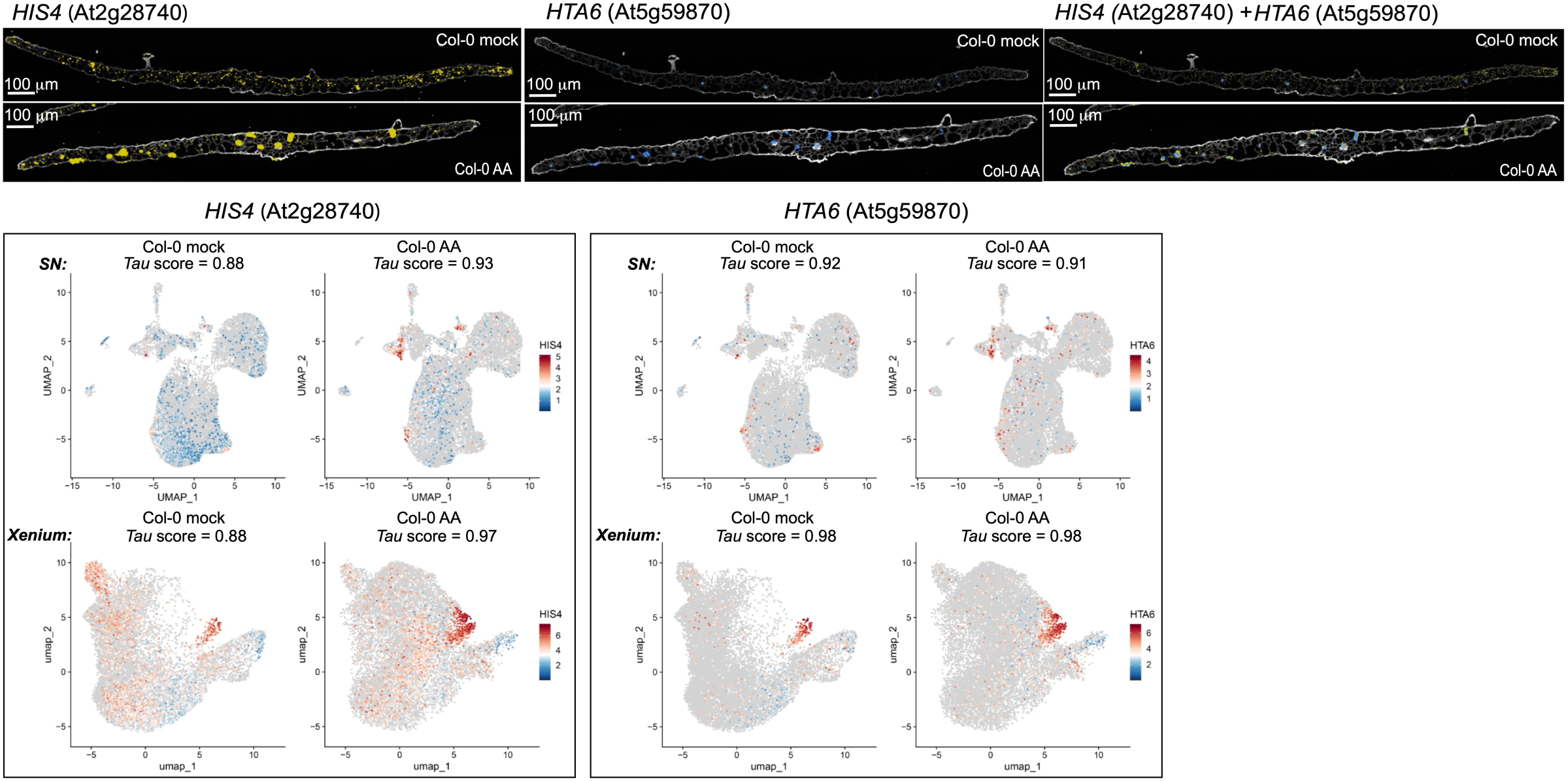
Spatially resolved induction of chromatin and nutrient signalling regulators during mitochondrial stress. Spatial localisation of HISTONE H4 (HIS4, At2g28740) and the S-phase marker HISTONE H2A 6 (HTA6, At5g59870) in Col-0 control (mock) and 3 h antimycin A (AA) treated seedlings. Top panels show 10x Xenium sections for HIS4, HTA6 and a merged image (HIS4 + HTA6) with transcripts. Bottom panels show UMAP feature plots for single-nucleus RNA-seq (SN; top row) and spatial RNA-seq (10x Xenium; bottom row) for HIS4 and HTA6 in Col-0 mock and Col-0 AA, coloured by scaled expression. Tau scores above each UMAP summarise cell type specificity (1 = highly specific, 0 = broad). AA treatment sharpens HIS4 expression into S-phase enriched domains that overlap with HTA6-positive nuclei.

In summary, many transcriptional regulators and stress mediators show not only cell type specific induction but also clear adaxial–abaxial and tissue-level polarity in response to mitochondrial dysfunction. This spatial organisation, visualised by integrating snRNA-seq and subcellular-resolution spatial transcriptomics, highlights that mitochondrial retrograde signalling is deployed through a complex and structured regulatory landscape across the leaf.

## DISCUSSION

This study reframes the MRR as a multicellular program that is resolved at the level of cell state, tissue position, and leaf polarity. Although mitochondrial inhibition was imposed broadly across seedlings, the transcriptional output was neither uniform across cell types nor homogeneous within canonical identities. Instead, mitochondrial dysfunction redistributed nuclei across multiple transcriptional states, including stress-associated states that were already detectable at low abundance in untreated samples. These observations argue that retrograde signaling is deployed through a mosaic of cell states whose prevalence, transcriptional composition, and spatial arrangement are reshaped as mitochondrial function declines.

Bulk transcriptomes have provided a coherent view of mitochondrial stress markers and their kinetics, including the characteristic peak in transcript abundance after antimycin A treatment around 3 h in wild type plants (Ng et al., 2013a). By resolving the response across hundreds of thousands of nuclei, our data show how such bulk trajectories can emerge from state-level redistribution within tissues. Multiple clusters assigned to the same broad lineage behaved differently in response to treatment, indicating that cell identity alone does not predict the magnitude or direction of transcriptional change. This is most apparent within mesophyll and epidermal lineages, where some clusters contracted while others expanded over the same time window. Such behaviour supports a model in which mitochondrial perturbation shifts the occupancy of existing transcriptional states and promotes transitions into stress-associated states, rather than imposing a single program on all cells of a given type. Notably, ANAC017 that has characterised as a high regulator of MRS shows a relatively uniform constitutive expression, thus while it is absolutely required for MRS, its expression pattern alone (or that of one of its targets ANAC013) (Ng et al., 2013a, De Clercq et al., 2013), is not sufficient to explain the cell, cluster and spatial responses uncovered in this study. It is possible that it is post-transcriptionally activated in different cells, but this still leaves the questions of how the specificity of the response is achieved. Given the cell specific expression observed for a variety of other transcription factors, many of which do not require ANAC017 for induction, it is likely that the complex spatial patterns of response is achieved by a number of factors. For many genes that are markers of MRS, in addition to NAC transcription factor binding sites, WRKY, ABA and ERF binding has been experimentally demonstrated (Dojcinovic et al., 2005; Giraud et al., 2009; He et al., 2022; Van Aken et al., 2013; Vanderauwera et al., 2012). A study at a proteome level analying the response of mitochondria to carbon starvation for three or six day revealed differences between guard and mesophyll cells (Boussardon et al., 2025).

A consistent feature of the dataset is that clusters that expanded strongly under mitochondrial inhibition were not absent from controls. Instead, a small fraction of nuclei in untreated samples already occupied transcriptional states resembling those enriched after treatment. Treatment therefore increases the proportion of nuclei in these pre-existing states rather than generating entirely new states *de novo.* This behaviour is reminiscent of previously proposed sensor, primer, and environmentally responsive state (ERS) cell populations (Nobori et al., 2025). ERS cells were described in deep single-cell profiling of Arabidopsis roots, where subsets of cells across lineages showed heightened responsiveness to environmental cues and their abundance shifted with external conditions (Oliva et al., 2022). Whether such primed states reflect local microenvironmental differences, developmental asynchrony, or stochastic fluctuations in regulatory networks remains to be tested. It also appears that mesophyll cell types were more strongly affected than other cell types (Figure 2A and 2D). Caution needs to be taken in intreperting this due to the fact that mesophyll cell mitochondria dominate the mitochondrial mass of a leaf (Hater et al., 2025). Nevertheless, there greater response may be at least in part due to the fact that mesophyll cell undergo a strong diurnal rhythm in gene expression associated with photosynthesis. Previously, it has been shown that ANAC017 binds the same *cis*-promoter elements as a variety of diurnal regulators (Zhu et al., 2023), and thus diurnal periodicity may contribute to responsiveness on mesophyll cells to mitochondrial perturbation (Alber and Vanlerberghe, 2021). Also, mitochondrial function supports efficient photosynthesis by reducing excess reducing equivalents generated in photosynthesis, and thus any perturbation of mitochondrial function would have a greater effect in mesophyll cells as they dominate the photosynthetic capacity of leaves.

A central outcome of integrating single-nucleus and spatial transcriptomics is the demonstration that mitochondrial stress signaling has strong spatial organization. Some responses are broadly distributed, consistent with markers that are induced across many cell types, whereas other responses are spatially restricted to specific tissue domains or show clear adaxial–abaxial polarity. Such patterns are difficult to infer from dissociative profiling alone and are largely invisible in bulk measurements. These findings also prompt a re-evaluation of how mitochondrial-derived signals such as ROS, predominantly hydrogen peroxide in the context of MRR, and redox switches operate. Most *in vivo* measurements have been made at whole-organ or leaf-disc scale using redox and ROS sensors (Khan et al., 2024; Scherschel et al., 2024). Our results raise a central question: do most cells generate similar primary signals but interpret them differently, or do distinct transcriptional states generate qualitatively and quantitatively different upstream signals? Early work using cell-type-resolved ROS reporters already indicates that ROS production differs across cell layers (Dopp et al., 2023; Kock et al., 2025), which is consistent with the latter possibility and provides a direction for future work linking state identity to signal production.

Spatial heterogeneity in organelle function has precedent, particularly for chloroplasts. Differences between cell types have been documented at the level of physiology, including non-photochemical quenching (Gorecka et al., 2014), at the level of proteome composition such as aminotransferases and phosphoenolpyruvate transporters (Beltrán et al., 2018; Lundquist et al., 2014), and at the level of nucleic acids and signaling properties (Dopp et al., 2023; Fang et al., 2019). For mitochondria, proteomic studies have revealed differences between guard cell, vascular, and mesophyll mitochondria, including altered respiratory components and distinct starvation responses (Boussardon et al., 2025; Hater et al., 2025). Other work has reported differences in ATP production per mitochondrion and in the complement of alternative NAD(P)H dehydrogenases between guard cell and mesophyll mitochondria (Ditz et al., 2025). The spatial and polarity patterns we observe at the transcript level are consistent with this broader literature and argue that mitochondria, like chloroplasts, operate within a tissue context that shapes both basal activity and stress signaling outputs.

ANAC017 is a central regulator of mitochondrial retrograde signaling in Arabidopsis and controls major transcriptional outputs under mitochondrial dysfunction (He et al., 2022; Zhu et al., 2023). In line with this role, loss of ANAC017 altered both the redistribution of nuclei among stress-associated states and the gene-level programs deployed within clusters. Importantly, the differences were not limited to attenuation of canonical marker induction. Instead, they included genotype-dependent cluster behaviour and divergent temporal trajectories, consistent with ANAC017 functioning as an organizing node that consolidates upstream cues into a coordinated transcriptional response. An interpretation of this data is that ANAC017 supports rapid establishment of transcriptional homeostasis after mitochondrial perturbation. In wild type seedlings, transcriptome remodeling peaks around 3 h and then stabilizes, as previously reported in bulk profiles (Ng et al., 2013a). In *anac017*, the continued accumulation of transcript changes suggests that compensatory routes respond more slowly, or with altered specificity, leading to prolonged remodeling rather than consolidation. This model fits with genetic evidence that ANAC017 activates downstream regulators while also engaging feedback control, providing both amplification and restraint within the network (He et al., 2022; Zhu et al., 2023). Furthermore, our transcription factor analysis reinforces that regulatory control of mitochondrial stress signaling is deployed in a cell-state-specific manner. Many transcription factors retained high specificity across control and treatment conditions, indicating that identity programs remain robust even as stress states expand. At the same time, subsets of regulators altered their specificity with treatment, either extending into additional cell types or becoming more spatially restricted.

Despite numerous single-cell and single-nucleus studies in Arabidopsis, many clusters still lack experimentally validated cell-type markers, even with two recent Arabidopsis Atlas studies providing substantial marker information (Guo et al., 2025; Lee et al., 2025). This suggests that while some markers appear to be robust over development and treatment, others are more developmentally and growth condition dependent. To overcome this barrier we did carry out a comprehensive validation process using spatial and promoter GFP-tagging. Many promoter GFP reporters did not match the predicted localization, which is likely to reflect limitations of promoter fragments and transgene context. Short promoter regions can miss distal regulatory elements that restrict expression in the native locus, and insertion site or local chromatin effects can broaden reporter activity. This is supported by the observation that of the 29 cell type markers that overlapped between the 10x Xenium and GFP based systems, only six gave a similar pattern in GFP and 10x Xenium (Supplemental Table 9). Thus, spatial transcriptome markers were essential for confirming or assigning cell types and for tracking how these assignments changed with treatment. Bulk sequencing already shows that treatment responses are highly variable; for example, drought meta-analyses identify shared and previously missed genes, yet overlap between studies often represents only ∼25% of the genes detected in any single study (Rest et al., 2016), with a similar pattern reported for ROS responses (Willems et al., 2016). Single-cell transcriptome studies are likely to reveal an even greater diversity, shaped not only by developmental stage, tissue context, and growth conditions that influence bulk sequencing, but also by the number of cells profiled, which strongly affects clustering, sensitivity, and resolution.

In conclusion, this study provides an in-depth view of the MRR at cellular, spatial and polarity levels in *Arabidopsis* seedlings. We show that mitochondrial dysfunction elicits highly structured transcriptional changes across cell types, within cell type specific clusters and along adaxial–abaxial and tissue axes. These principles are likely to extend to other abiotic stresses. In addition, the cell type specific marker sets and information on whether their specificity is maintained or remodelled upon stress offer a resource for future work that aims to dissect abiotic stress responses at cellular resolution.

## Supporting information

All Supplemental Figures

## SUPPLEMENTAL FIGURE LEGENDS

**Supplemental Figure 1. 10x Xenium spatial validation of marker genes used for cell-type annotation**

10x Xenium *in situ* hybridization was used to validate marker genes for all major cell types and clusters identified in 14-day-old *Arabidopsis thaliana* seedling leaves after mock or antimycin A treatment. For each marker gene, fluorescence images show 10x Xenium probe signal in cross sections of Col-0 leaves, alongside feature plots of single-nucleus RNA-seq and 10x Xenium spatial UMAPs that display normalized expression.

Mesophyll identity is supported by markers including UDP-glycosyltransferase 85A1 (UGT85A1), Beta carbonic anhydrase 1 (CA1), Allene oxide cyclase 2 (AOC2), Beta carbonic anhydrase 2 (CA2), Suppressor of Yeast gpa1, Phosphate-regulated 81 and Xenotropic and Polytropic Retrovirus receptor family protein 1–like protein 1 (SPX1), Ribulose-1,5-bisphosphate carboxylase/oxygenase small subunit 2B (RBCS2B) and Early responsive to dehydration 1 (ERD1). Epidermal and leaf pavement cell identity is confirmed with markers such as Cysteine-rich receptor-like protein kinase 42 (CRK42), Lysine histidine transporter 1 (LHT1), Homeobox-leucine zipper protein 2 (HAT2), Beta-galactosidase 4 (BGAL4), MYB domain protein 96 (MYB96), Glycosylphosphatidylinositol-anchored lipid transfer protein 1 (LTPG1), Short hypocotyl 2/Auxin-responsive protein IAA3 (SHY2), YABBY domain protein 5 (YAB5), Lipid transfer protein 1 (LTP1), Beta-glucosidase PYK10 (PYK10) and Methionine sulfoxide reductase B6 (MSRB6).

**Supplemental Figure 2. Confirmation of Cell markers using Promoter GFP tagging.**

Transverse sections of leaves expressing promoter::GFP fusions for 52 candidate marker genes are shown as GFP, chlorophyll autofluorescence, bright-field and merged images at the mid-vein, lamina middle and edge. Predicted cell-type enrichment based on single-nucleus and spatial transcriptome data is indicated above each panel, and the experimentally observed pattern is indicated below. Guard-cell, epidermal and mesophyll markers were successfully validated for the first 9 genes. The remaining 43 reporters showed broad, weak or ambiguous expression across tissues and therefore did not confirm specific or enhanced expression in the predicted cell types.

**Supplemental Figure 3.** Example of an epidermal, mesophyll and companion cell marker from all sections used in this study. All sections and genes can be viewed in 10x Xenium Explorer with the links provided in data availability. Antimycin A: AA.

**Supplemental Figure 4. Sub-clustering resolves heterogeneous cell states within three stress-responsive unknown clusters.**

Single-nucleus RNA sequencing data were re-analysed by extracting nuclei from cluster 3, cluster 7 and cluster 11 and performing independent unsupervised subclustering for each parent cluster. For each parent cluster, the left panel shows a UMAP embedding of nuclei coloured by subcluster identity (numbers indicate subclusters). The legend adjacent to each UMAP indicates the dominant cell-type assignment for each subcluster based on marker enrichment (ME, mesophyll; LPC, leaf pavement cell; EP, epidermis; LGC, leaf guard cell; XYL, xylem; PP, phloem parenchyma; SP, S phase; UK, unknown), highlighting that each parent cluster contains multiple transcriptionally distinct states.

For each parent cluster, the right panel shows a dot plot of the top two marker genes per subcluster, ranked by differential expression within that subclustering analysis. Dot colour intensity represents average scaled expression, and dot size represents the percentage of nuclei within each subcluster expressing the gene. The x-axis indicates subcluster identity, and the y-axis lists marker genes. As outlined in Figure 1 the predicted cell types were confirmed with using 10x Xenium Chip cellular identification (red dot), promoter GFP localization (green dot) or verified published markers (black dot). (see Supplemental Table 1 for cell annotation).

**Supplemental Figure 5. Time-resolved shifts in cluster abundance during mitochondrial inhibition in Col-0.**

Nuclei counts and relative abundance for each snRNA-seq cluster (rows, clusters 0–27) in Col-0 after 1 h, 3 h, 6 h, and 12 h under mock, antimycin A (AA), or myxothiazol (MXO) treatment (columns). For each time point, AA and MXO are compared to the time-matched mock control. Asterisks denote clusters with a significant change in abundance relative to the matched mock control, with red indicating an increase and blue indicating a decrease. Total nuclei analysed per condition are shown at the bottom of each column. Student’s t-test with p ≤ 0.05, n = 3.

**Supplemental Figure 6**. **The transcriptional response to mitochondrial dysfunction after 1 (A), 6 (B) and 12 h (C) treatment with antimycin A or myxothiazol.** The left panel depicts the changes in the number of nuclei after 1, 6 and 12 h treatment. The clusters are sorted according to cell type. A red asterisk indicated a significant increase in number and a blue start indicates a significant decrease in number. The right panel depicts examples of clusters that are significantly increased in the treated compared to control samples. (see Supplemental Table 1 for cell annotation).

**Supplemental Figure 7. The response of Col-0 and *anac017* to antimycin A (AA) treatment at A) 1 and B) 6 h.** Three different comparisons were carried out i) The response of Col-0 mock to AA, indicated with blue, ii) Col-0 mock treated compared to *anac017* mock treated, indicated in purple, and iii) the response *of anac017* to antimycin A treatment, indicated in green. A red asterisk indicated a significant increase in number and a blue asterisk indicates a significant decrease in numbers.

**Supplemental Figure 8. Unsupervised cell cluster of Col-0 and *anac017* at A) 1, B) 3, and C) 6 h treatment with antimycin A.** The samples were clustered using Sauret. The cell clusters were initially annotated using Single Cell Plant DataBase (scPlantDB) (Ryu et al., 2021) and subsequently the cell types were confirmed with using 10x Xenium Chip cellular identification (red dot) and promoter GFP localization (green dot). For the purposes of reproducibility the clusters initially annotated as unknown are left annotated as unknown in this diagram in black text, with subsequent identification by 10x Xenium Chip and promoter GFP tagging identified the cell type, which is indicated in red (confirmed using 10x Xenium Spatial Transcriptome) or green (using promoter GFP-tagging) (see Supplemental Table 4 for cell annotation). mesophyll (ME), epidermis (EP), leaf pavement cells (LPC), bundle sheath (BS), phloem (PH), phloem parenchyma (PP), companion cells (CC), leaf guard cells (LGC), xylem (XYL), G2/M phase (G2M), S phase (SP) and unknown (UK).

**Supplemental Figure 9. A gene level view of the mitochondrial stress response at a cellular level at A) 1, B) 3, C) 6 and D 12 h. Top Panel -** The number of genes that changed in transcript abundance after 1, 3, 6 and 12 hours of antimycin A (AA) and myxothiazol (MXO) treatment (see Supplemental Table 6 for full details). mesophyll (ME), epidermis (EP), leaf pavement cells (LPC), phloem parenchyma (PP), xylem (XYL), bundle sheath (BS), phloem (PH), companion cells (CC), leaf guard cells (LGC), G2/M phase (G2M), S phase (SP) and unknown (UK). **Venn Diagrams** - The number of genes that were increased or decreased in transcript abundance after 1-, 3-, 6- and 12-hours treatment with antimycin A or myxothiazol and how they overlap in epidermal (EP), leaf pavement cells (LPC) and mesophyll cells (ME). **Bottom Panel -** The common and unique genes that increased or decreased in the epidermal, leaf pavement cell, mesophyll and bundle sheath clusters after 1-, 3-, 6- and 12-hours treatment with antimycin A or myxothiazol if there are two or more clusters in the same cell type.

**Supplemental Figure 10. The number and overlap in differentially expressed genes (DEGs) in Col-0 and *anac017* after A) 1, B) 3, and C) 6 hours of antimycin A treatment.** Venn diagrams show differentially expressed genes (DEGs) in Col-0 (blue) and *anac017* (green) after antimycin A treatment for (A) 1 h, (B) 3 h, and (C) 6 h, resolved by snRNA-seq cluster (cluster ID and cell type indicated). Numbers denote total and shared DEGs, with up- and downregulated genes indicated where shown.

**Supplemental Figure11. Analysis of differentially expressed genes in Col-0 and *anac017* after A) 1, B) 3, and C) 6 hours of treatment with antimycin A. i)** UMAP cluster visualization of Col-0 and *anac017* after 1, 3 and 6 h treatment showing cell type that were predicted and re-annotated with experimental verification as outlined in Figure 2**. ii)** The top two marker genes in each cluster (see Supplemental Table 4). **iii)** The number of nuclei in each cluster in control (mock) and antimycin A (AA) treated samples after different times of treatment. Changes in the number of nuclei compared to control are indicated. Student’s t-test with p ≤ 0.05, n = 3. Three different comparison were carried out i) The response of Col-0 to AA, indicated with blue, ii) Col-0 mock treated compared to *anac017* mock treated, indicated in purple, and iii) the response *of anac017* to antimycin A treatment, indicated in green. A red asterisk indicated a significant increase in number and a blue asterisk indicates a significant decrease in numbers. **iv)** The total number of differentially expressed genes (DEGs) and genes encoding transcription factors in each cluster in Col-0 and *anac017* after **A)** 1, **B)** 3, and **C)** 6 hours of treatment with AA. The cluster number and cell type are indicated in the outer circle with clusters experimentally confirmed as in Figure 3. The yellow asterisks indicate a significance (p < 0.001) in the proportion of genes that changed in transcript abundance in Col-0 compared to *anac017* for each cluster based on all the total number of genes detected. The number of transcription factors is too small to carry out a proportionality test. mesophyll (ME), epidermis (EP), leaf pavement cells (LPC), phloem parenchyma (PP), xylem (XYL), bundle sheath (BS), phloem (PH), companion cells (CC), leaf guard cells (LGC), G2/M phase (G2M), S phase (SP) and unknown (UK). Clusters re-designated using experimentally based markers are indicated in brackets.

**Supplemental Figure 12. Single nuclei and spatial transcriptome generated UMAP projections of cell clusters after treatment with antimycin A for 3 h in Col-0 and *anac017*.** Each panel shows an individual gene with abbreviation and gene identity. The section for each treatment represents a single section of a total of 20-24 section for each treatment. The feature plots depict the UMAP projection for the snRNA-seq (top row in each panel) and spRNA-seq (bottom row in each panel). The calculated tau value, as a measure of expression specificity, where 1 is most specific and 0 is non-specific is shown. The feature plot clusters and cell types are designated in Figure 4. *GSTF6 = Glutathione s-transferase 6*, GSTF7 = *Glutathione S-transferase 7,* mock = control, AA = Antimycin A, SN = single nuclei.

**Supplemental Figure 13. Spatial transcriptome patterns of genes encoding transcription factors.** Each panel shows an individual gene with abbreviation and gene identity. The section for each treatment represents a single section of a total of 24 sections for each treatment (12 after 3 h and 12 after 6 h of treatment with AA). The feature plots depict the UMAP projection for the snRNA-seq (top row in each panel) and spRNA-seq (bottom row in each panel). The calculated *tau* value, as a measure of expression specificity, where 1 is most specific and 0 is non-specific is shown. The feature plot clusters and cell types are designated in Figure 5. *FAMA* = bHLH transcription factor, *HB-8* = Homeobox gene 8, MYB28 and MYB29 = MYB domain protein 28 and 29, mock = control, AA = Antimycin A, SN = single nuclei.

**Supplemental Figure 14. Images of nuclei after purification by flow cytometry.** The nuclei were free of debris and were observed to be intact, free of doublets and thus judged to be suitable for sequencing using microfluidics-based lysis and library preparation. Scale bar = 10 μm.

## SUPPLEMENTAL TABLES

**Supplemental Table 1. List of genes that were used in the spatial transcriptome analysis.** The list of 465 genes and the reason they were used in the analysis. Note that the design is an iterative process and some genes initially selected failed the manufacturer’s quality control and could not be included. The final design and the list of genes used to design are shown, from the differential gene expression observed with the snRNAseq data, predicted markers for each cluster and sub-cluster, published markers at the time of design, gene markers of mitochondrial dysfunction (SPD) and genes encoding proteins for various functions in mitochondria.

**Supplemental Table 2. Annotation of cell types for antimycin A and myxothiazol Col-0 samples after 1-, 3-, 6- and 12-hours treatment.**

Cell types were initially predicted using various Plant Cell databases and then experimentally confirmed using a combination of 10x Xenium spatial transcriptome and promoter GFP-tagging approaches. Text in black is predicted cell type annotated from databases, red is annotated from 10x Xenium spatial transcriptome (see Supplemental Figure 1 for images of all spatial markers), and green is annotated from promoter GFP-tagging (see Supplemental Figure 1 for images of all promoter-GFP markers).

**Supplemental Table 3. Nuclei numbers, median UMIs and median genes detected per nucleusin samples.**

The number of nuclei for each sample that was used as input for the 10x workflow to obtain single nuclei sequencing. Note that the number of nuclei in each sample will vary by a small extent with different analysis, while the sample cut-off imposed was the same in all the analysis, the different samples included, i.e. all verse single time, will result in slightly different numbers being retained for the analysis. So for the same sample if all samples are clustered together **(A)** (i.e. 1, 3, 6 and 12 h mock and treated with antimycin A and myxothiazol) the number may be slightly different than a single time point is re-clustered **(B)** (for pseudobulk analysis for determining differentially expressed genes). **(C)** median UMIs and median genes detected per nucleus in each sample from 2 analysis groups.

**Supplemental Table 4. Definition of Cell Types for antimycin A treated Col-0 and *anac017* samples after 1, 3, and 6 hours treatment.** Cell types were initially predicted using various Plant Cell databases and then experimentally confirmed using a combination of spatial transcriptome and Promoter GFP-tagging approaches.

**Supplemental Table 5. Nuclei numbers in Col-0 and *anac017* samples at 1, 3 and 6 hours after treatment with antimycin A.** The number of nuclei for each sample that was used as input for the 10x workflow to obtain single nuclei sequencing.

**Supplemental Table 6. Differential gene expression for Col-0 plants at a cell and cluster level treated with antimycin A and myxothiazol for 1, 3, 6 and 12 hours.**

**Supplemental Table 7. Differential gene expression for Col-0 and *anac017* plants at a cell and cluster level treated with antimycin A for 1, 3 and 6 hours.**

**Supplemental Table 8. *Tau* scores for 1119 all transcription factors detected in this study.**

**Supplemental Table 9. The list of promoter-GFP constructs used to confirm cell identity and they overlap with the 10x Xenium spatial transcriptome markers.**

## Methods

### Plant growth and treatment antimycin A and myxothiazol

*Arabidopsis thaliana* Col-0 and *anac017* seeds were surface sterilized by incubation in 75% (v/v) ethanol and 0.1% Triton X-100 (v/v) for 5 min, followed by three times incubations in 75% (v/v) ethanol for 5 min. Seeds were dried on sterilized filter paper. Sterilized seeds were placed on Gamborg’s (B5) medium with 1% (w/v) sucrose, 0.8% (w/v) agar and stratified for 48 h at 4°C in the dark, and grown under a 16 h: 8 h day: night photoperiod at 23°C with 100 μmol.m^-2^.s^-1^ photosynthetic photon flux density. Plants grown for 14 days were sprayed with antimycin A (50 μM) or myxothiazol (50 μM) in 0.01% (v/v) Tween-20 solution, or with 0.01% (v/v) Tween-20 alone as mock treatment.

GFP tagged lines with predicted marker genes were grown for 3 weeks on a 3:1 soil vermiculite mixture in 5 cm^2^ pots with a volume of 200 ml with 16 h: 8 h day: night photoperiod at 23°C with 100 μmol.m^-2^.s^-1^ photosynthetic photon flux density.

### Nuclei extraction and single-nucleus sequencing

Plants shoots treated with antimycin A or myxothiazol for 1h, 3h, 6h or 12 h were harvested and 1 g fresh weight plant shoots were manually chopped with a razor blade in 3 mL freshly prepared nuclei isolation buffer (400 U/mL RNase inhibitor, 1.25 mM DTT, 0.2 mM Spermine and 0.5 mM Spermidine in Mobi^TM^ Plant Nuclear Extraction Buffer A (MOBIDROP, China)) on ice for 5 min to release nuclei. Three biological replicates were performed. The crude nuclei extract was filtered through 30 μm cell strainers and centrifugated at 500 g for 5 min at 4°C to pellet the nuclei. The pellet was resuspended in 1 mL nuclei resuspension buffer (2% (w/v) BSA, 200 U/mL RNase inhibitor and 1.25 mM DTT in PBS buffer) and 10 μL 100 μg/mL DAPI. Fluorescence activated nuclei sorting (FANS) was carried out using a by BD FACSAria-III flow cytometer. Nuclei were identified by making an initial gate on a bi-plot of scatter at a ninety-degree angle relative to the laser (SSC) versus a DAPI peak. Debris were removed by plotting the DAPI emission channel against the height or area forward scatter (FSC) and SSC value. Doublets and clumps were removed by plotting each DAPI emission channel’s height and area against each other (Berendzen et al., 2023). Finally, a small number of nuclei were stained with 0.2% (w/v) Trypan blue to visualize the morphology under microscope. Intact nuclei were round and debris free, confirming their quality for subsequent analysis (Supplemental Figure 14). Nuclei were then counted using cell analyzer (Countstar Rigel S3, China). The purified nuclei were loaded into a 10x Genomics Chip and barcoded with a Chromium Controller (10x Genomics). Reverse transcription mRNA and sequencing libraries were constructed with a Chromium Next GEM Single Cell 3□ Kit v3.1 (10x Genomics) according to the manufacturer’s instructions. Sequencing was performed with a NovaSeq 6000 instrument (Illumina).

### Data processing and analysis

Raw fastq files were analysed by Cell Range v 7.2.0 (10x Genomics) to generate gene by cell matrices and mapping to the TAIR10 reference genome (https://www.arabidopsis.org/) and Araport11 annotations. Cells containing mitochondrial or chloroplast encoded genes were removed. The output from Cell Ranger was further analysed with Seurat v4.1.1 (10x Genomics). Doublet filtering was performed on each dataset by using the method of DoubletFinder. Nuclei that did not meet the cutoff of > 500 genes were removed. Cells were then normalized using the LogNormalize method, factor = 10000. The method of vst was used to find variable features, nfeatures = 2000.

Dimensionality reduction was performed with PCA. The "RunHarmony" function was then performed to reduce batch effects between samples (Korsunsky et al., 2019). Construction of the shared nearest neighbours and nuclei clustering was performed using the FindNeighbors and FindClusters (Louvain method, resolution 0.8) functions, respectively. Sub-clustering was carried out with similar parameters using the cell population of the unknown cluster. RunUMAP was used to visualize the data, with the parameter (dims = 1:20, n.neighbors=30, min.dist=0.3). FindAllMarkers function was used to identify cluster markers for each cluster with return.thresh=0.01 and logfc.threshold = 0.26.

### Cell type annotation

Cell type was initially annotated from three databases scPlantDB, PlantCellMarker, and PlantscRNAdb (Chen et al., 2021; He et al., 2024; Or and Wang, 2014). This initial predicted annotation was left on all UMAPs. The top ten markers from these predictions are listed in the Supplemental Tables associated with the Figures, and the cell types were confirmed using spatial transcriptomics using a 10x Xenium platform and promoter GFP tagging, that are indicated on the Figures in red and green respectively. The 10x Xenium and GFP markers are shown in Supplemental Figure 2 and 3).

### Generation of transgenic GFP tagged lines

Promoters (∼ 2 kb upstream of the start codon) of genes (Supplemental Table 9) were cloned to pBWA(V)BII vectors (Biorun Biosciences Co., Ltd, Wuhan, China) using Golden Gate Assembly technology according to manufacturer’s instructions (NEB) to drive the expression of GFP. The primer details were listed in Table Supplemental Table 9. All resulting plasmids were transformed into *Agrobacterium tumefaciens* GV3101 strain and subsequently transformed into *Arabidopsis thaliana* Col-0 by floral dipping as previously described (Clough and Bent, 1998). Transgenic plants were screened on 1/2 MS plates supplemented with 5 mg/L glufosinate ammonium. About 10 independent T1 lines for each gene promoter were analysed for GFP signals.

### Confocal settings

Fluorescence images were acquired using a Zeiss LSM 710 confocal microscope equipped with a 25× water-immersion objective. GFP was excited with a 488 nm argon laser, and emission was collected at 500–530 nm. Chlorophyll autofluorescence was excited with the same 488 nm laser and detected at 650–750 nm.

### 10x Xenium-based spatial transcriptomics

Fresh Arabidopsis leaves from 14 days old plants were collected and fixed in FAA (G1108, Servicebio, China) for 20 h, with the fixative replaced after the first 10 h. The fixed leaves were dehydrated through a graded ethanol series: 75% of ethanol (v/v, 30 min), 85% ethanol (v/v, 30 min), 95% ethanol (v/v, 30 min), and 100% ethanol (v/v, 30 min). Subsequently, the samples were immersed in tissue clearing agent (GS-06, GreenSpecimen, China) and then immersed with molten paraffin. Three leaves from different plants were embedded in paraffin in one embedding mold. The embedded leaf samples were sectioned into 10 μm slices using a microtome (Minux S700, RWD Life Science, China). The sections were gently lifted with forceps and floated on a 40°C water bath to flatten. A 10x Xenium Gene Expression panel was carefully submerged into the water and lifted vertically to transfer the section onto the panel. Each panel was profiled across 4 sections. The panels were dried in a 42°C oven for 3 hours and then transferred to a desiccator overnight at 25°C to ensure complete moisture removal. Probe hybridization, ligation, and rolling circle amplification were performed according to the manufacturer’s protocol (10x Genomics, CG000749, CG000584). Background fluorescence was chemically quenched, and imaging and signal decoding were conducted using the 10x Xenium Analyzer instrument (10x Genomics). The leaf sections were stained in 0.04% FB28 (R053752, Rhawn, China) in the dark for 1 min, followed by image acquisition using a Digital scanner (Pannoramic MIDI II, 3DHISTECH, Hungary).

The 10x Xenium Gene Expression panel was custom-designed with 465 genes (Supplemental Table 1). From this gene list, markers were selected to validate the identity of each predicted cluster. These included genes drawn from cluster-specific marker sets,, 140 genes selected from those differentially expressed in Col-0 or *anac017 ko-1* after AA treatment, 38 genes curated from previously published cell type marker studies, and 59 genes known to response to mitochondrial stress, perturbation or dysfunction. The selection further encompassed genes encoding mitochondrial proteins involved in core mitochondrial functions, such as protein import and respiration. The 10x Xenium Gene Expression panel were designed by the 10x Xenium Panel Designer (https://cloud.10xgenomics.com/xenium-panel-designer), with the following requirements, 1) a single gene < 120 TP10k (transcripts per 10,000) in all cell types, 2) total panel utilization (all genes combined) < 600 TP10k, 3) each gene has >2 probe sets.

## Data availability

The raw single nuclei RNA sequencing data generated in this study have been deposited in the Genome Sequence Archive (GSA) at the National Genomics Data Centre (NGDC), China National Centre for Bioinformation (CNCB), under BioProject accession number PRJCA048906. The data are publicly accessible at https://ngdc.cncb.ac.cn/bioproject/browse/PRJCA048906.

The raw data for the spatial RNA sequencing generated in this study is available at Zenodo (https://zenodo.org).

## Author Contributions

Conceptualization: Y.Z., J.W.

Data Curation: Y.Z., M.L.,

Formal Analysis: M.L., Y.Z., F.H., X.Z., X. W., Y. L., R.S.

Funding acquisition: J.W., H.S., M.L.,

Investigation: Y.Z., M.L., J. L., C.Z.

Methodology: Y.Z., M.L., J. L., C.Z.

Project Administration: J.W.

Resources: J.W., M.L., H.S.

Software: F.H., X.Z., X.W., Y.L.

Supervision: J.W. H.S. G.A.K., M.L.

Validation: Y.Z., M.L., J. L., R.S., C.Z.

Visualisation: Y.Z., M.L., J.L., G.A.K., M.L., J.W.

Writing – original draft: Y.Z., M.L., G.A.K. J.W.,

Writing – review and editing. Y.Z., M.L., J. L., G.A.K. H.S. M.L. J.W.,

## Acknowledgments

Support Funding from Zhejiang University, and the Yangtze River Fellowship scheme to J.W.; Australian Research Council (DE210101200) to G.A.K.; Australian Research Council CE230100015, DP220102840, IH240100024 to M.G.L.

