## Supplementary material for "Mitochondrial Retrograde Signaling in *Arabidopsis thaliana*: heterogenous, spatial and polarised aspects": All Supplemental Figures

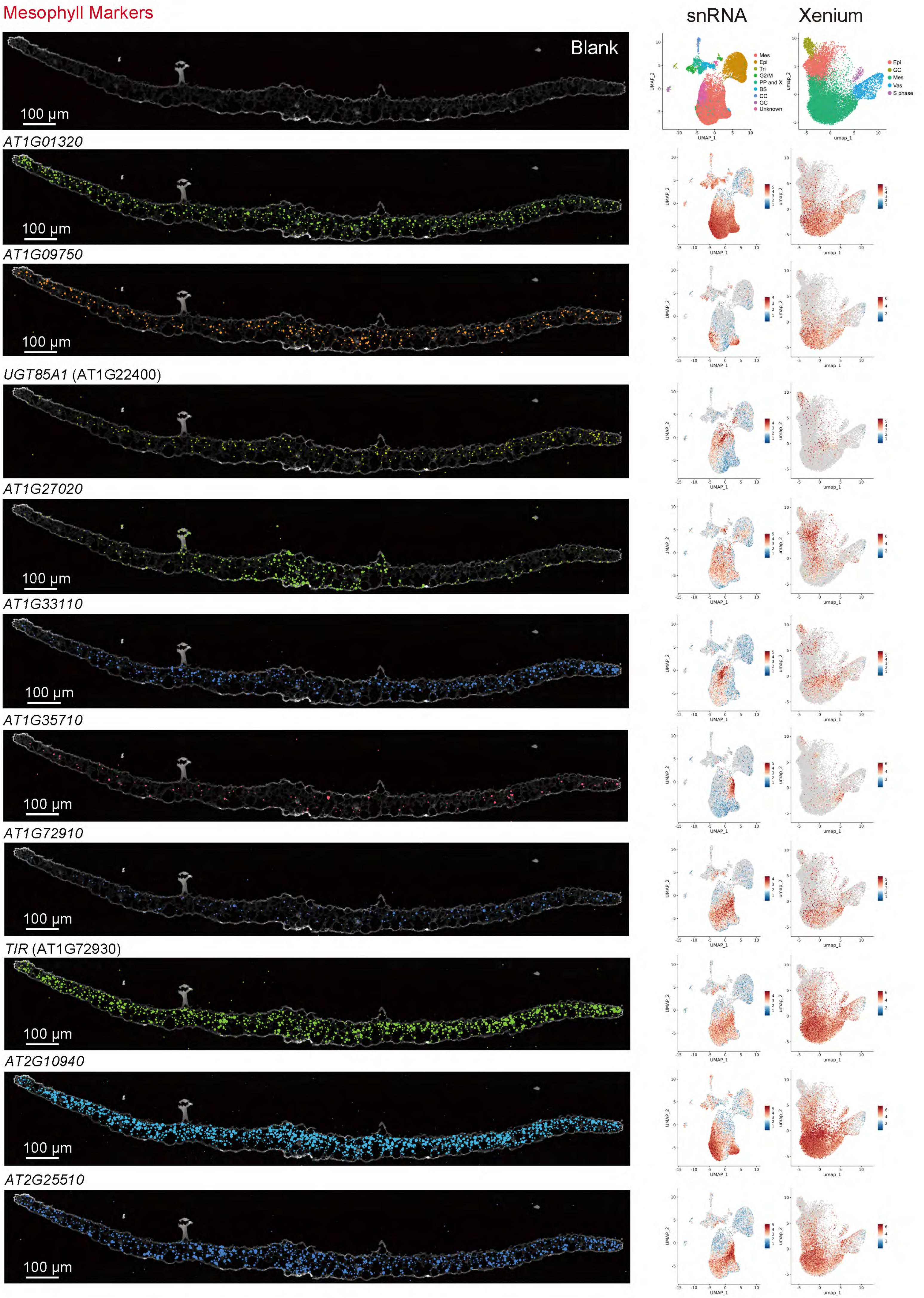
Mesophyll Markers
snRNA
Xenium
Blank
100 μm
Mes
Epi
Tri
G2/M
PP and X
BS
CC
GC
Unknown
Epi
GC
Mes
Vas
S phase
UMAP_1
UMAP_2
umap_1
umap_2
AT1G01320
AT1G09750
UGT85A1 (AT1G22400)
AT1G27020
AT1G33110
AT1G35710
AT1G72910
TIR (AT1G72930)
AT2G10940
AT2G25510

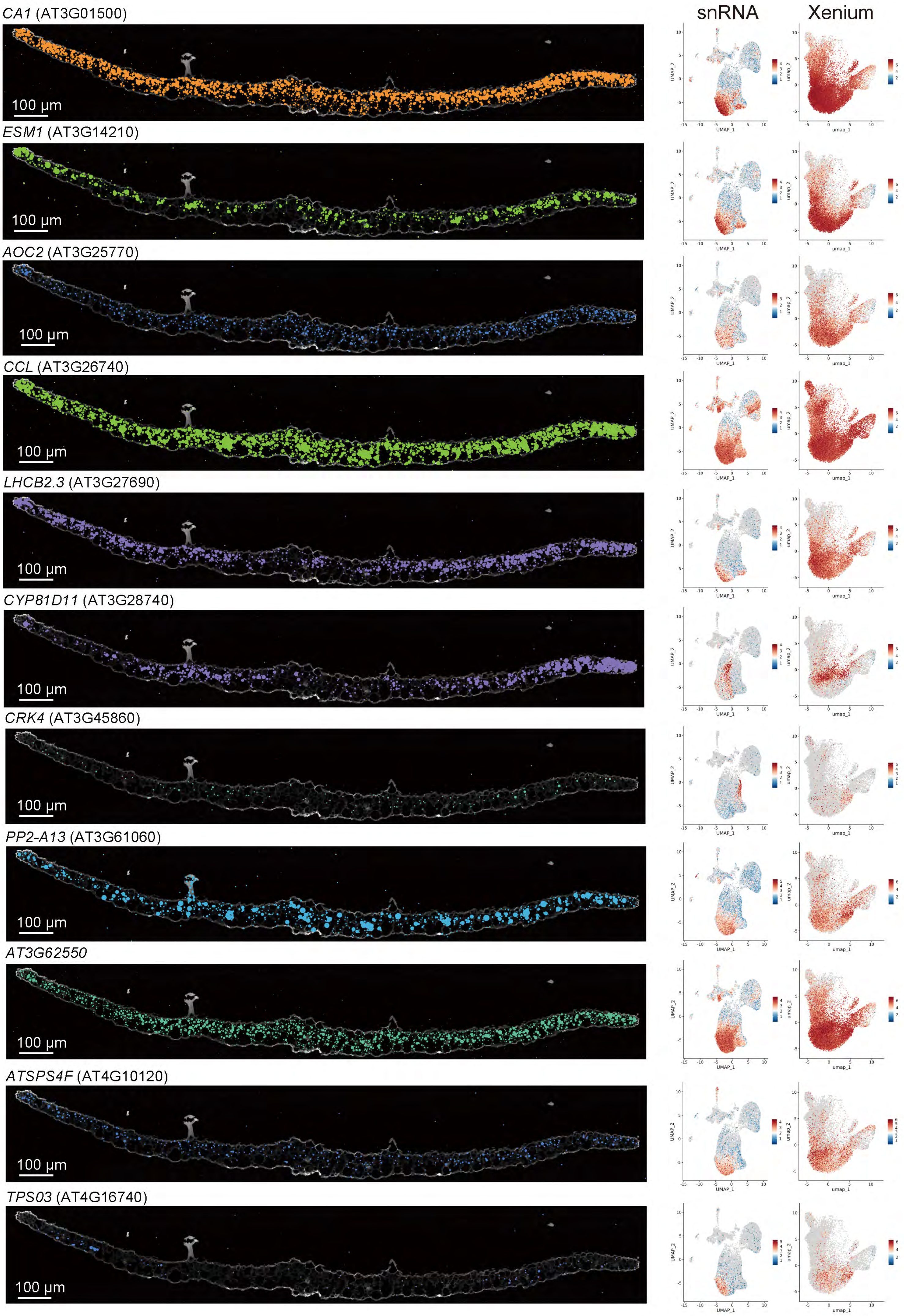
snRNA
Xenium
CA1 (AT3G01500)
ESM1 (AT3G14210)
AOC2 (AT3G25770)
CCL (AT3G26740)
LHCB2.3 (AT3G27690)
CYP81D11 (AT3G28740)
CRK4 (AT3G45860)
PP2-A13 (AT3G61060)
AT3G62550
ATSPS4F (AT4G10120)
TPS03 (AT4G16740)
100 µm
UMAP_1
UMAP_2
umap_1
umap_2

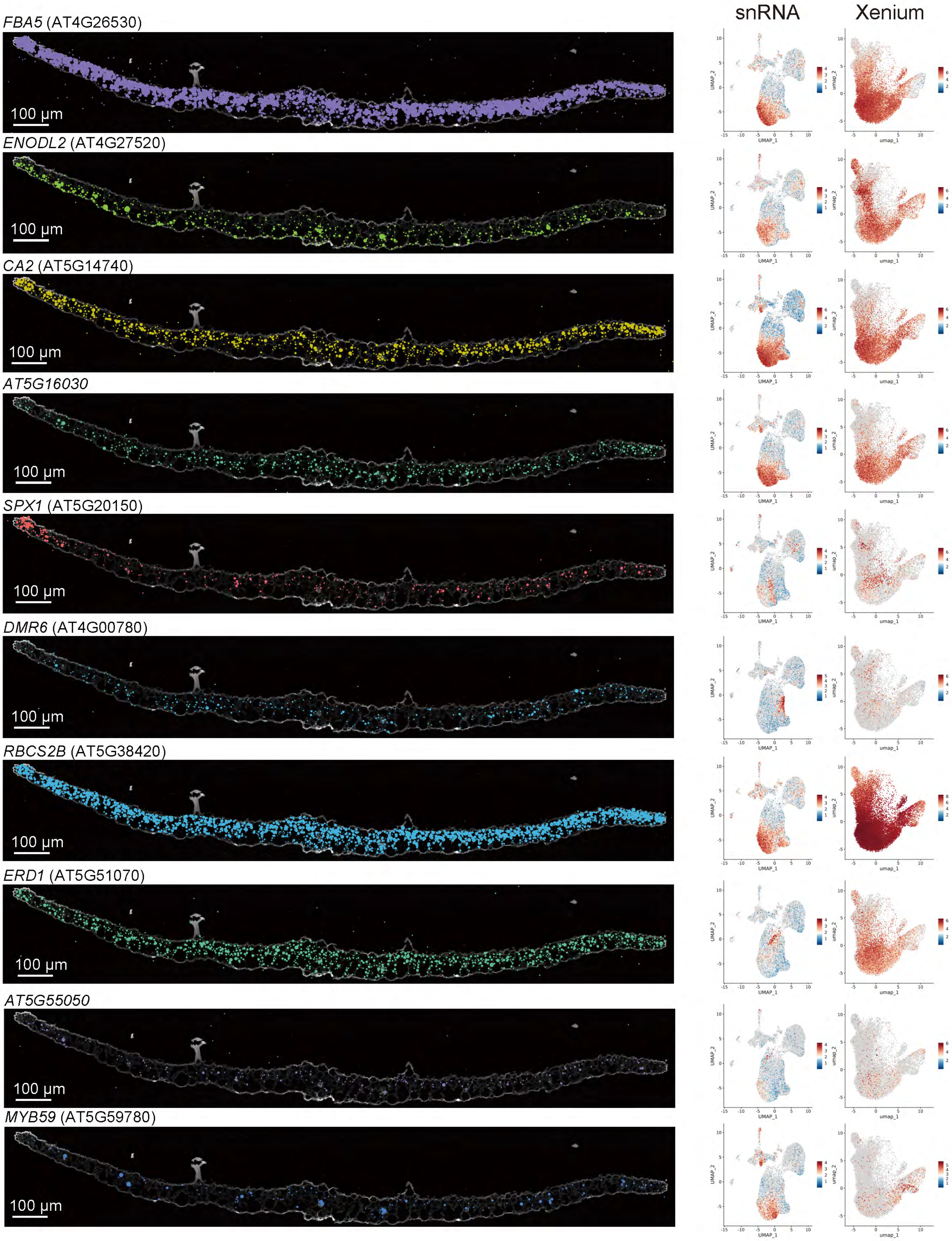
snRNA
Xenium
FBA5 (AT4G26530)
ENODL2 (AT4G27520)
CA2 (AT5G14740)
AT5G16030
SPX1 (AT5G20150)
DMR6 (AT4G00780)
RBCS2B (AT5G38420)
ERD1 (AT5G51070)
AT5G55050
MYB59 (AT5G59780)
100 µm
UMAP_1
UMAP_2
umap_1
umap_2

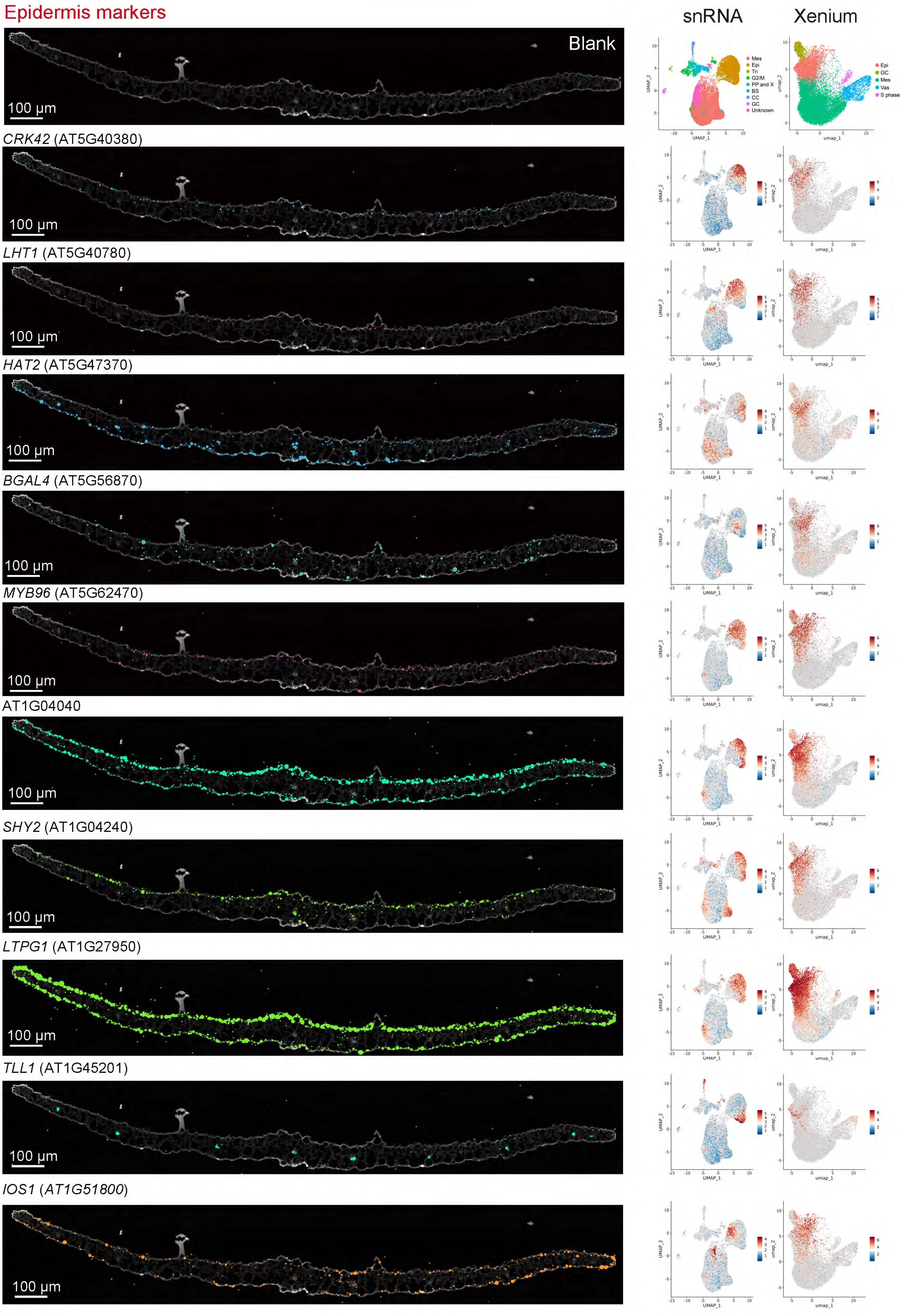

Epidermis markers
snRNA
Xenium
Blank
100 µm
CRK42 (AT5G40380)
LHT1 (AT5G40780)
HAT2 (AT5G47370)
BGAL4 (AT5G56870)
MYB96 (AT5G62470)
AT1G04040
SHY2 (AT1G04240)
LTPG1 (AT1G27950)
TLL1 (AT1G45201)
IOS1 (AT1G51800)
Mes
Epi
Tri
G2/M
PP and X
BS
CC
GC
Unknown
Epi
GC
Mes
Vas
S phase
UMAP_1
UMAP_2
umap_1
umap_2

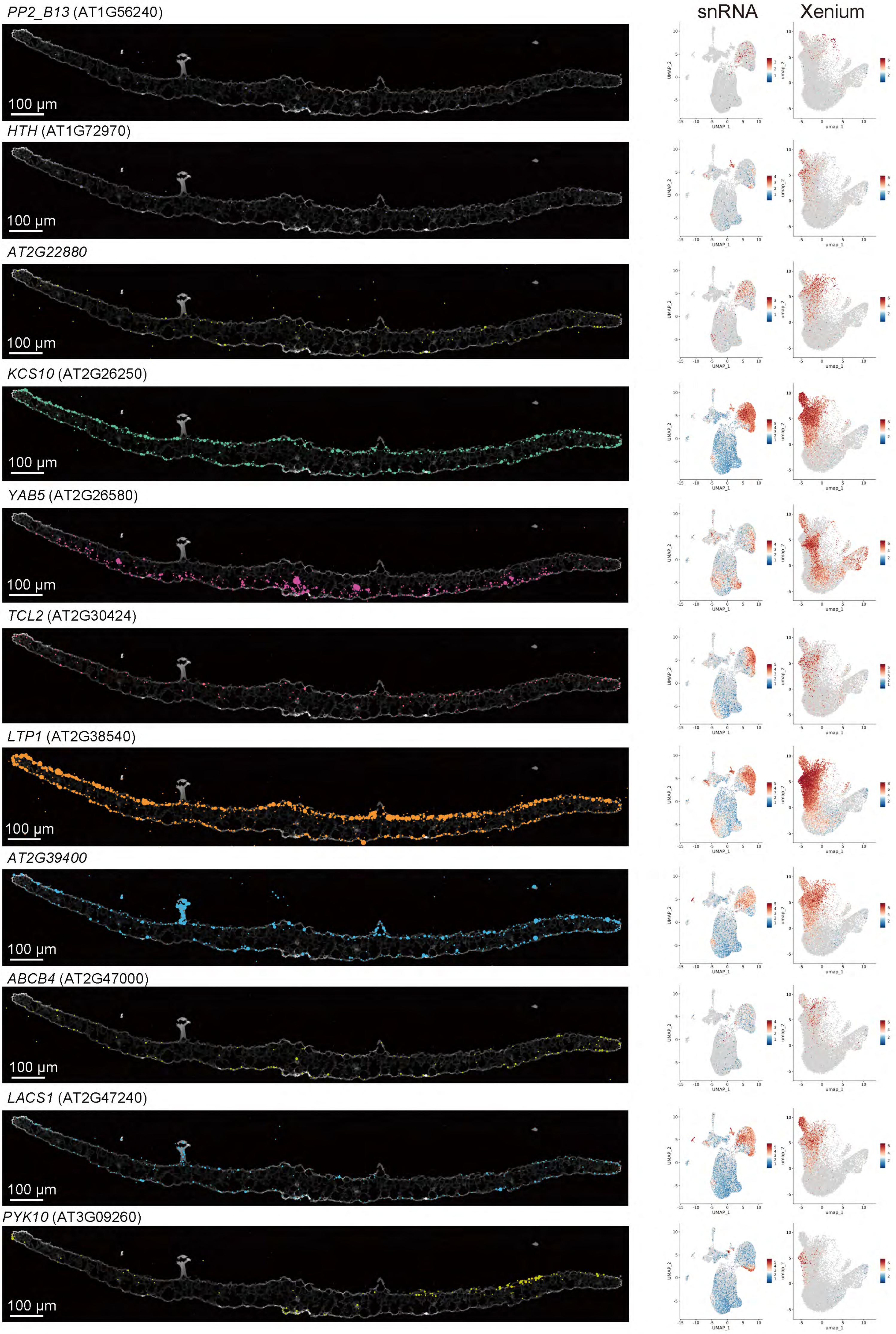
snRNA
Xenium
PP2_B13 (AT1G56240)
HTH (AT1G72970)
AT2G22880
KCS10 (AT2G26250)
YAB5 (AT2G26580)
TCL2 (AT2G30424)
LTP1 (AT2G38540)
AT2G39400
ABCB4 (AT2G47000)
LACS1 (AT2G47240)
PYK10 (AT3G09260)
100 µm
UMAP_1
UMAP_2
umap_1
umap_2

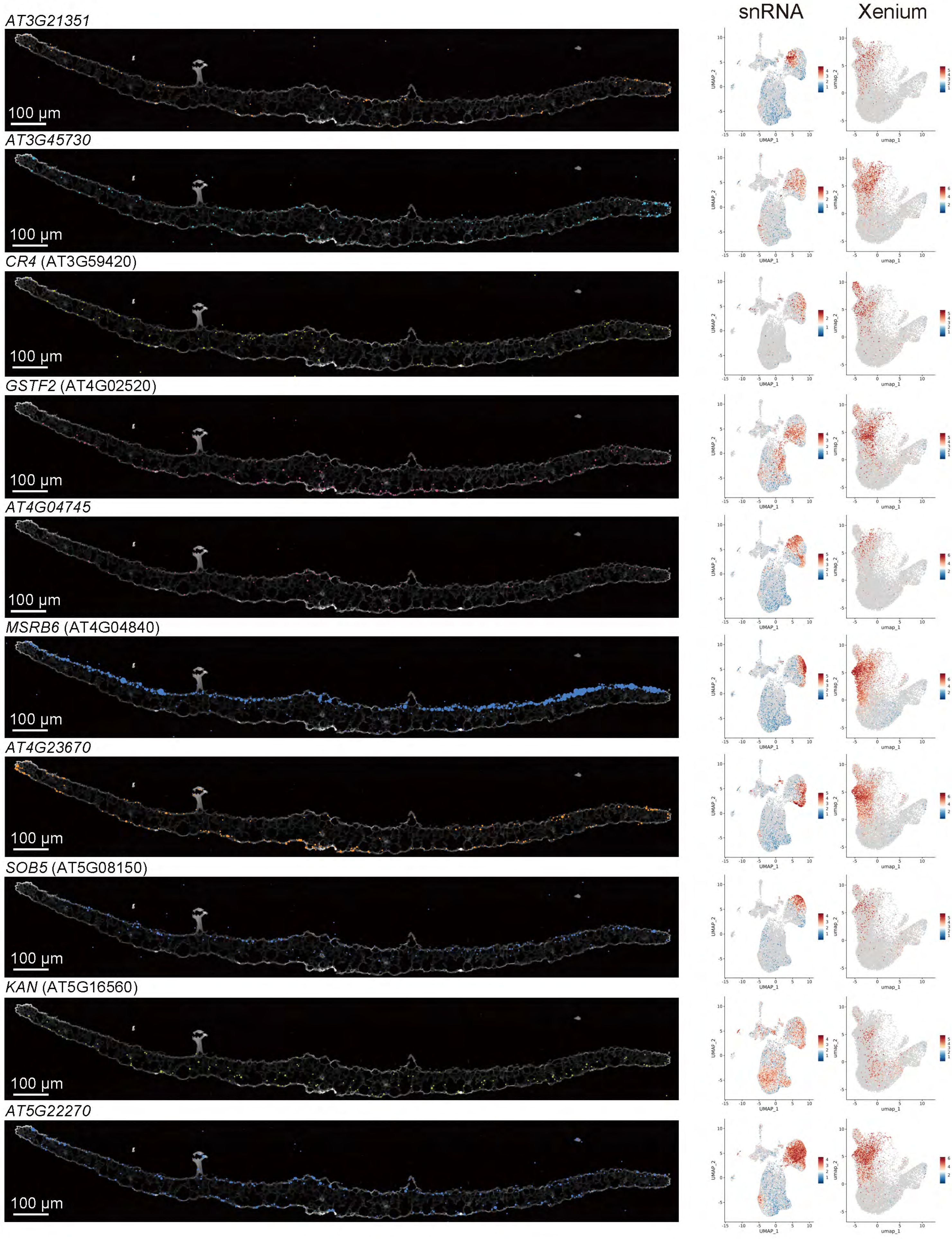
snRNA
Xenium
AT3G21351
AT3G45730
CR4 (AT3G59420)
GSTF2 (AT4G02520)
AT4G04745
MSRB6 (AT4G04840)
AT4G23670
SOB5 (AT5G08150)
KAN (AT5G16560)
AT5G22270
100 µm
UMAP_1
UMAP_2
umap_1
umap_2

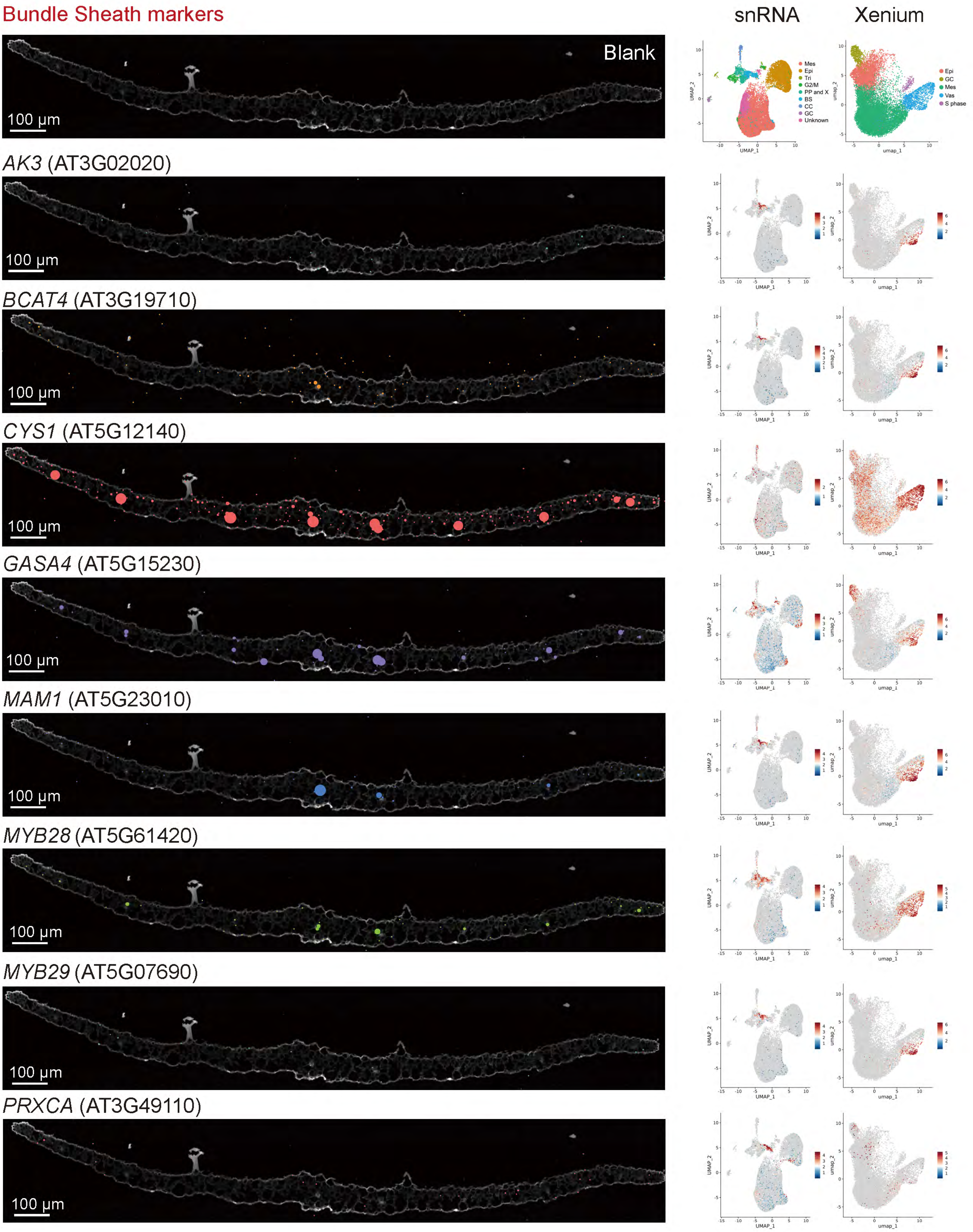

Bundle Sheath markers
snRNA
Xenium
Blank
100 µm
Mes
Epi
Tri
G2/M
PP and X
BS
CC
GC
Unknown
Epi
GC
Mes
Vas
S phase
UMAP_1
UMAP_2
umap_1
umap_2
*AK3* (AT3G02020)
*BCAT4* (AT3G19710)
*CYS1* (AT5G12140)
*GASA4* (AT5G15230)
*MAM1* (AT5G23010)
*MYB28* (AT5G61420)
*MYB29* (AT5G07690)
*PRXCA* (AT3G49110)

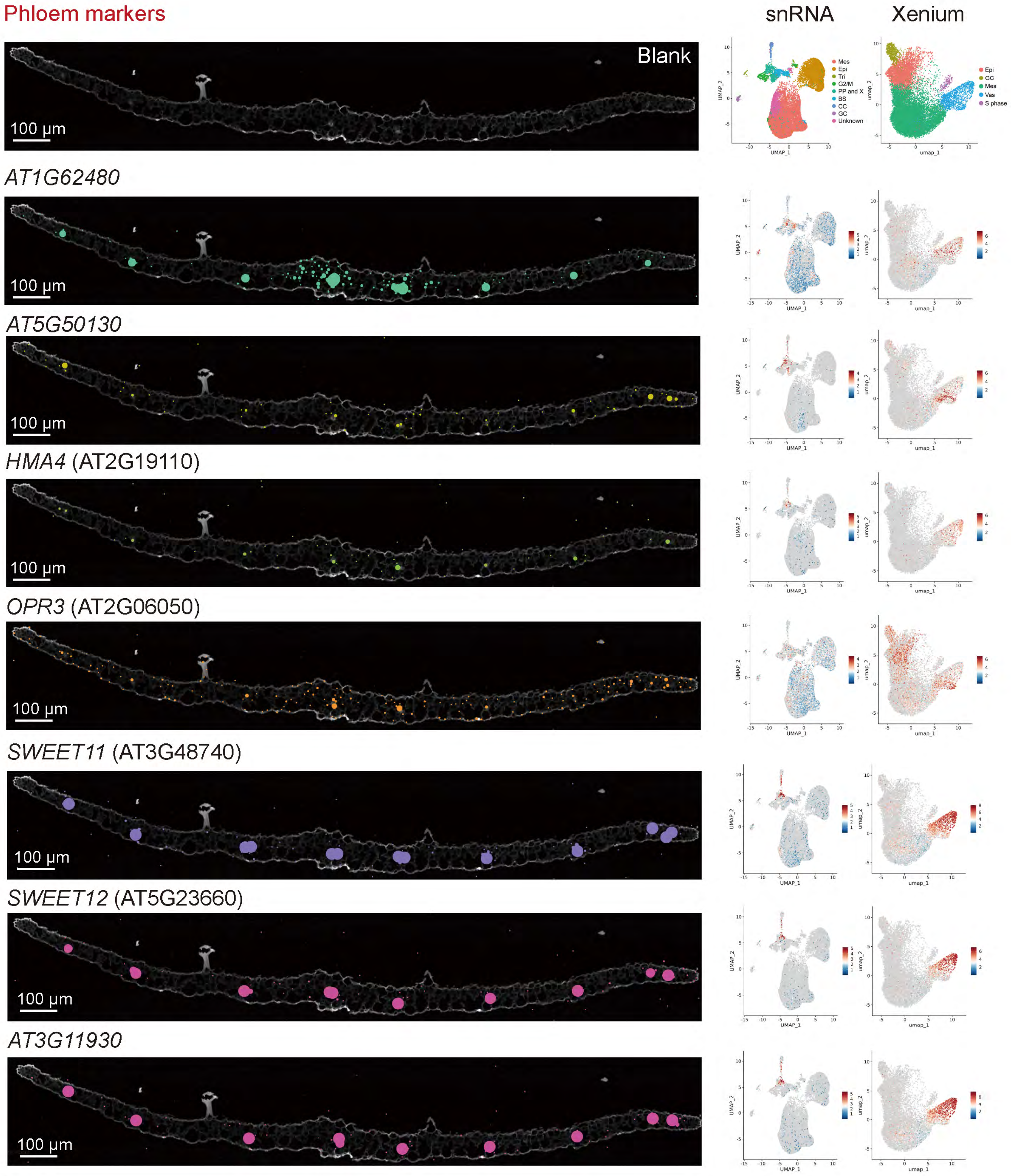

Phloem markers
snRNA
Xenium
Blank
100 µm
Mes
Epi
Tri
G2/M
PP and X
BS
CC
GC
Unknown
Epi
GC
Mes
Vas
S phase
UMAP_1
UMAP_2
umap_1
umap_2
AT1G62480
AT5G50130
HMA4 (AT2G19110)
OPR3 (AT2G06050)
SWEET11 (AT3G48740)
SWEET12 (AT5G23660)
AT3G11930

### Companion cell markers

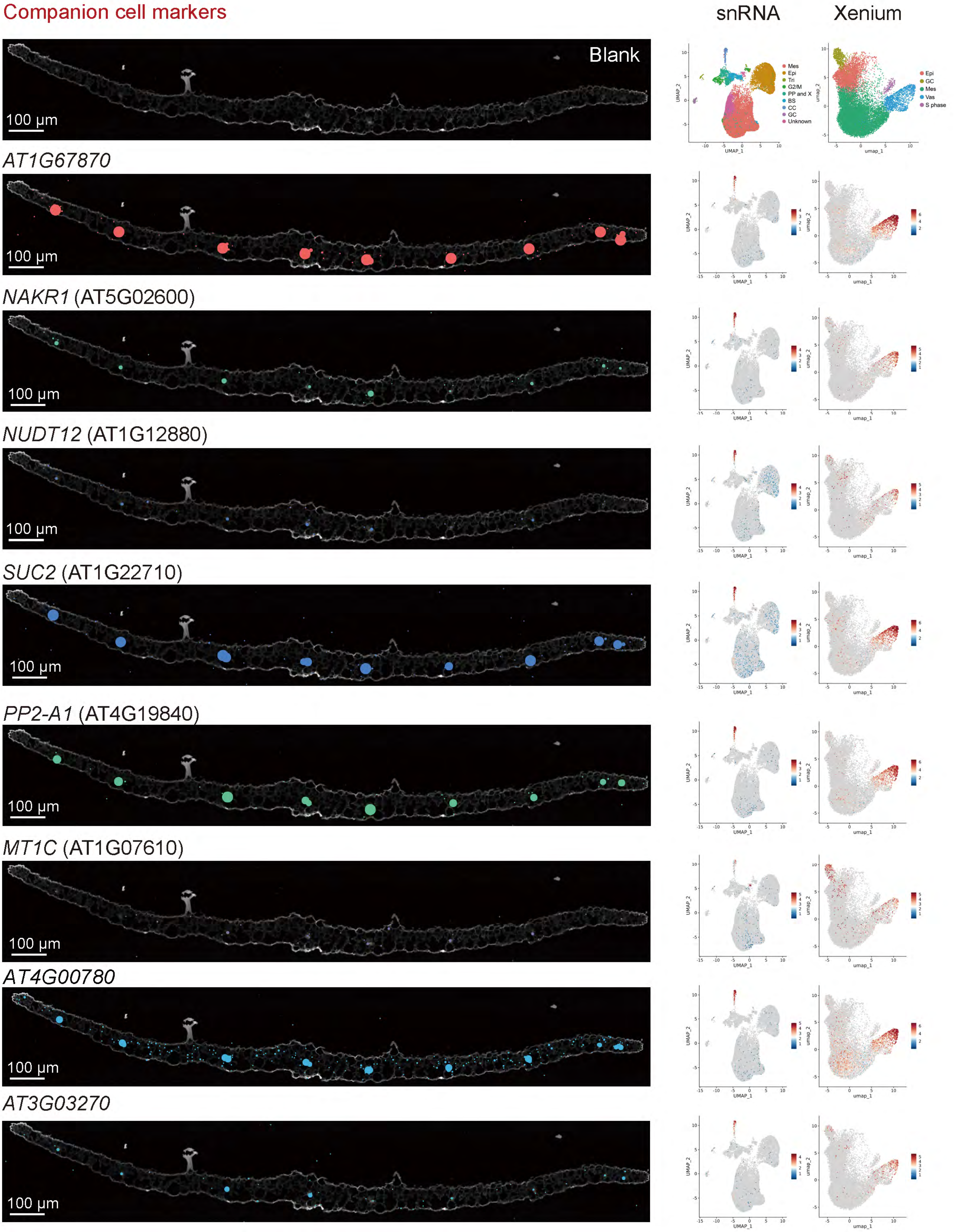

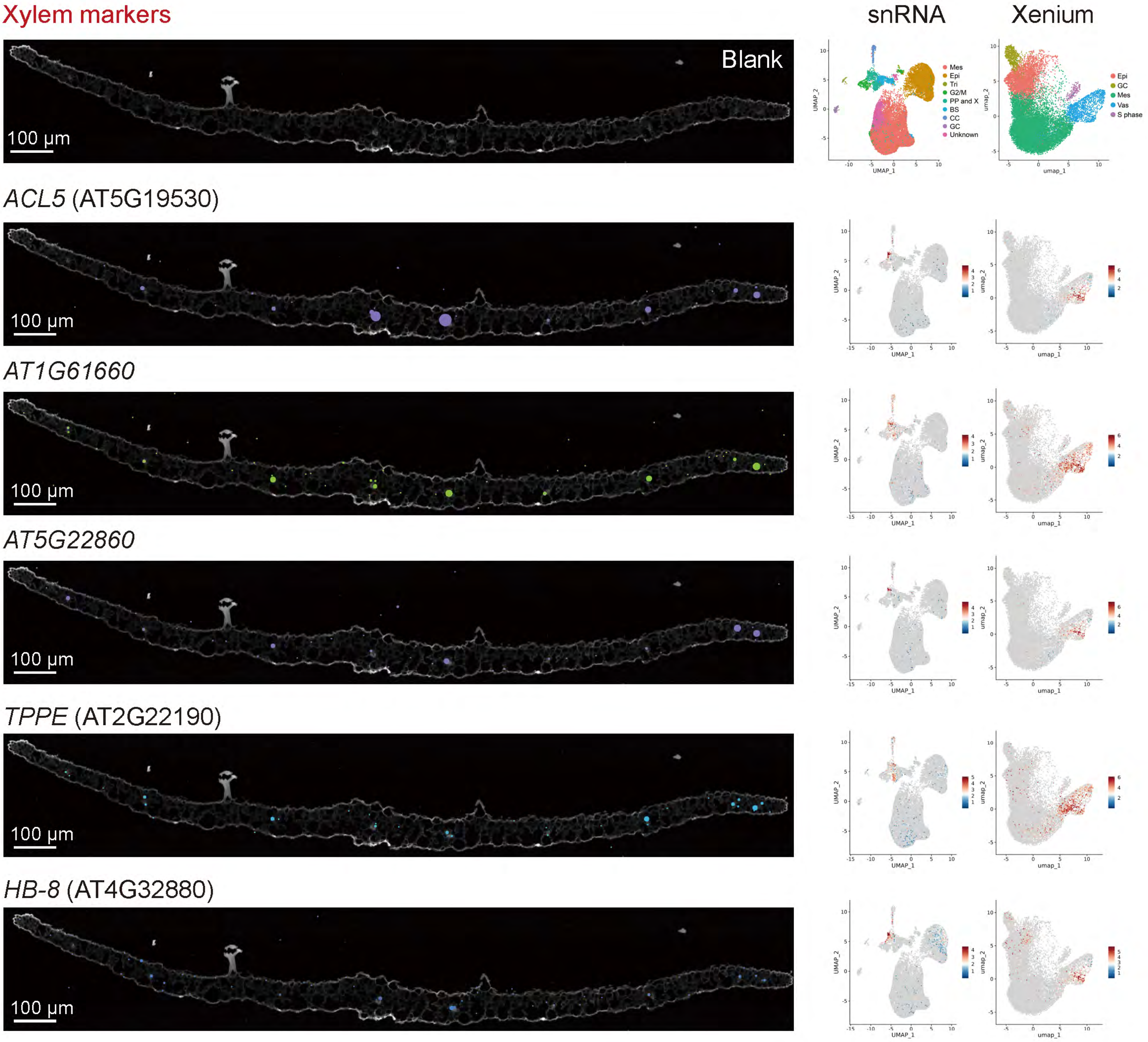
Xylem markers
snRNA
Xenium
Blank
100 µm
Mes
Epi
Tri
G2/M
PP and X
BS
CC
GC
Unknown
Epi
GC
Mes
Vas
S phase
UMAP_1
UMAP_2
umap_1
umap_2
ACL5 (AT5G19530)
AT1G61660
AT5G22860
TPPE (AT2G22190)
HB-8 (AT4G32880)

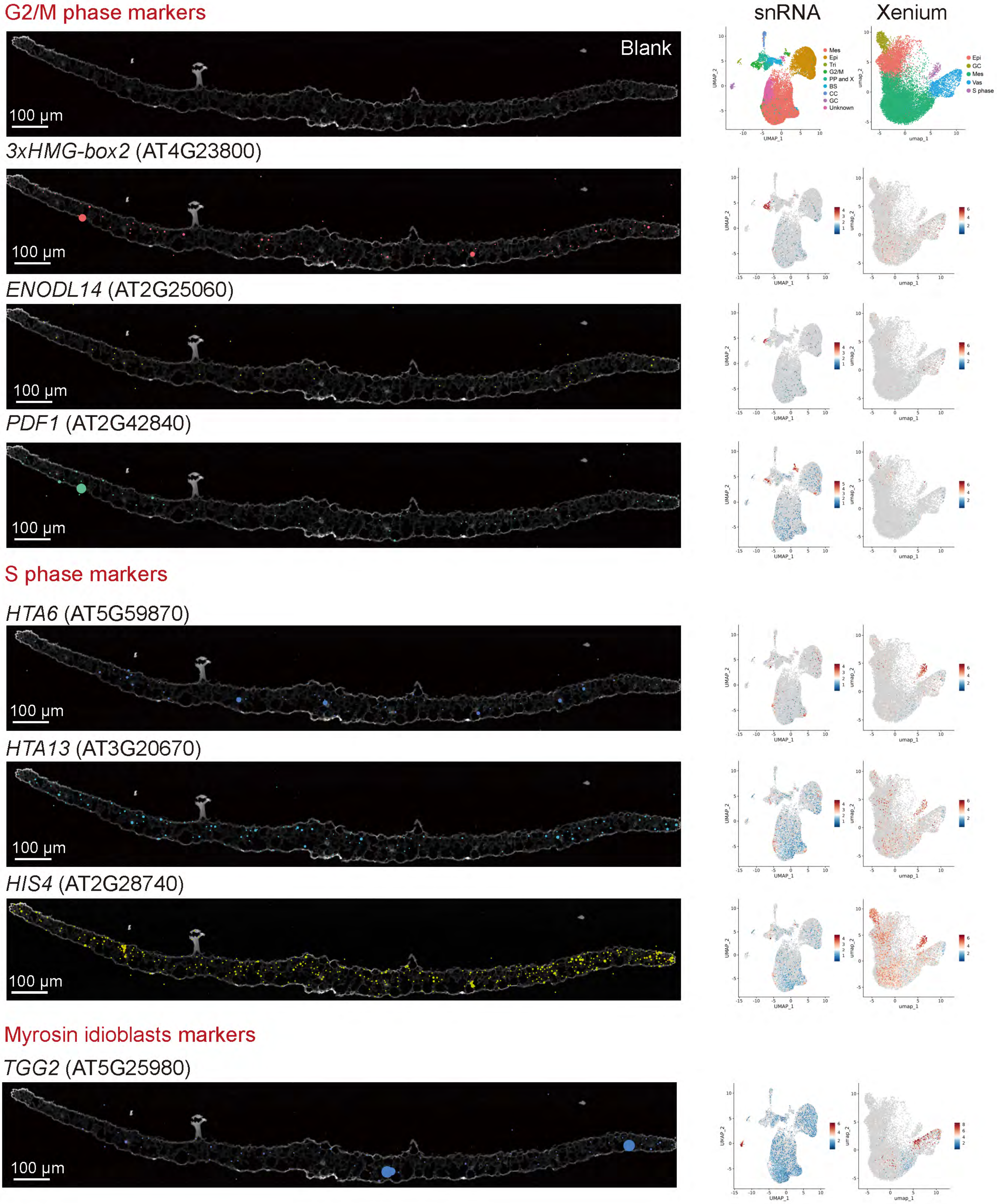

G2/M phase markers
snRNA
Xenium
Blank
100 µm
Mes
Epi
Tri
G2/M
PP and X
BS
CC
GC
Unknown
Epi
GC
Mes
Vas
S phase
UMAP_1
UMAP_2
umap_1
umap_2
3xHMG-box2 (AT4G23800)
ENODL14 (AT2G25060)
PDF1 (AT2G42840)
S phase markers
HTA6 (AT5G59870)
HTA13 (AT3G20670)
HIS4 (AT2G28740)
Myrosin idioblasts markers
TGG2 (AT5G25980)

**Supplemental Figure 1. 10x Xenium spatial validation of marker genes used for cell-type annotation**

10x Xenium *in situ* hybridization was used to validate marker genes for all major cell types and clusters identified in 14-day-old *Arabidopsis thaliana* seedling leaves after mock or antimycin A treatment. For each marker gene, fluorescence images show 10x Xenium probe signal in cross sections of Col-0 leaves, alongside feature plots of single-nucleus RNA-seq and 10x Xenium spatial UMAPs that display normalized expression.

Mesophyll identity is supported by markers including UDP-glycosyltransferase 85A1 (UGT85A1), Beta carbonic anhydrase 1 (CA1), Allene oxide cyclase 2 (AOC2), Beta carbonic anhydrase 2 (CA2), Suppressor of Yeast gpa1, Phosphate-regulated 81 and Xenotropic and Polytropic Retrovirus receptor family protein 1–like protein 1 (SPX1), Ribulose-1,5-bisphosphate carboxylase/oxygenase small subunit 2B (RBCS2B) and Early responsive to dehydration 1 (ERD1). Epidermal and leaf pavement cell identity is confirmed with markers such as Cysteine-rich receptor-like protein kinase 42 (CRK42), Lysine histidine transporter 1 (LHT1), Homeobox-leucine zipper protein 2 (HAT2), Beta-galactosidase 4 (BGAL4), MYB domain protein 96 (MYB96), Glycosylphosphatidylinositol-anchored lipid transfer protein 1 (LTPG1), Short hypocotyl 2/Auxin-responsive protein IAA3 (SHY2), YABBY domain protein 5 (YAB5), Lipid transfer protein 1 (LTP1), Beta-glucosidase PYK10 (PYK10) and Methionine sulfoxide reductase B6 (MSRB6).

### Specific markers

#### AT2G26250 Ketoacyl-CoA Synthase (KCS10/FDH)

Predicted : Epidermis

Experimental result : Epidermis

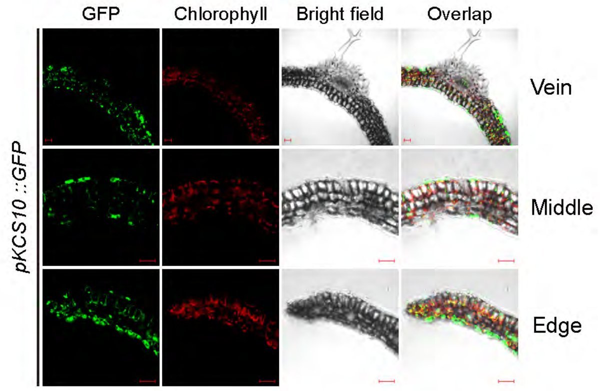


#### AT2G47240 LONG-CHAIN ACYL-COA SYNTHASE 1 (LACS1)

Predicted : Epidermis

Experimental result : Epidermis

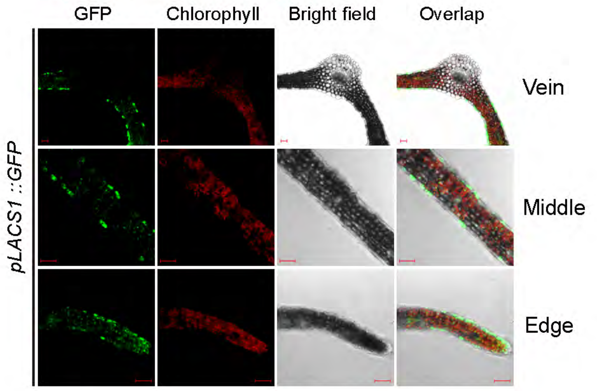


#### AT5G03760 CELLULOSE SYNTHASE LIKE A9 (CSLA9)

Predicted : Epidermis

Experimental result : Epidermis

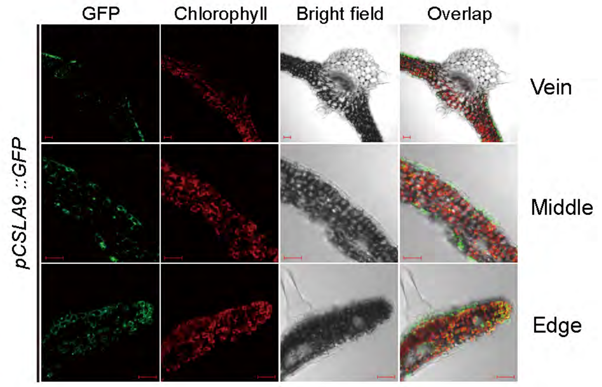


#### AT1G29670 GDSL-like Lipase/Acylhydrolase superfamily protein

Predicted : Epidermis

Experimental result : Epidermis

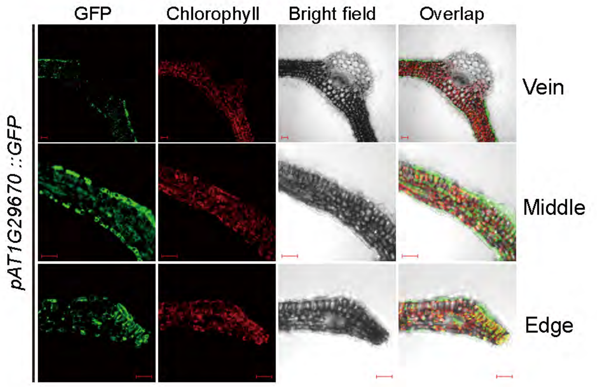


#### AT1G22400 UDP-GLUCOSYL TRANSFERASE 85A1 (UGT85A1)

Predicted : Unknown

Experimental result : Guard cell and epidermis

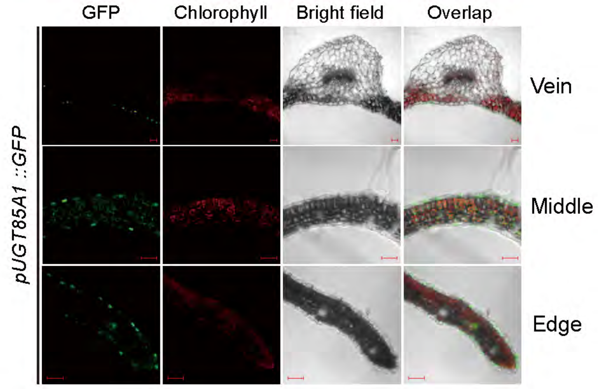


#### AT1G62300 WRKY family transcription factor 6 (WRKY6)

Predicted : Epidermis

Experimental result : Guard cell and epidermis

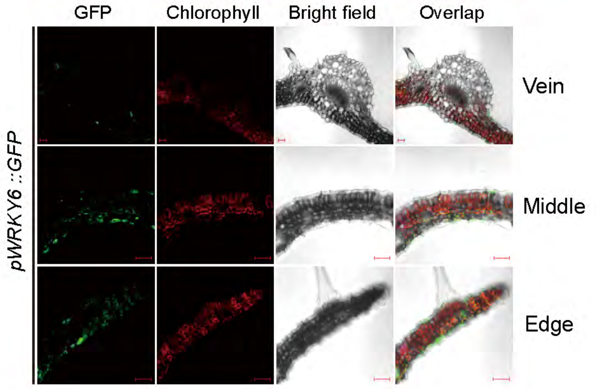


#### AT4G37870 phosphoenolpyruvate carboxykinase 1 (PCKA)

Predicted : Unknown

Experimental result : Guard cell and epidermis

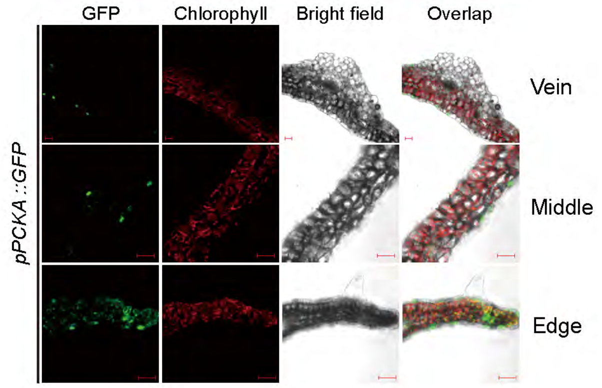


#### AT3G01500 BETA CARBONIC ANHYDRASE 1 (BCA1)

Predicted : Mesophyll

Experimental result : Mesophyll

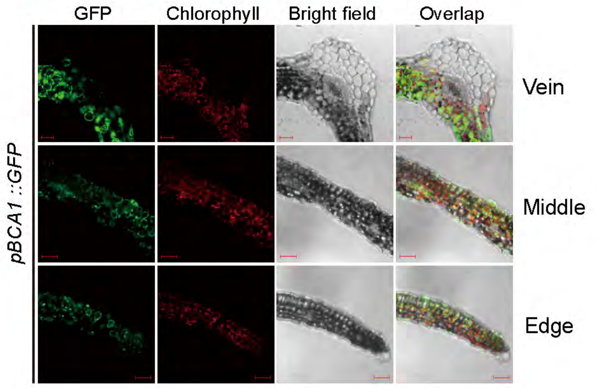

**AT5G59780 MYB DOMAIN PROTEIN 59 (MYB59)**

Predicted : Mesophyll

Experimental result : Mesophyll

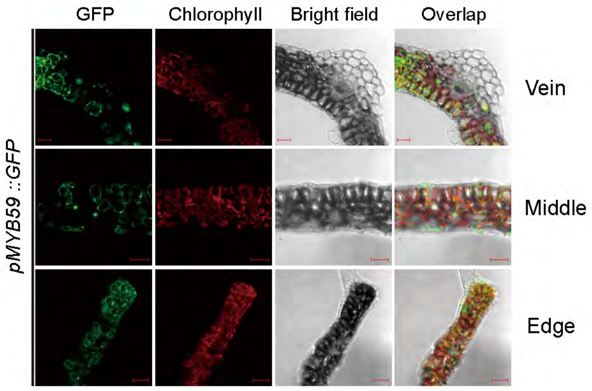

### Unspecific or low expression markers

#### AT2G24600 Ankyrin repeat family protein

Predicted : Mesophyll

Experimental result : Mainly mesophyll

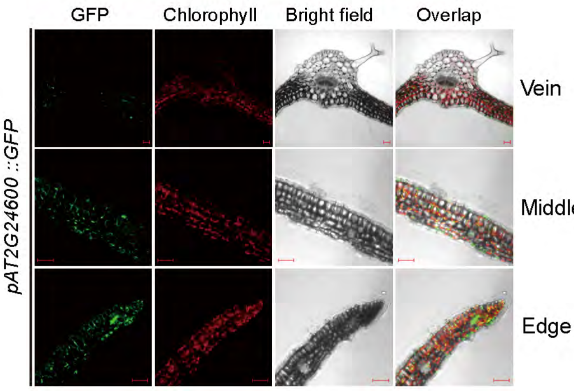


#### AT1G07610 metallothionein 1C (MT1C)

Predicted : Unknown

Experimental result : Mesophyll, epidermis and vascular

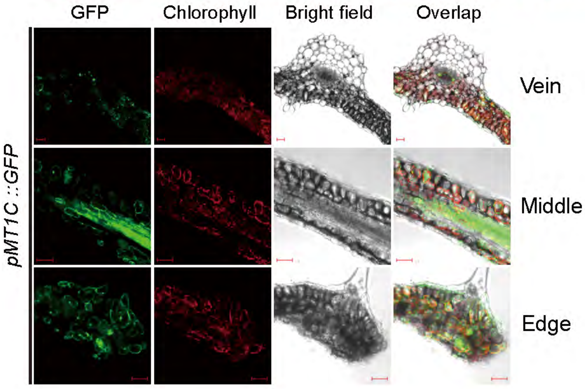


#### AT1G09932 Phosphoglycerate mutase family protein

Predicted : Phloem parenchyma

Experimental result :Mesophyll, epidermis and lateral vascular

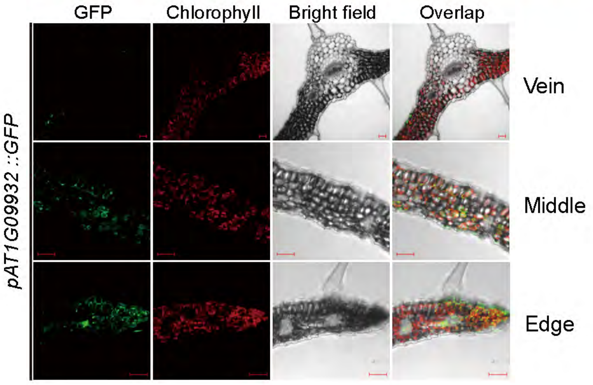


#### AT1G35140 Phosphate-responsive 1 family protein (EXL1)

Predicted : Epidermis

Experimental result : Mesophyll, epidermis and vascular

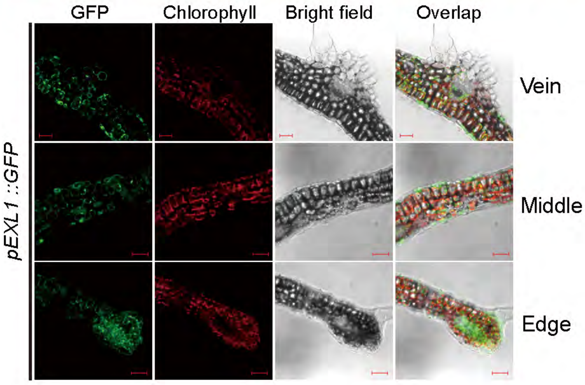


#### AT1G56060 cysteine-rich/transmembrane domain protein B

Predicted : Mesophyll

Experimental result : Mesophyll, epidermis and vascular

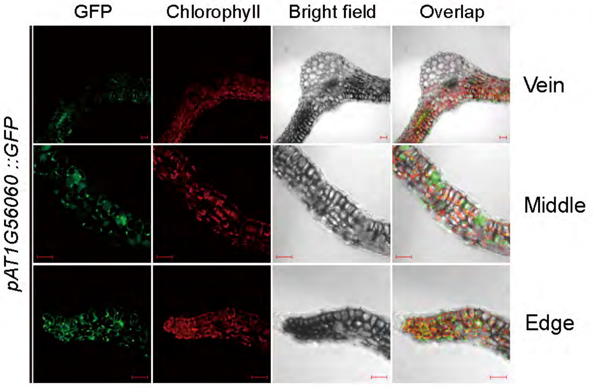


#### AT1G56650 MYELOBLASTOSIS PROTEIN 75 (MYB75)

Predicted : Unknown

Experimental result : Mesophyll, epidermis and vascular

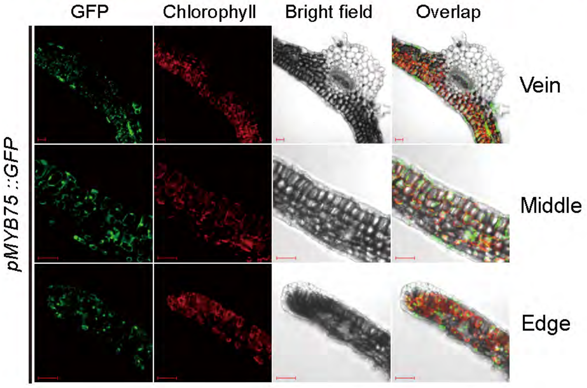


#### AT1G61660 basic helix-loop-helix (bHLH) DNA-binding superfamily protein 112 (BHLH112)

Predicted : Xylem

Experimental result : Mainly vascular and mesophyll

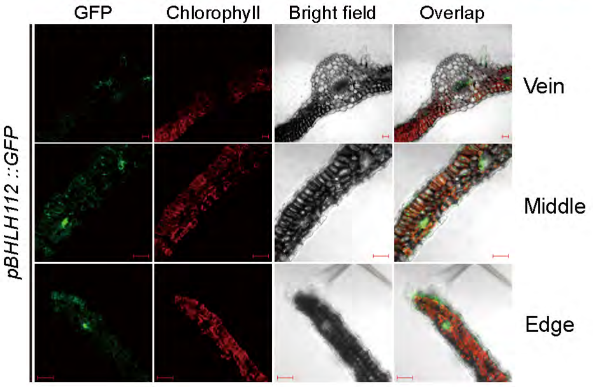


#### AT2G03760 sulfotransferase 12 (SOT12)

Predicted : Mesophyll

Experimental result : Epidermis and mesophyll

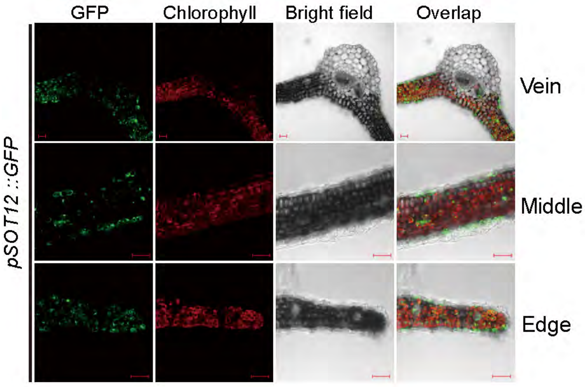

**AT2G04050 MATE efflux family protein (DTX3)**

Predicted : Unknown

Experimental result : Mesophyll, epidermis and vascular

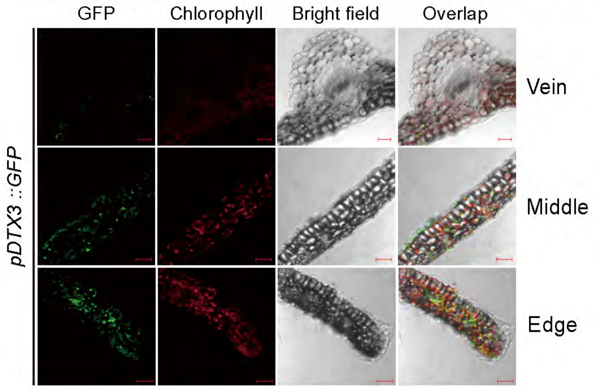


**AT2G11810 MONOGALACTOSYL DIACYLGLYCEROL SYNTHASE 3 (MGD3)**

Predicted : Mesophyll

Experimental result : Mesophyll and epidermis

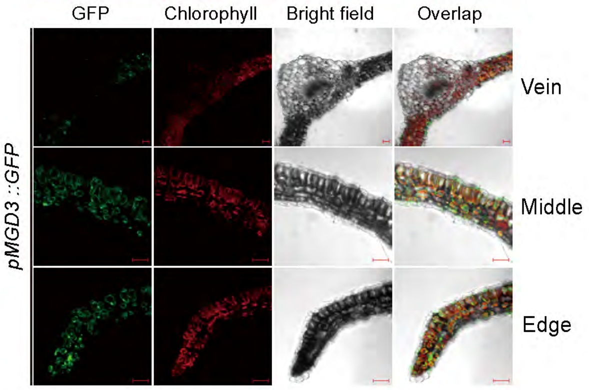


**AT2G19800 myo-inositol oxygenase 2 (MIOX2)**

Predicted : Epidermis

Experimental result :Mesophyll, epidermis and vascular

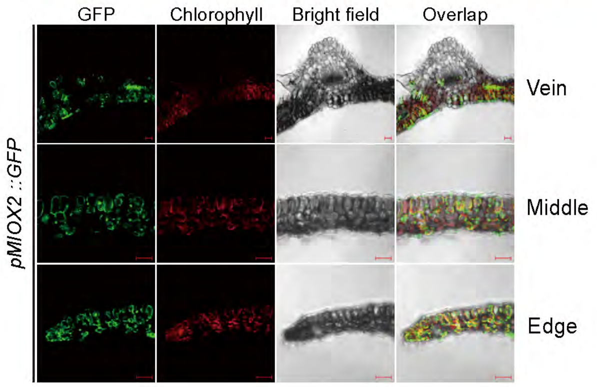


**AT2G28630 3-ketoacyl-CoA synthase 12 (KCS12)**

Predicted : Epidermis

Experimental result : Mesophyll, epidermis and vascular

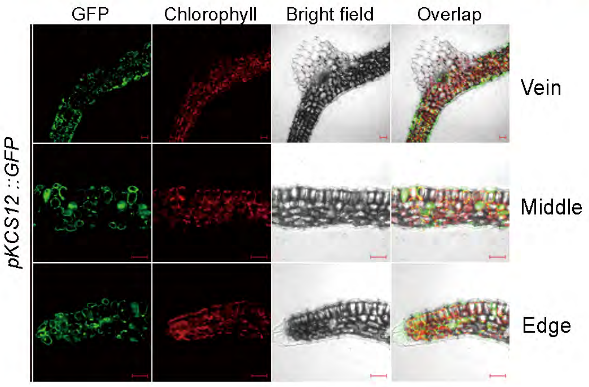


**AT2G38470 WRKY DNA-binding protein 33 (WRKY33)**

Predicted : Mesophyll

Experimental result : Mesophyll and epidermis

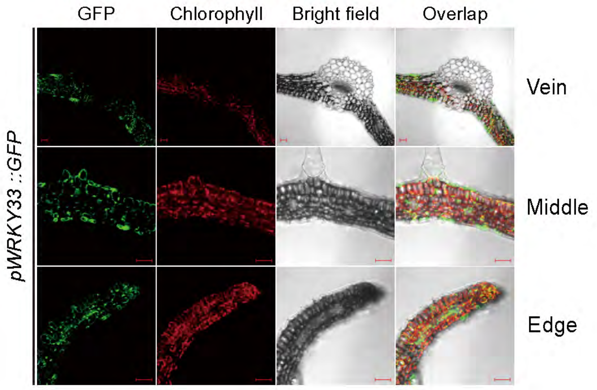


**AT2G38940 phosphate transporter 1;4 (PHT1-4)**

Predicted : Mesophyll

Experimental result : Mainly mesophyll and vascular

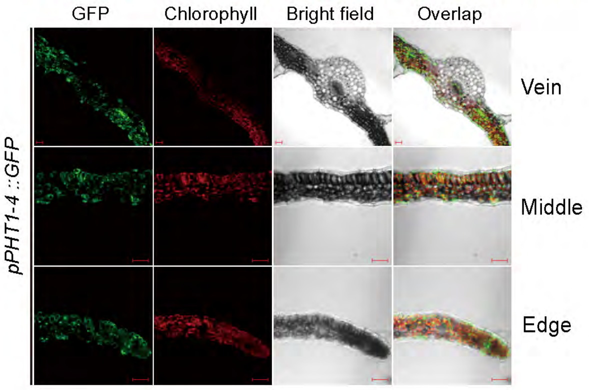


**AT3G01420 alpha-dioxygenase 1 (DOX1)**

Predicted : Unknown

Experimental result : Mesophyll, epidermis and vascular

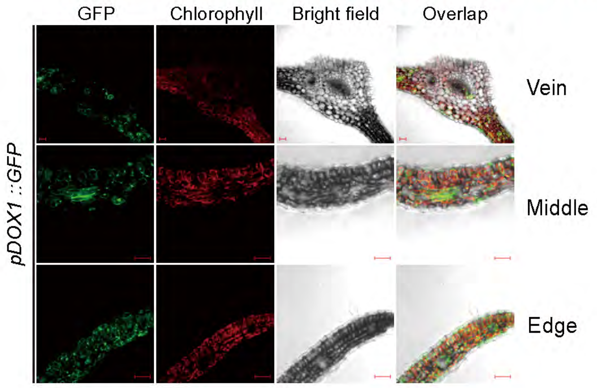


**AT3G15850 fatty acid desaturase 5 (ADS3)**

Predicted : Unknown

Experimental result : Epidermis and mesophyll

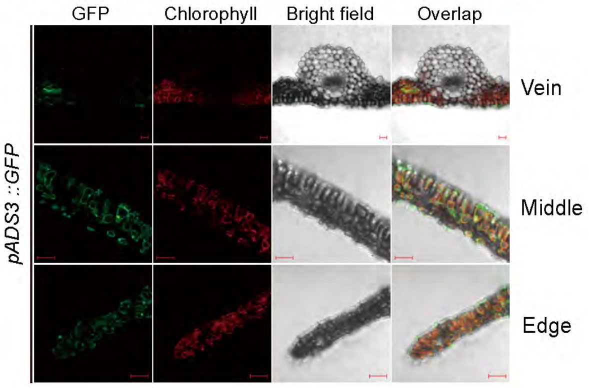

**AT3G19710 branched-chain aminotransferase4 (BCAT4)**

Predicted : Bundle Sheath

Experimental result : Mesophyll, epidermis and vascular

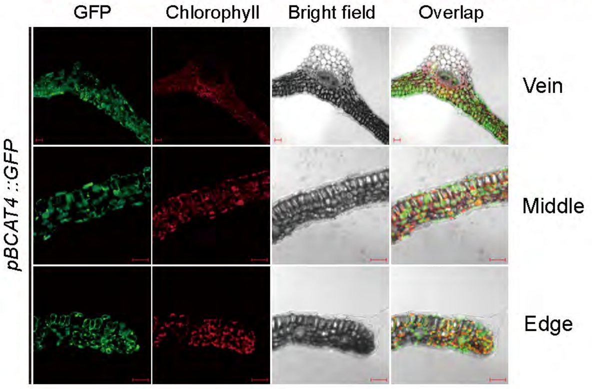


**AT3G23250 myb domain protein 15 (MYB15)**

Predicted : Phloem parenchyma

Experimental result : Mesophyll, epidermis and vascular

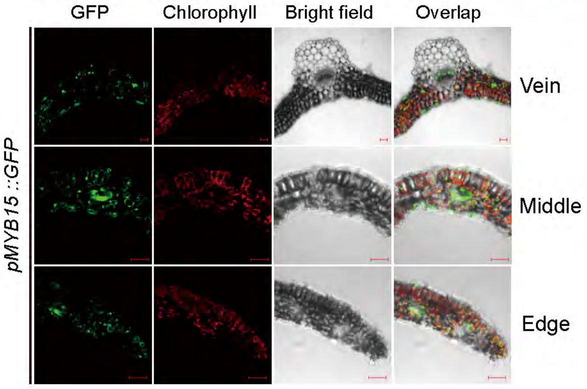


**AT3G45970 expansin-like A1 (EXLA1)**

Predicted : Epidermis

Experimental result :Mesophyll, epidermis and vascular

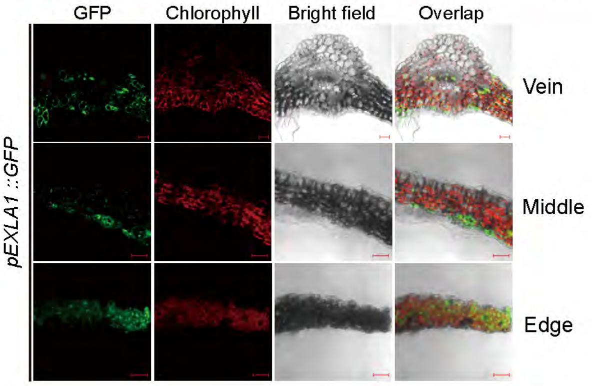


**AT3G46490 2-oxoglutarate (2OG) and Fe(II)-dependent oxygenase superfamily protein**

Predicted : Guard cell

Experimental result : Mesophyll, epidermis and vascular

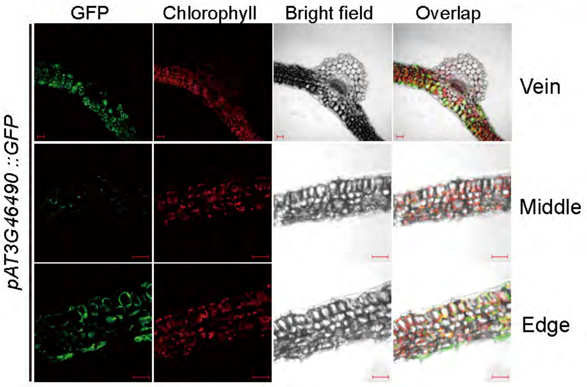


**AT3G52720 alpha carbonic anhydrase 1 (ACA1.1)**

Predicted : Unknown

Experimental result : Mesophyll, epidermis and vascular

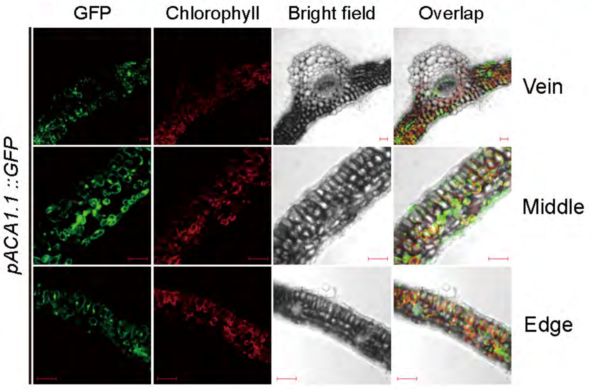


**AT3G54640 tryptophan synthase alpha chain (TSA1.1)**

Predicted : Bundle Sheath

Experimental result : Mesophyll, epidermis and vascular

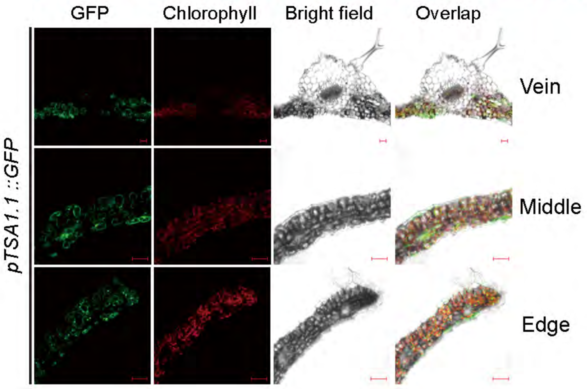


**AT4G01460 basic helix-loop-helix (bHLH) DNA-binding superfamily protein 57 (BHLH57)**

Predicted : Mesophyll

Experimental result : Mainly mesophyll

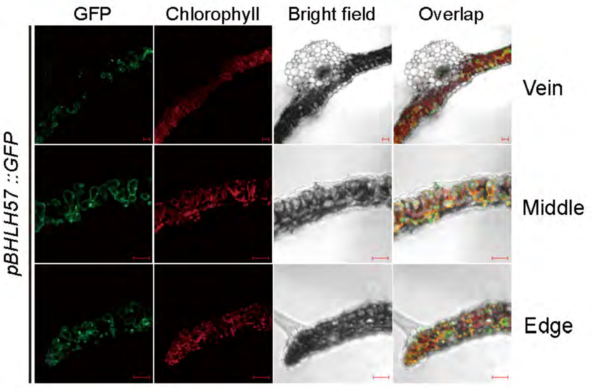


**AT4G01720 WRKY family transcription factor (WRKY47)**

Predicted : Mesophyll

Experimental result : Mesophyll, epidermis and vascular

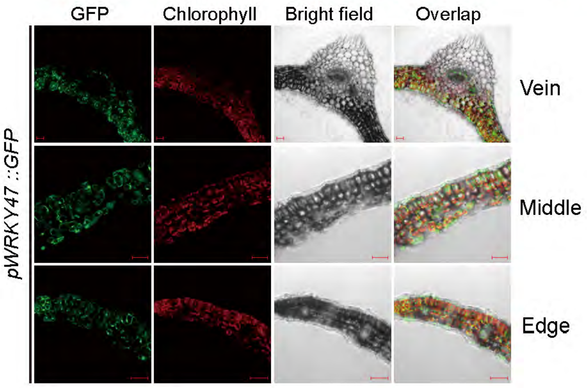

**AT4G03210 xyloglucan endotransglucosylase/hydrolase 9 (XTH9)**

Predicted : G2/M phase proliferating cell

Experimental result : Mesophyll, epidermis and vascular

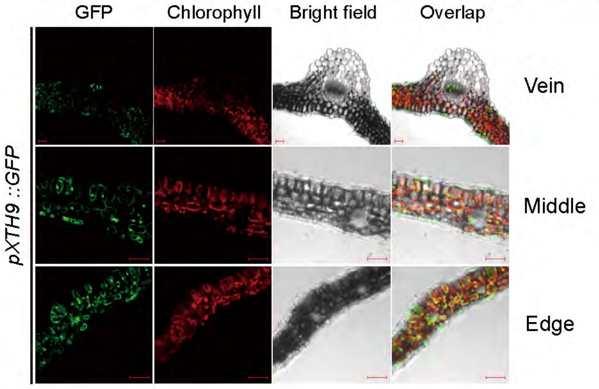


**AT4G23810 WRKY family transcription factor 53 (WRKY53)**

Predicted : Unknown

Experimental result : Mesophyll, epidermis and vascular

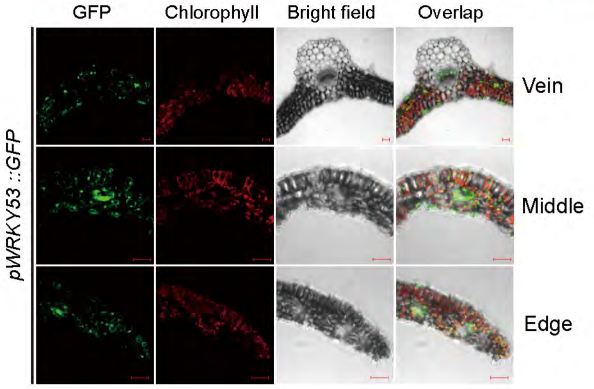


**AT4G25810 xyloglucan endotransglucosylase/hydrolase 23 (XTH23)**

Predicted : Unknown

Experimental result :Mesophyll, epidermis and vascular

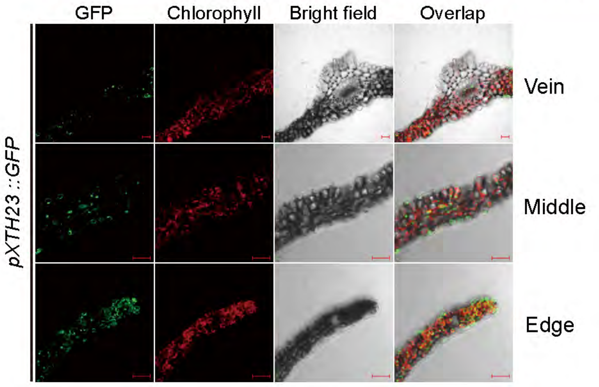


**AT5G07100 WRKY DNA-binding protein 26 (WRKY26)**

Predicted : Unknown

Experimental result : Mesophyll, epidermis and vascular

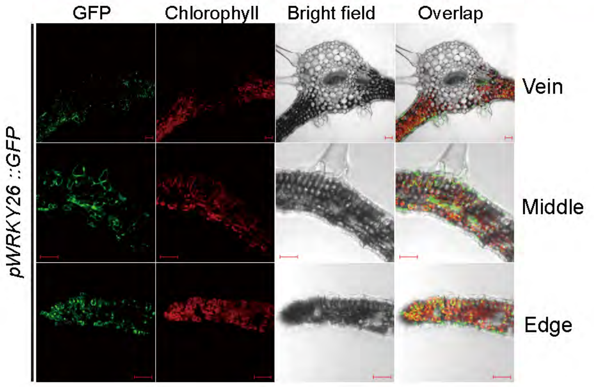


**AT5G16560 Homeodomain-like superfamily protein (KAN1)**

Predicted : Unknown

Experimental result : Mesophyll, epidermis and vascular

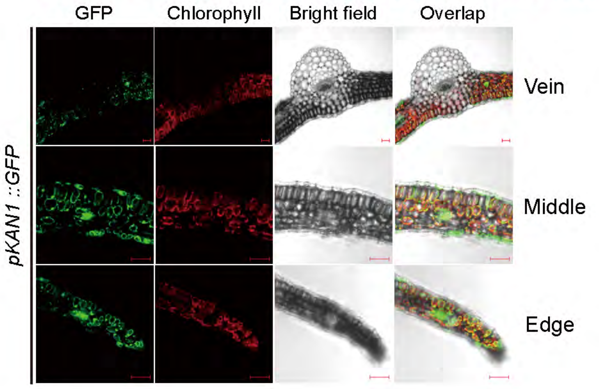


**AT5G22580 Stress responsive A/B BarrelDomain-containing protein**

Predicted : Unknown

Experimental result : Mesophyll, epidermis and vascular

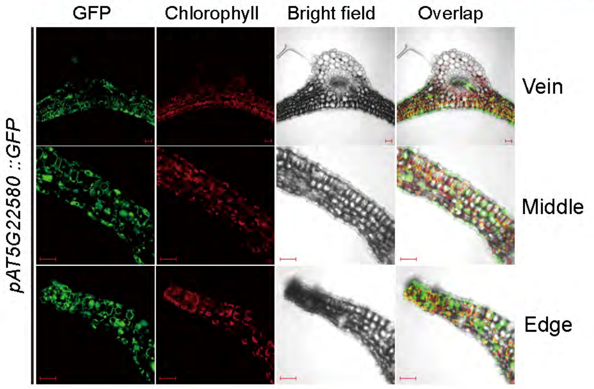


**AT5G48570 FKBP-type peptidyl-prolyl cis-trans isomerase family protein (FKBP65)**

Predicted : Phloem parenchyma

Experimental result : Mesophyll, epidermis and vascular

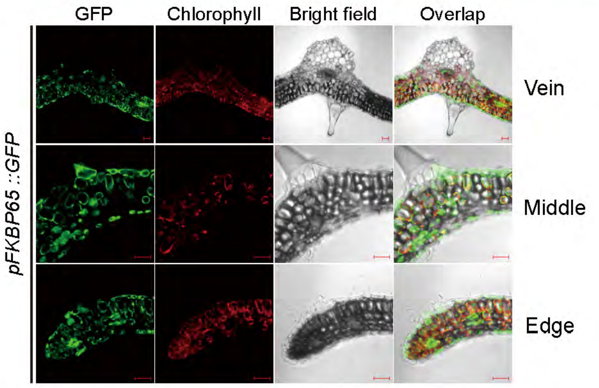


**AT5G48850 SULPHUR DEFICIENCY-INDUCED 1 (SDI1)**

Predicted : Unknown

Experimental result : Mesophyll, epidermis and vascular

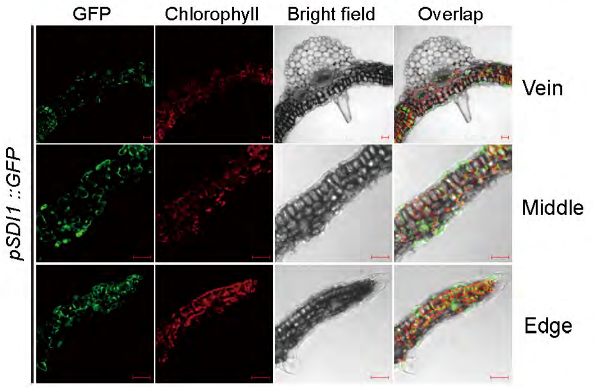

**AT5G62480 glutathione S-transferase tau 9 (GSTU9)**

Predicted : Unknown

Experimental result : Mesophyll, epidermis and vascular

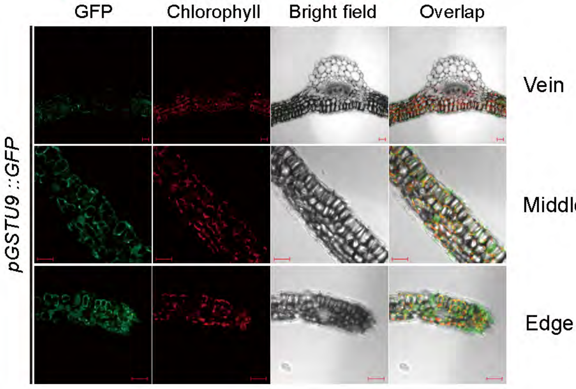


**AT5G49520 WRKY DNA-binding protein 48 (WRKY48)**

Predicted : Unknown

Experimental result : Mesophyll and epidermis

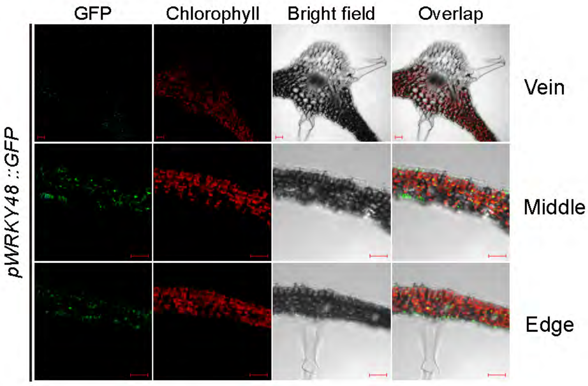


**AT2G28815 50S ribosomal protein L16**

Predicted : Unknown

Experimental result : Unknown

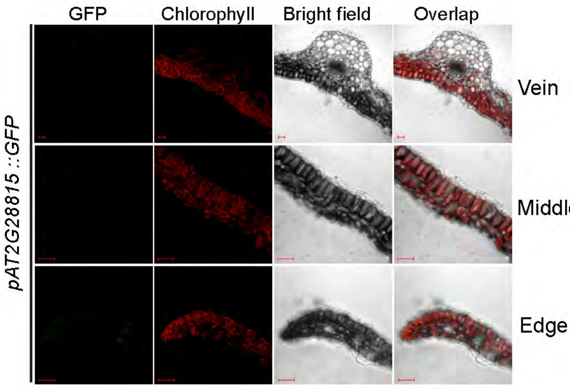


**AT2G19190 FLG22-induced receptor-like kinase 1 (SIRK)**

Predicted : Mesophyll

Experimental result : Mesophyll and abaxial epidermis of middle vein

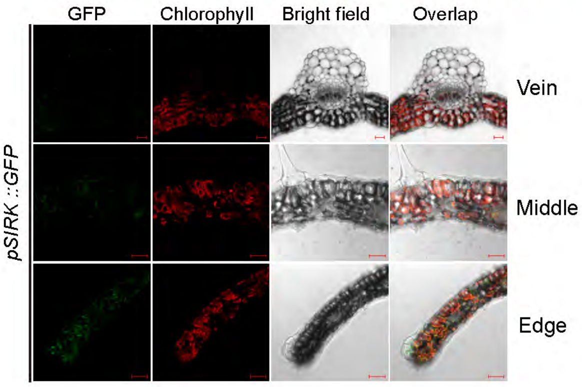


**AT2G20720 Pentatricopeptide repeat (PPR) superfamily protein**

Predicted : Unknown

Experimental result : Unknown

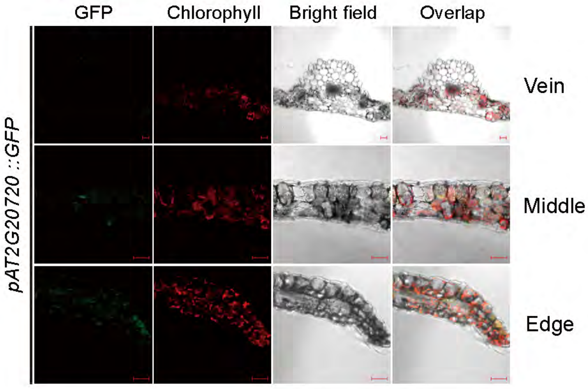


**AT1G05680 Uridine diphosphate glycosyltransferase 74E2 (UGT74E2)**

Predicted : Unknown

Experimental result : Unknown

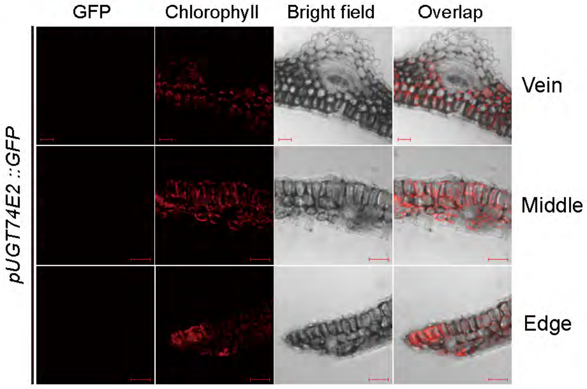


**AT3G28740 Cytochrome P450 superfamily protein (CYP81D11)**

Predicted : Mesophyll

Experimental result : Unknown

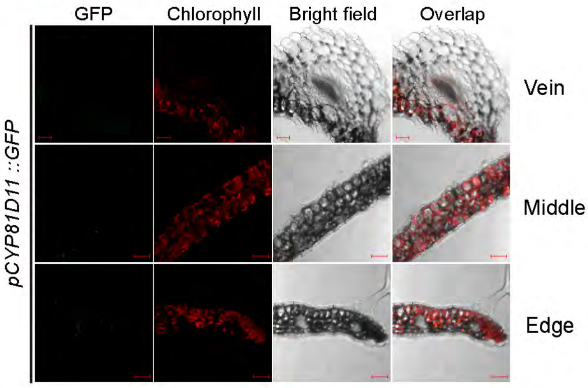


**AT5G24530 DOWNY MILDEW RESISTANT 6 (DMR6)**

Predicted : Mesophyll

Experimental result : Unknown

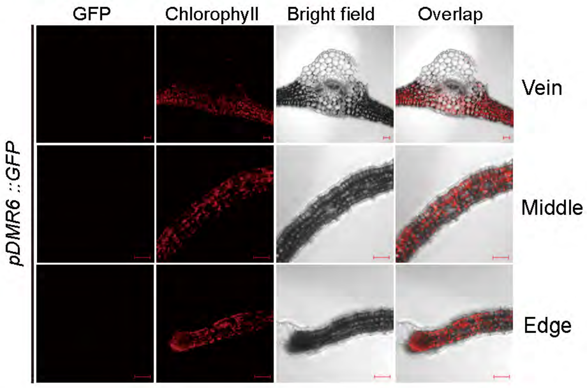

**AT1G25250 indeterminate(ID)-domain 16 (AtIDD16)**

Predicted : Mesophyll

Experimental result : Unknown

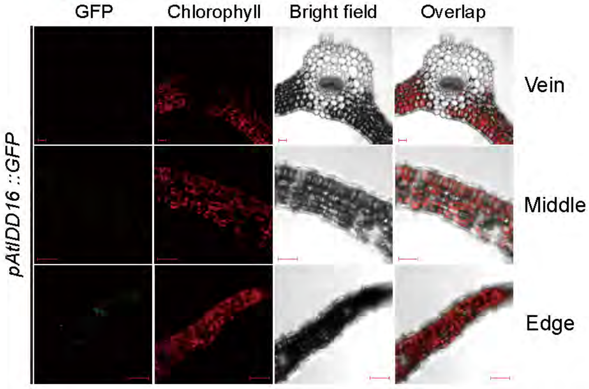


**AT5G61420 myb domain protein 28 (MYB28)**

Predicted : Bundle sheath

Experimental result : Unknown

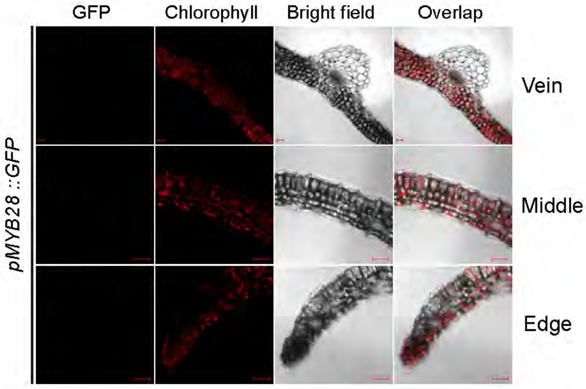


**AT5G44572 SERINE RICH ENDOGENOUS PEPTIDE 6 (PROSCOOP6)**

Predicted : Unknown

Experimental result : Mesophyll and abaxial epidermis of middle vein

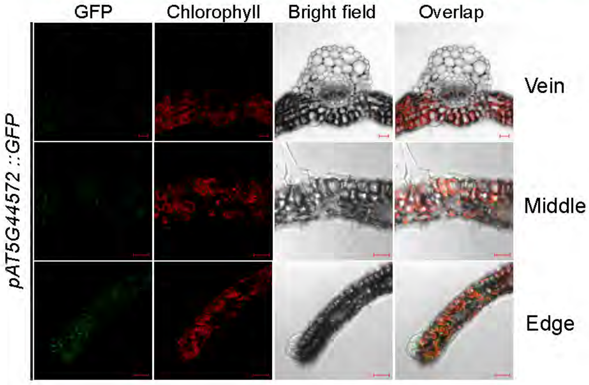


**Supplemental Figure 2. Confirmation of Cell markers using Promoter GFP tagging.** Transverse sections of leaves expressing promoter::GFP fusions for 52 candidate marker genes are shown as GFP, chlorophyll autofluorescence, bright-field and merged images at the mid-vein, lamina middle and edge. Predicted cell-type enrichment based on single-nucleus and spatial transcriptome data is indicated above each panel, and the experimentally observed pattern is indicated below. Guard-cell, epidermal and mesophyll markers were successfully validated for the first 9 genes. The remaining 43 reporters showed broad, weak or ambiguous expression across tissues and therefore did not confirm specific or enhanced expression in the predicted cell types.

AT1G04040 (Epidermis marker， marker of cluster 8 in integrated UMAP)
Scale Bar = 100 μm.

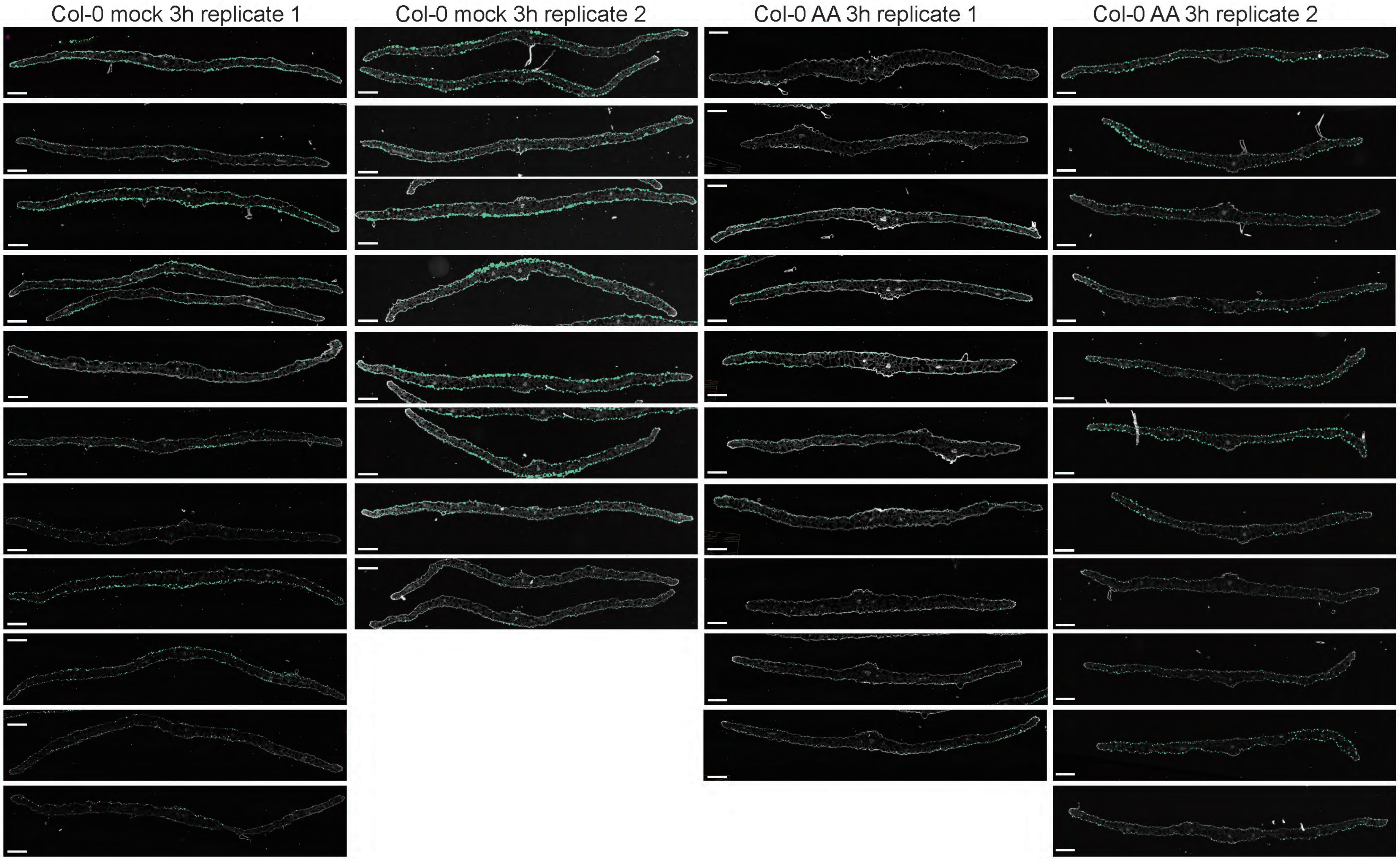

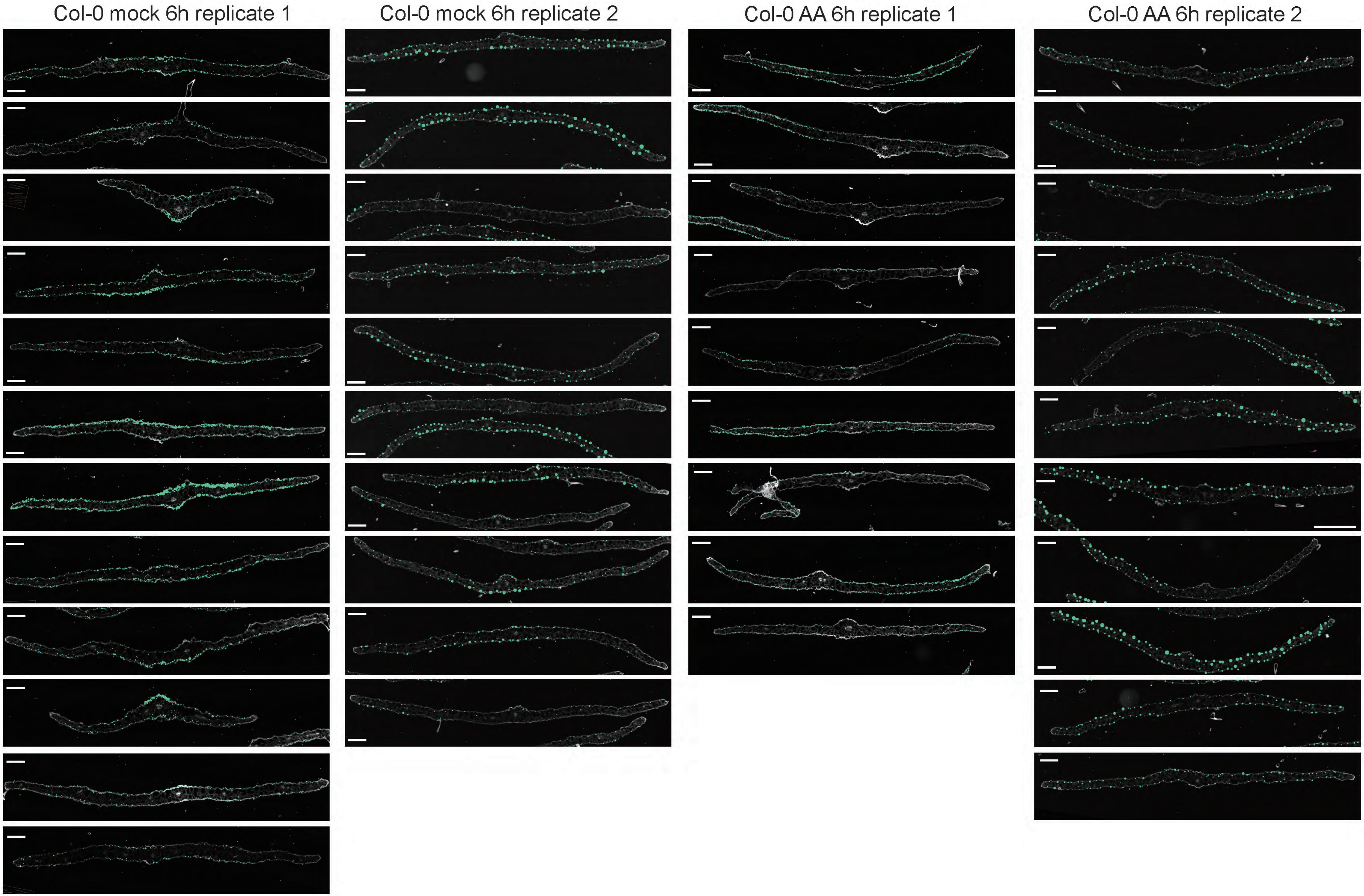

Col-0 mock 6h replicate 1
Col-0 mock 6h replicate 2
Col-0 AA 6h replicate 1
Col-0 AA 6h replicate 2

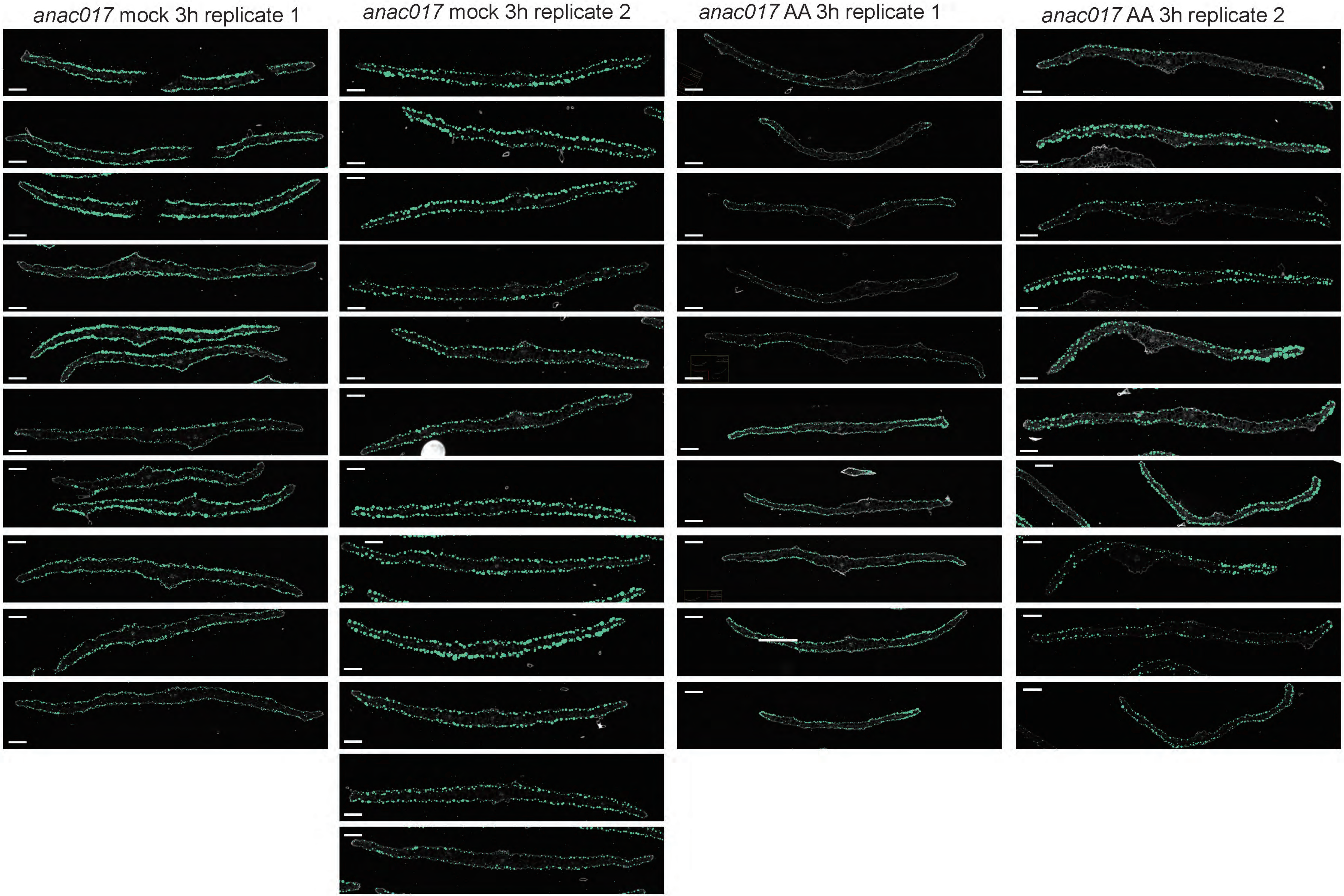
*anac017* mock 3h replicate 1
*anac017* mock 3h replicate 2
*anac017* AA 3h replicate 1
*anac017* AA 3h replicate 2

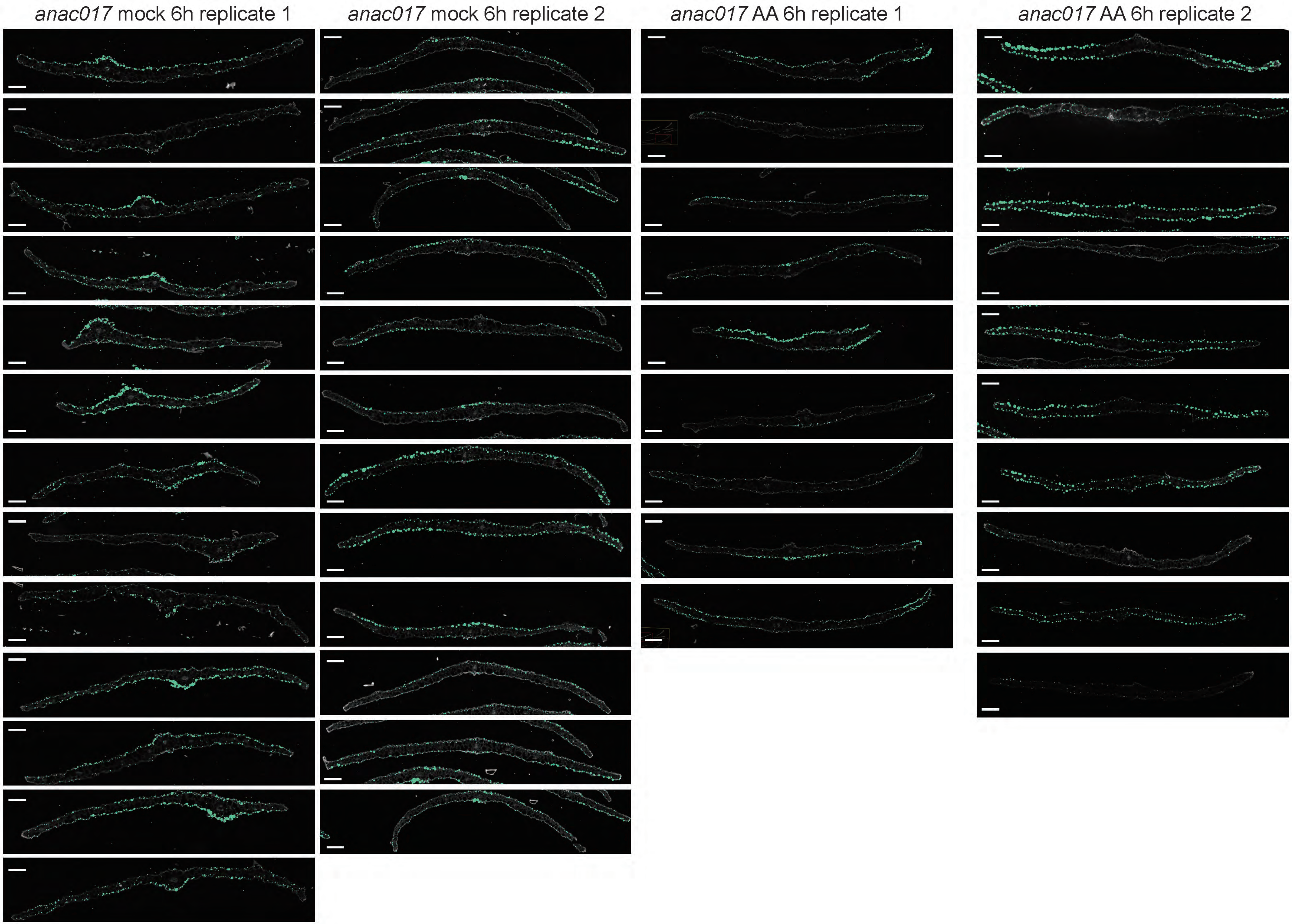

*anac017* mock 6h replicate 1
*anac017* mock 6h replicate 2
*anac017* AA 6h replicate 1
*anac017* AA 6h replicate 2

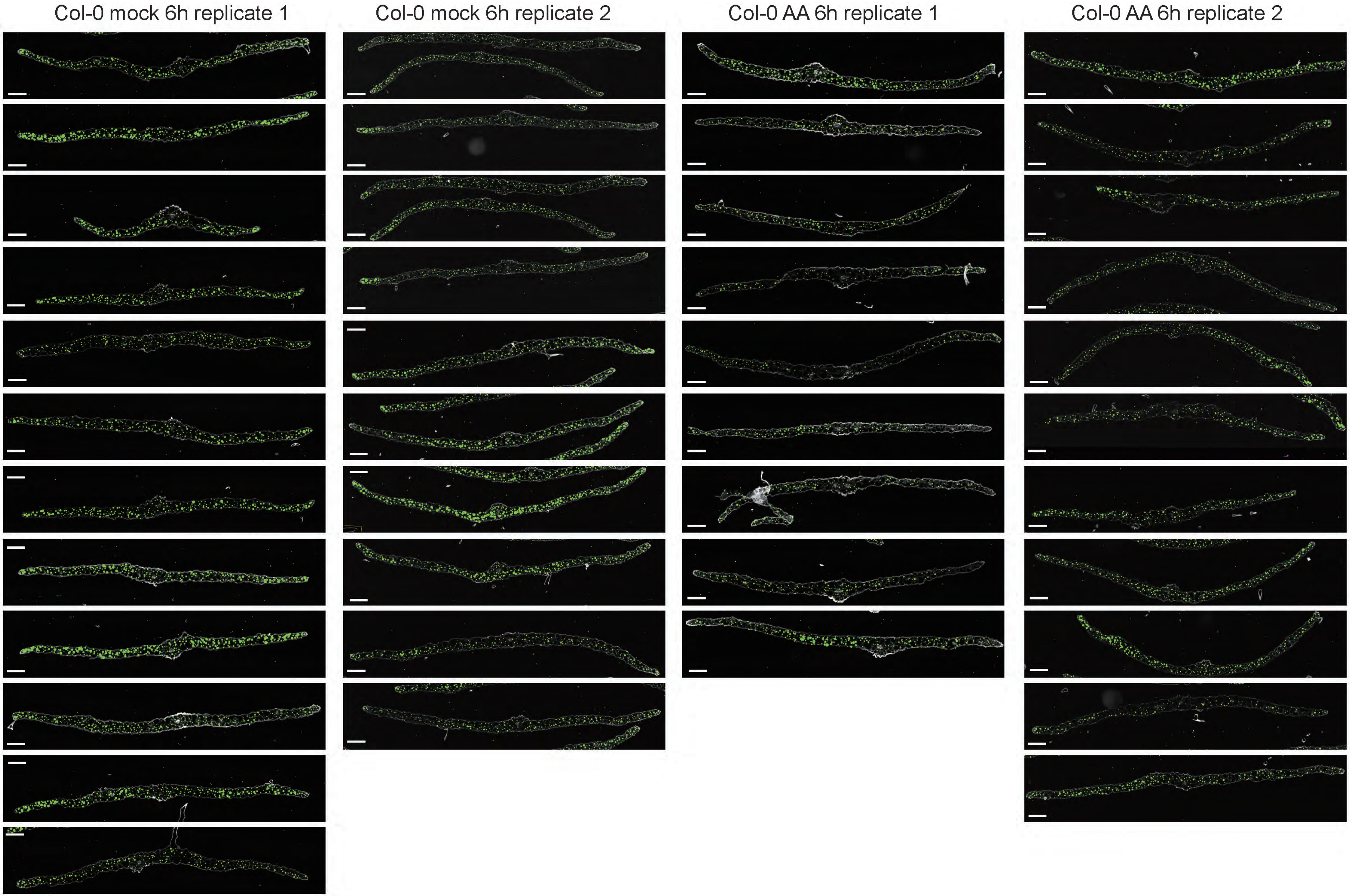

Col-0 mock 6h replicate 1
Col-0 mock 6h replicate 2
Col-0 AA 6h replicate 1
Col-0 AA 6h replicate 2

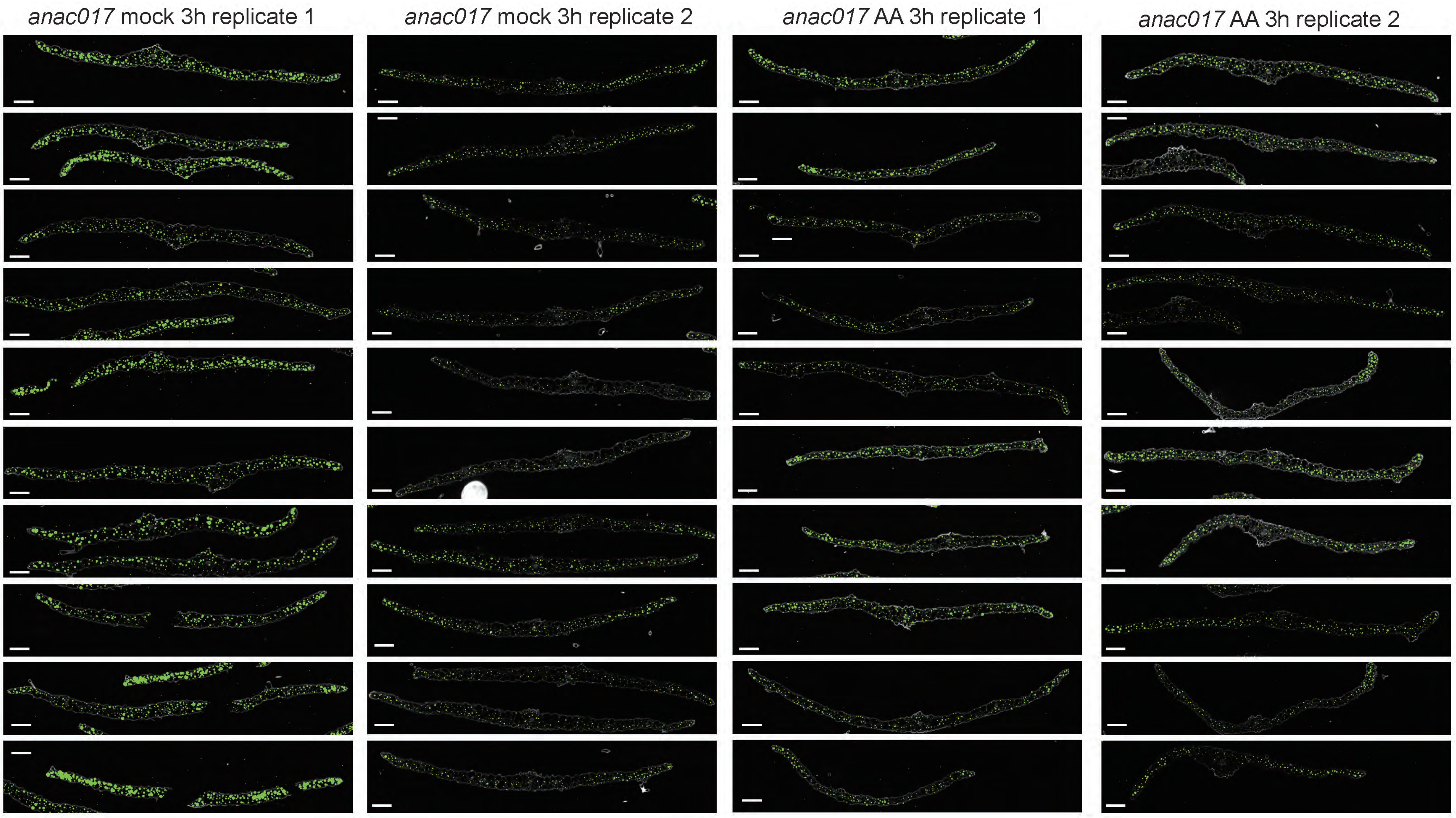
*anac017* mock 3h replicate 1
*anac017* mock 3h replicate 2
*anac017* AA 3h replicate 1
*anac017* AA 3h replicate 2

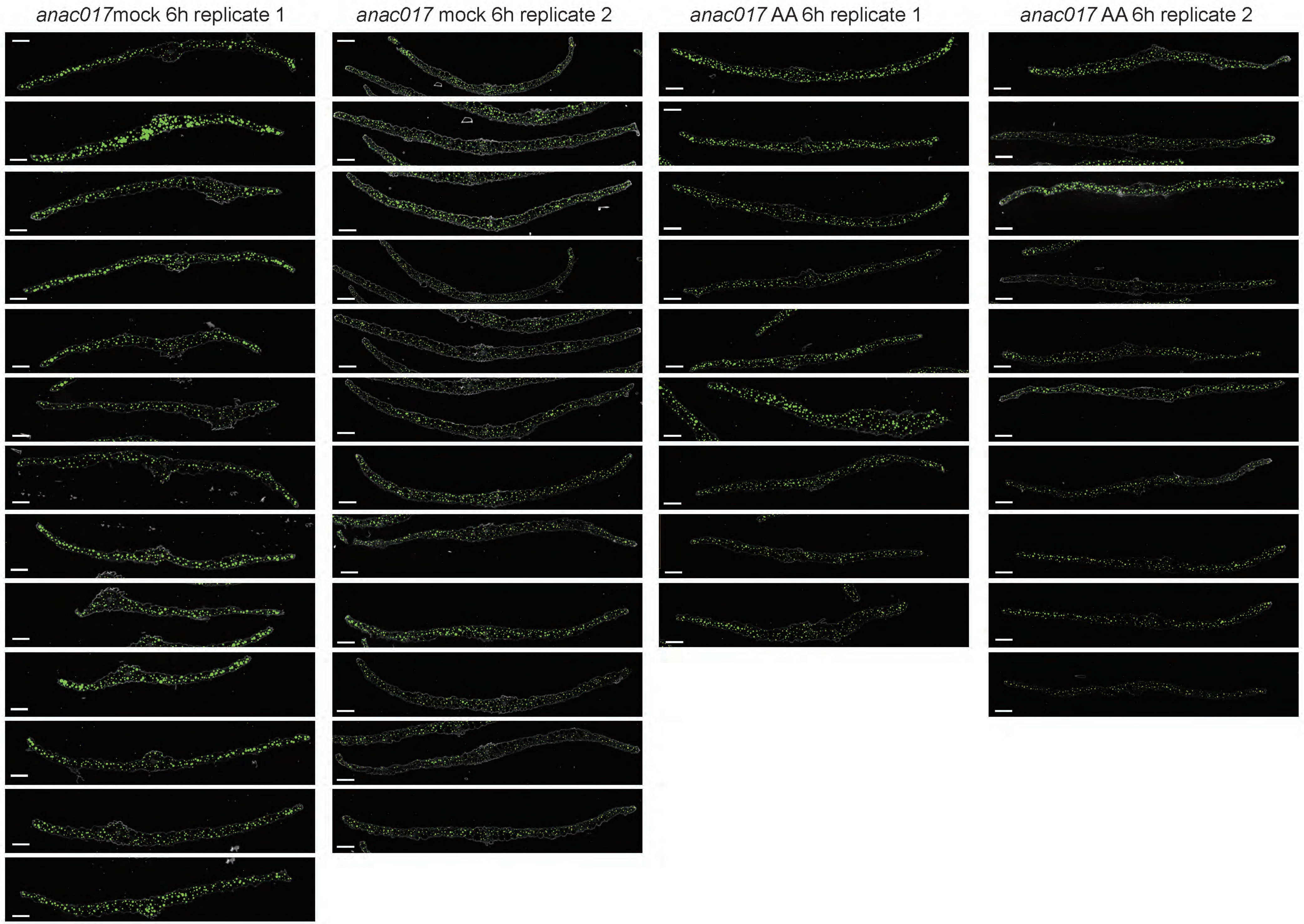
*anac017*mock 6h replicate 1
*anac017* mock 6h replicate 2
*anac017* AA 6h replicate 1
*anac017* AA 6h replicate 2

TIR (AT1G72930, predicted Mesophyl marker, marker of cluster 4 in integrated UMAP
Bar = 100 µm.

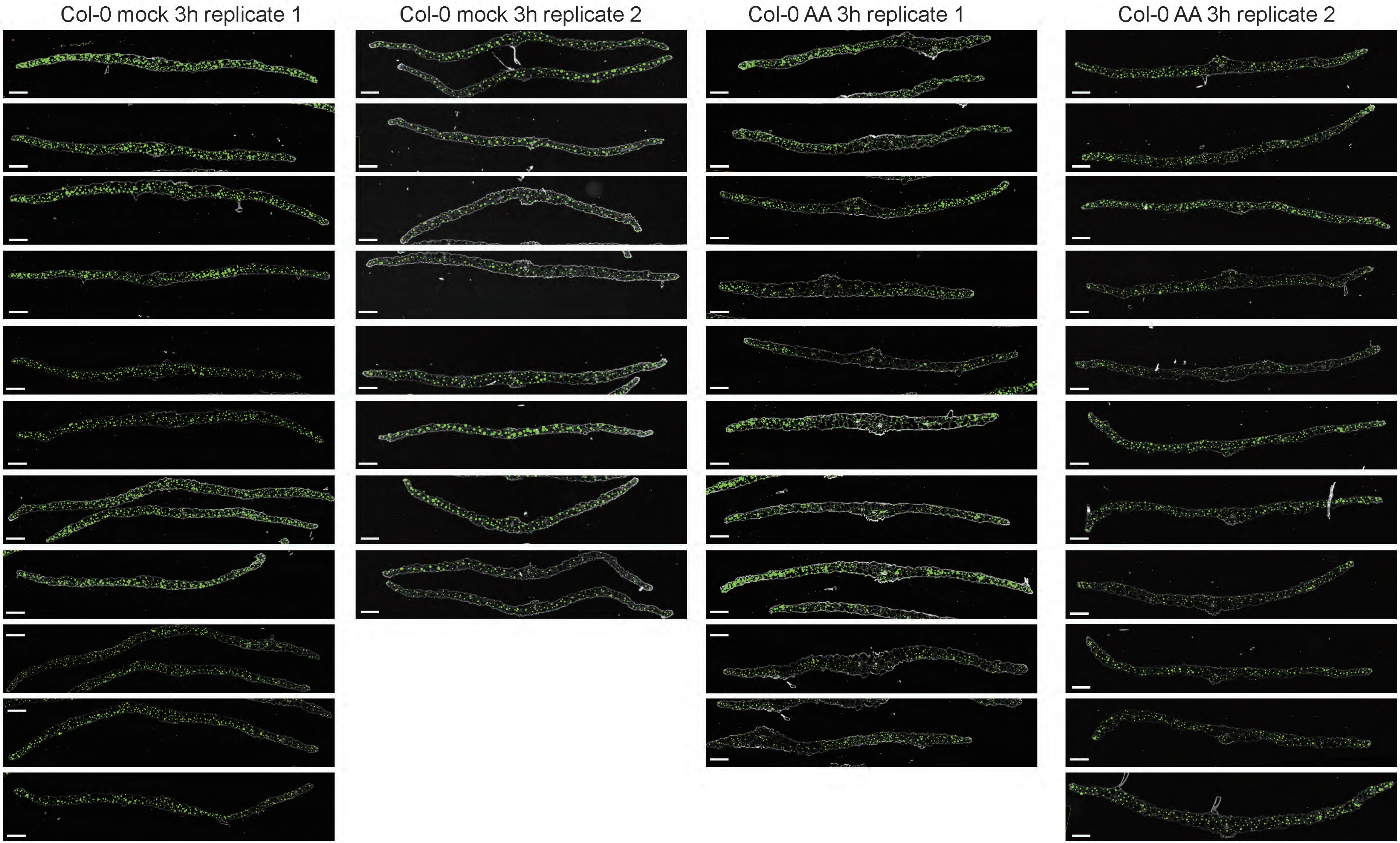

AT1G67870 (Predicted Companion cell marker, marker of cluster 18 in integrated UMAP)
Bar = 100 μm.

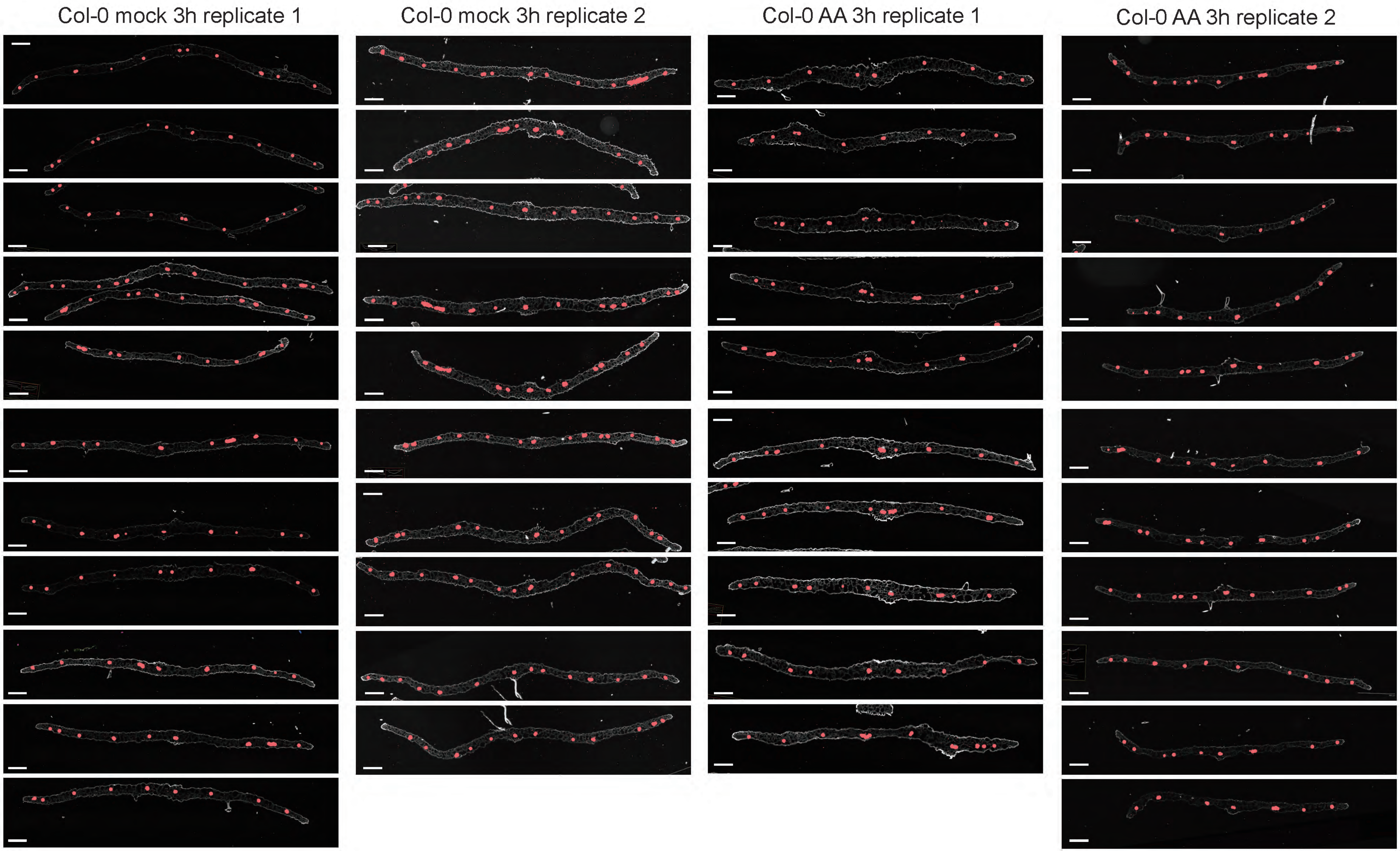

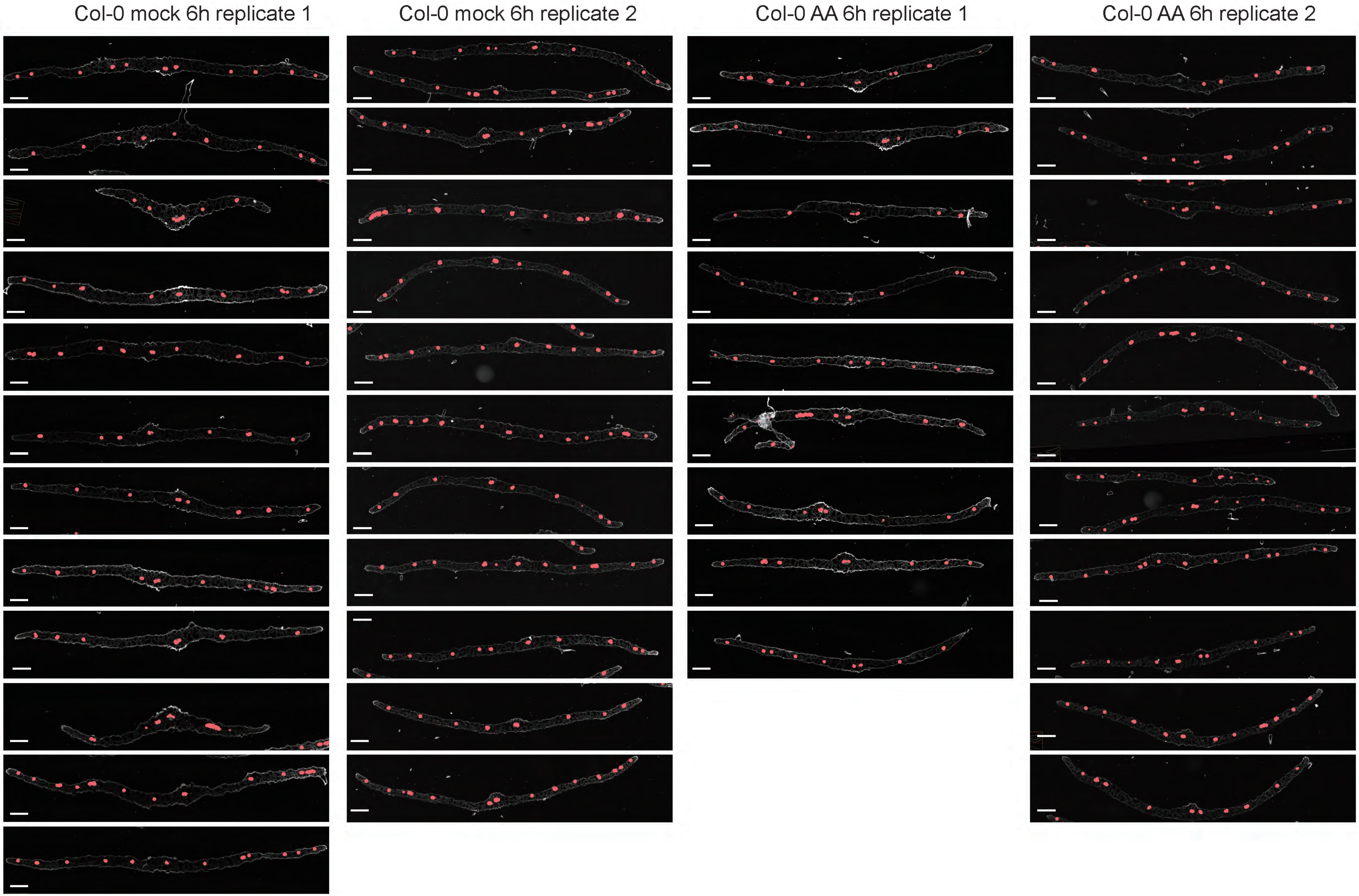
Col-0 mock 6h replicate 1
Col-0 mock 6h replicate 2
Col-0 AA 6h replicate 1
Col-0 AA 6h replicate 2

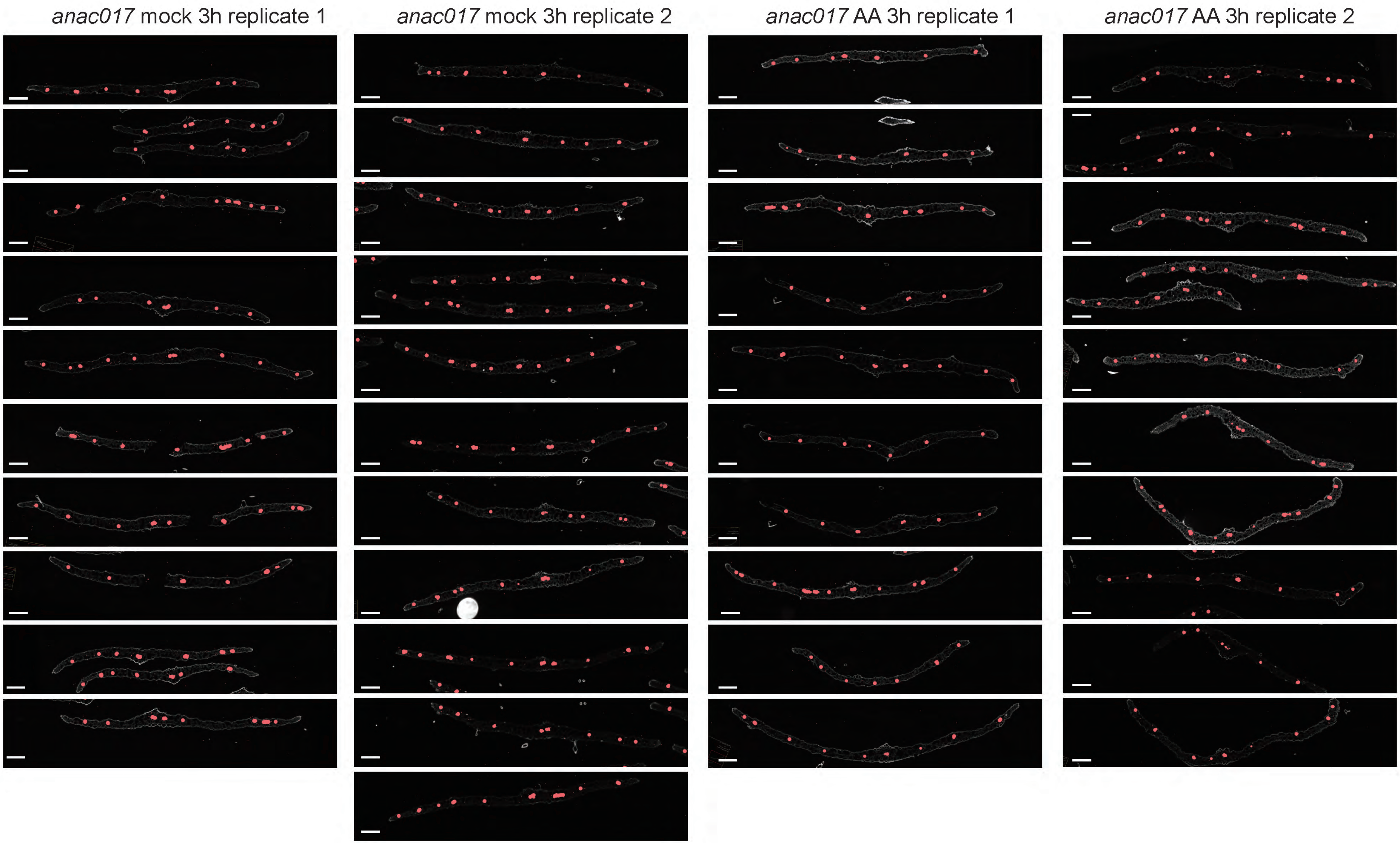
*anac017* mock 3h replicate 1
*anac017* mock 3h replicate 2
*anac017* AA 3h replicate 1
*anac017* AA 3h replicate 2

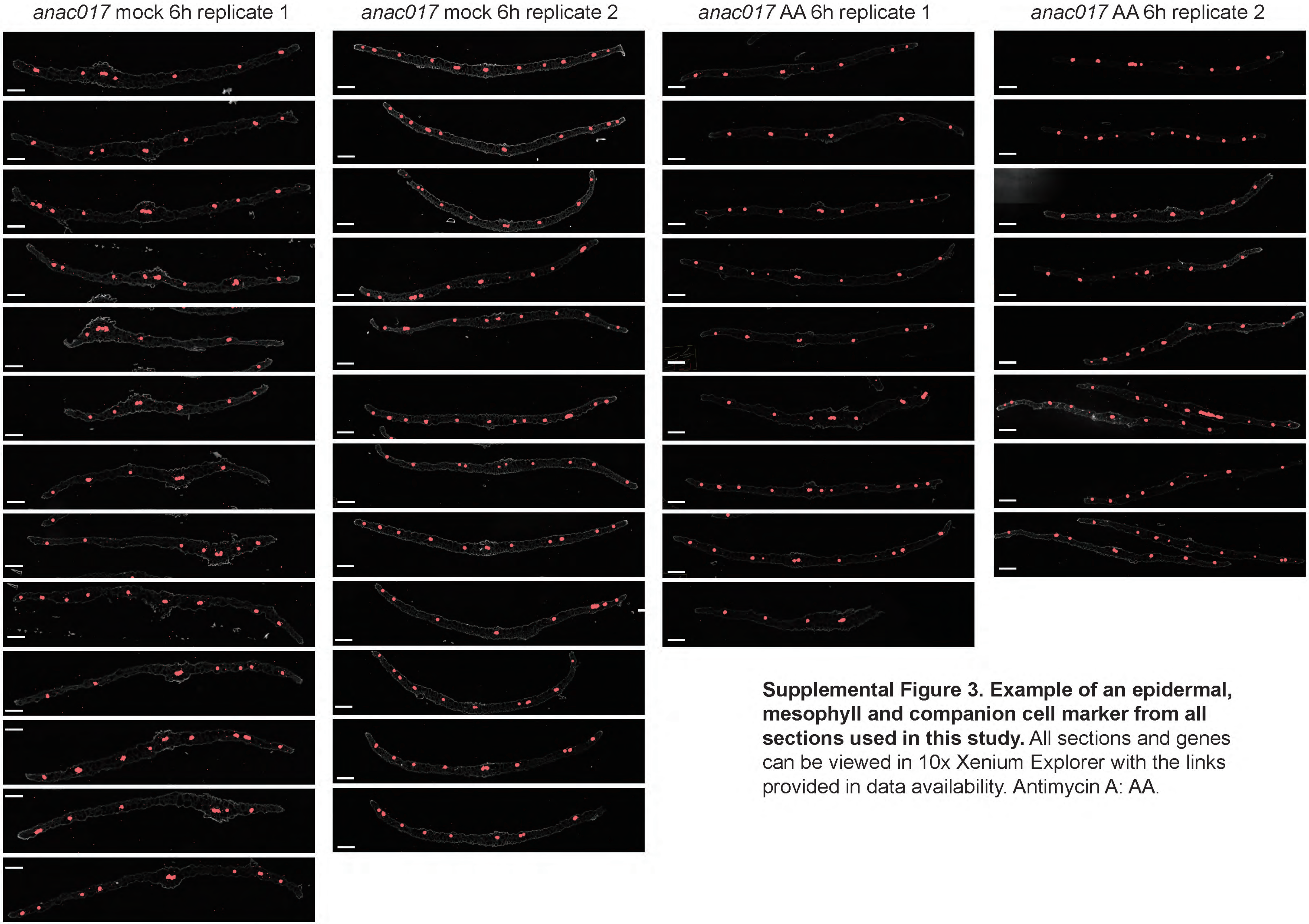


**Supplemental Figure 3. Example of an epidermal, mesophyll and companion cell marker from all sections used in this study.** All sections and genes can be viewed in 10x Xenium Explorer with the links provided in data availability. Antimycin A: AA.

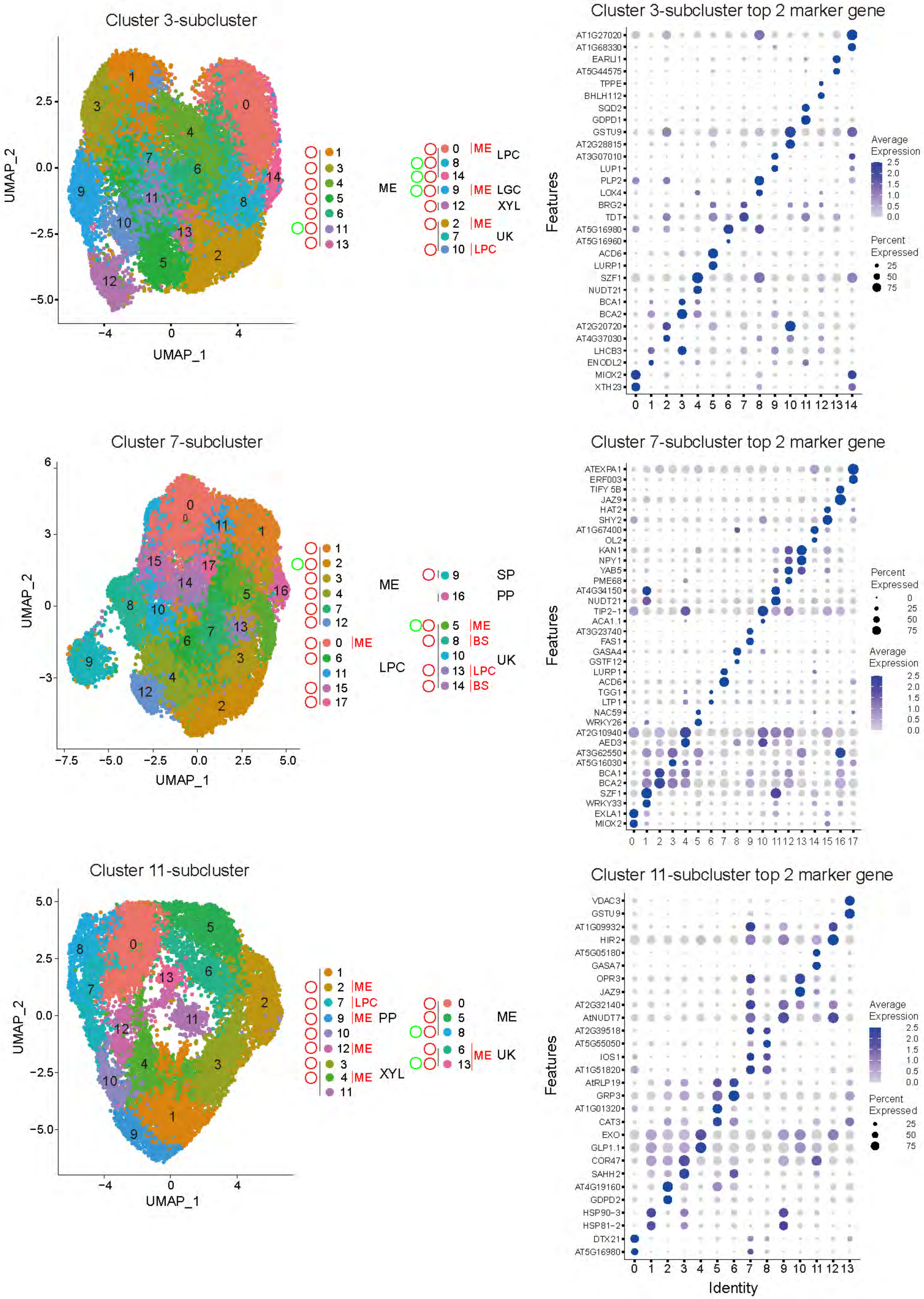

Cluster 3-subcluster
UMAP_1
UMAP_2
ME
LPC
LGC
XYL
UK
Cluster 3-subcluster top 2 marker gene
AT1G27020
AT1G68330
EARLI1
AT5G44575
TPPE
BHLH112
SQD2
GDPD1
GSTU9
AT2G28815
AT3G07010
LUP1
PLP2
LOX4
BRG2
TDT
AT5G16980
AT5G16960
ACD6
LURP1
SZF1
NUDT21
BCA1
BCA2
AT2G20720
AT4G37030
LHCB3
ENODL2
MIOX2
XTH23
Features
Average Expression
Percent Expressed
Cluster 7-subcluster
SP
PP
BS
Cluster 7-subcluster top 2 marker gene
ATEXPA1
ERF003
TIFY 5B
JAZ9
HAT2
SHY2
AT1G67400
OL2
KAN1
NPY1
YAB5
PME68
AT4G34150
NUDT21
TIP2-1
ACA1.1
AT3G23740
FAS1
GASA4
GSTF12
LURP1
ACD6
TGG1
LTP1
NAC59
WRKY26
AT2G10940
AED3
AT3G62550
AT5G16030
BCA1
BCA2
SZF1
WRKY33
EXLA1
MIOX2
Cluster 11-subcluster
Cluster 11-subcluster top 2 marker gene
VDAC3
GSTU9
AT1G09932
HIR2
AT5G05180
GASA7
OPR3
JAZ9
AT2G32140
AtNUDT7
AT2G39518
AT5G55050
IOS1
AT1G51820
AtRLP19
GRP3
AT1G01320
CAT3
EXO
GLP1.1
COR47
SAHH2
AT4G19160
GDPD2
HSP90-3
HSP81-2
DTX21
AT5G16980
Identity

**Supplemental Figure 4. Sub-clustering resolves heterogeneous cell states within three stress-responsive unknown clusters.**
Single-nucleus RNA sequencing data were re-analysed by extracting nuclei from cluster 3, cluster 7 and cluster 11 and performing independent unsupervised subclustering for each parent cluster. For each parent cluster, the left panel shows a UMAP embedding of nuclei coloured by subcluster identity (numbers indicate subclusters). The legend adjacent to each UMAP indicates the dominant cell-type assignment for each subcluster based on marker enrichment (ME, mesophyll; LPC, leaf pavement cell; EP, epidermis; LGC, leaf guard cell; XYL, xylem; PP, phloem parenchyma; SP, S phase; UK, unknown), highlighting that each parent cluster contains multiple transcriptionally distinct states.

For each parent cluster, the right panel shows a dot plot of the top two marker genes per subcluster, ranked by differential expression within that subclustering analysis. Dot colour intensity represents average scaled expression, and dot size represents the percentage of nuclei within each subcluster expressing the gene. The x-axis indicates subcluster identity, and the y-axis lists marker genes. As outlined in Figure 1 the predicted cell types were confirmed with using 10x Xenium Chip cellular identification (red dot), promoter GFP localization (green dot) or verified published markers (black dot). (see Supplemental Table 1 for cell annotation).

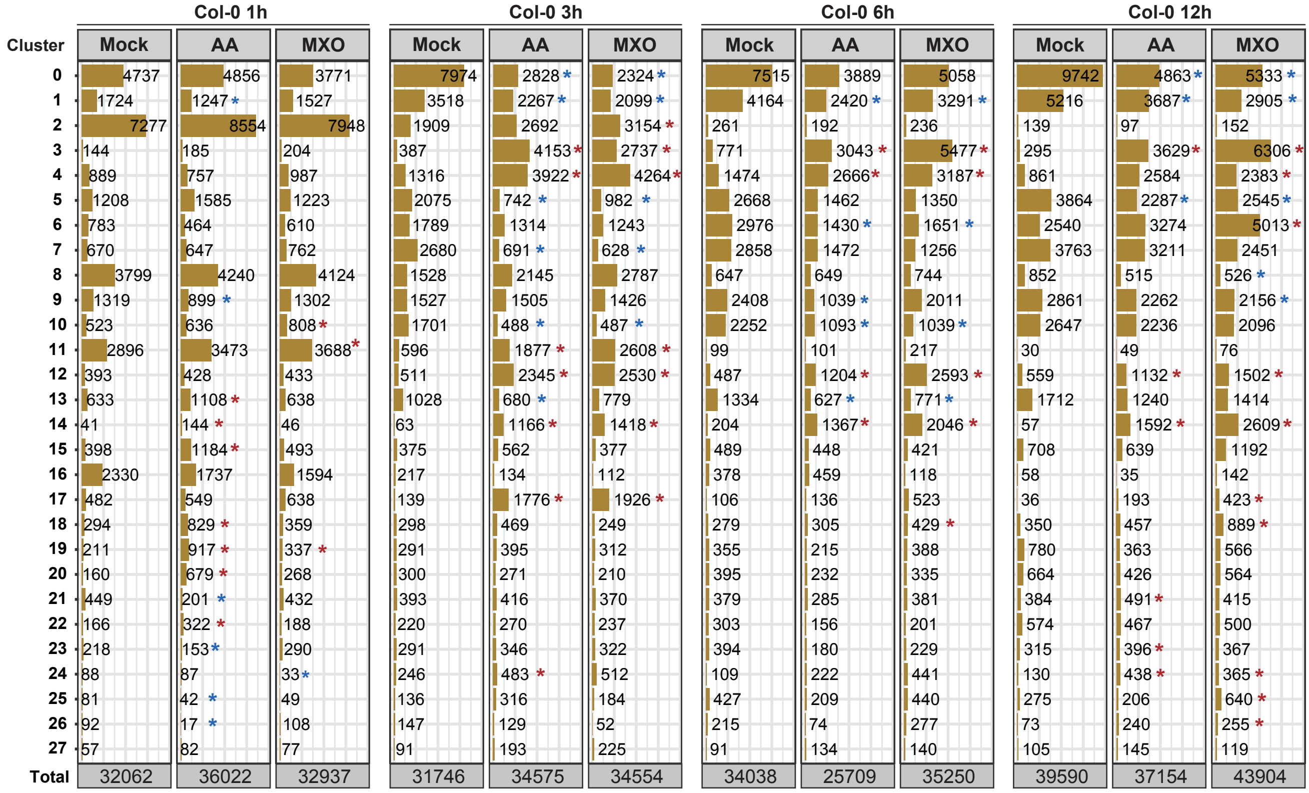


**Supplemental Figure 5. Time-resolved shifts in cluster abundance during mitochondrial inhibition in Col-0.**
Nuclei counts and relative abundance for each snRNA-seq cluster (rows, clusters 0–27) in Col-0 after 1 h, 3 h, 6 h, and 12 h under mock, antimycin A (AA), or myxothiazol (MXO) treatment (columns). For each time point, AA and MXO are compared to the time-matched mock control. Asterisks denote clusters with a significant change in abundance relative to the matched mock control, with red indicating an increase and blue indicating a decrease. Total nuclei analysed per condition are shown at the bottom of each column. Student's t-test with p  0.05, n = 3.

Figure 2.

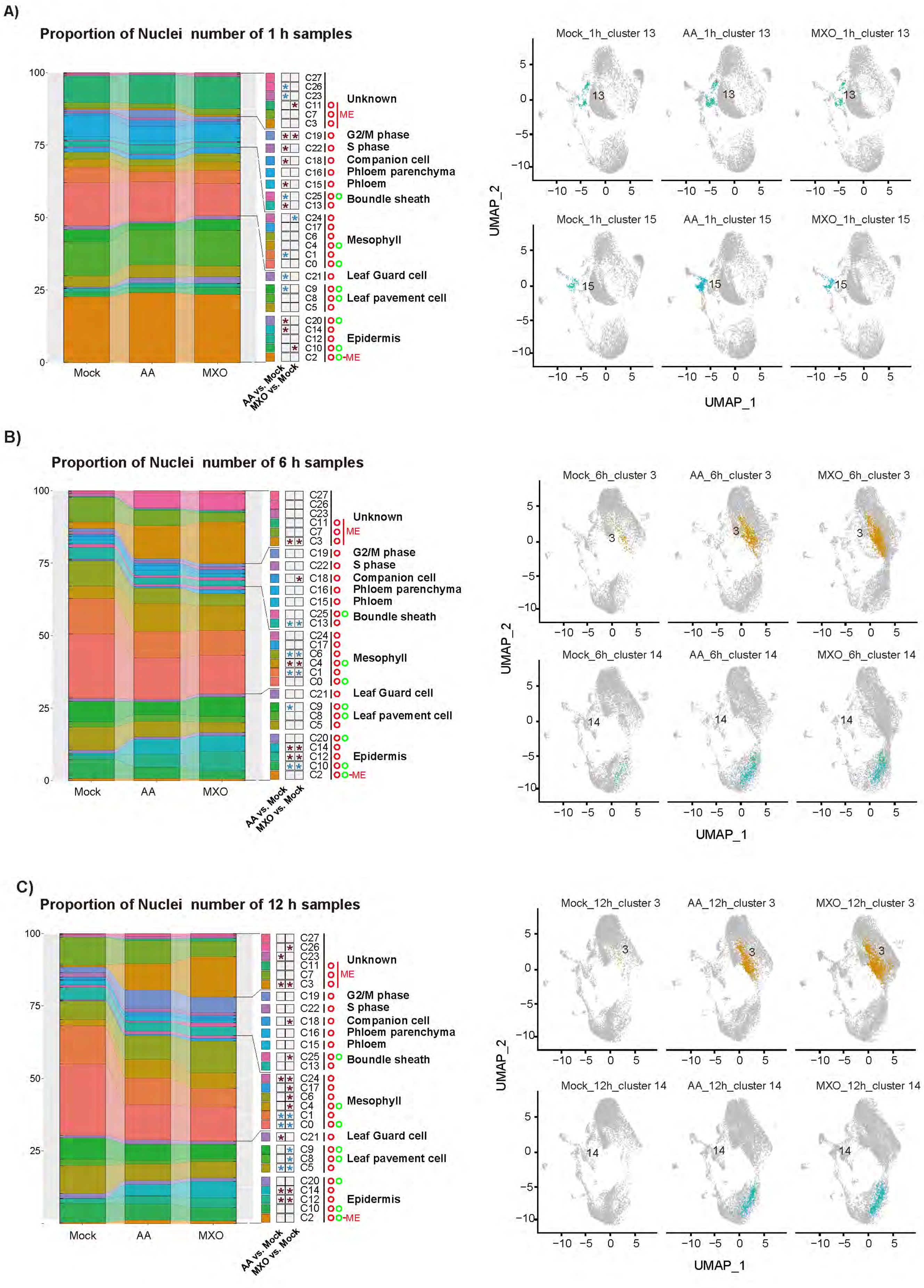


**Supplemental Figure 6. The transcriptional response to mitochondrial dysfunction after 1 (A), 6 (B) and 12 h (C) treatment with antimycin A or myxothiazol**. The left panel depicts the changes in the number of nuclei after 1, 6 and 12 h treatment. The clusters are sorted according to cell type. A red asterisk indicated a significant increase in number and a blue start indicates a significant decrease in number. The right panel depicts examples of clusters that are significantly increased in the treated compared to control samples. (see Supplemental Table 1 for cell annotation).

A)

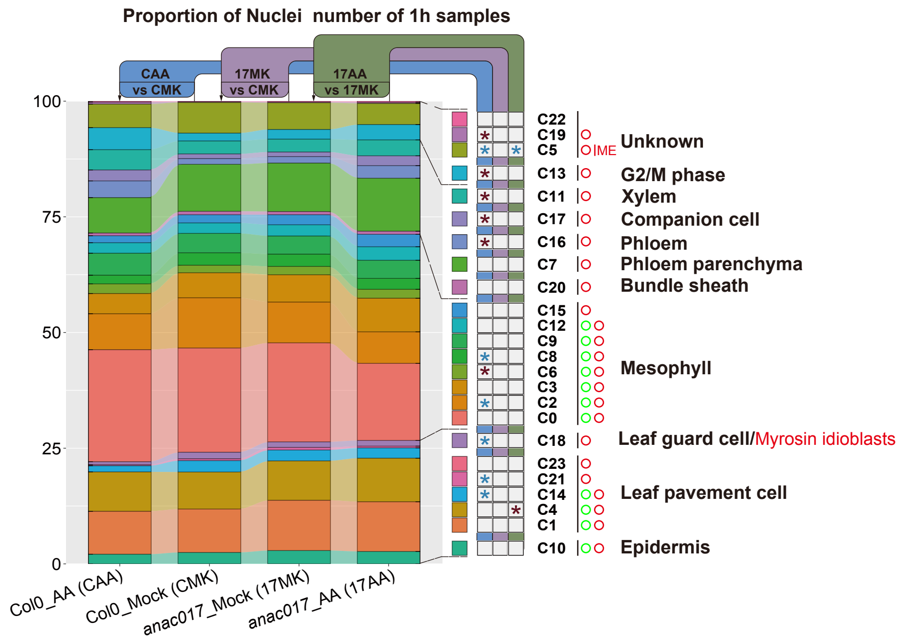


B)

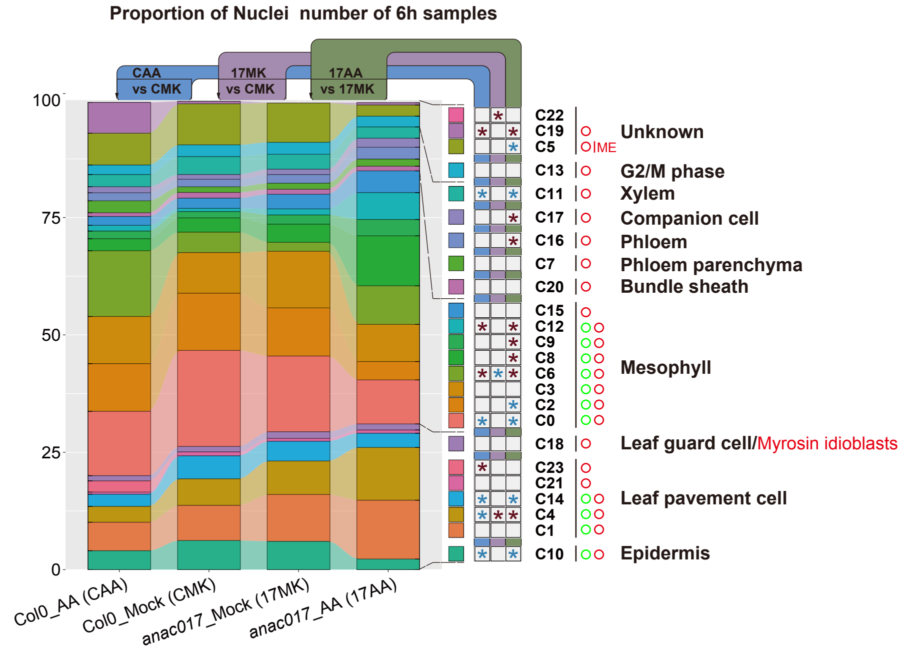


**Supplemental Figure 7. The response of Col-0 and anac017 to antimycin A (AA) treatment at A) 1 and B) 6 h.** Three different comparisons were carried out i) The response of Col-0 mock to AA, indicated with blue, ii) Col-0 mock treated compared to anac017 mock treated, indicated in purple, and iii) the response of anac017 to antimycin A treatment, indicated in green. A red asterisk indicated a significant increase in number and a blue asterisk indicates a significant decrease in numbers.

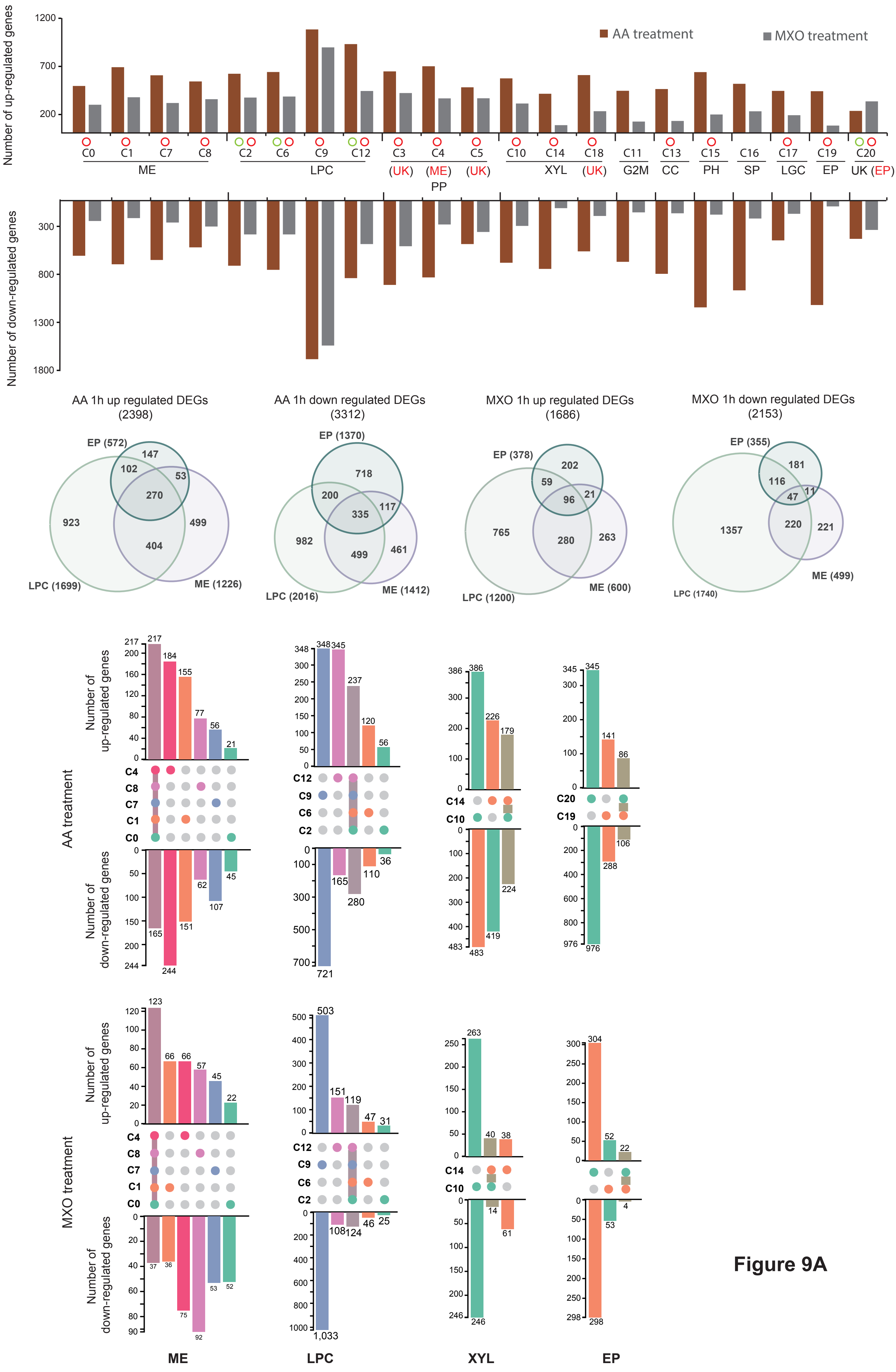
AA treatment
MXO treatment
Number of up-regulated genes
Number of down-regulated genes
C0 C1 C7 C8
ME
C2 C6 C9 C12
LPC
C3 C4 C5
(UK) (ME) (UK)
PP
C10 C14 C18
XYL
(UK)
C11
G2M
C13
CC
C15
PH
C16
SP
C17
LGC
C19
EP
C20
UK (EP)
AA 1h up regulated DEGs
(2398)
EP (572)
LPC (1699)
ME (1226)
AA 1h down regulated DEGs
(3312)
EP (1370)
LPC (2016)
ME (1412)
MXO 1h up regulated DEGs
(1686)
EP (378)
LPC (1200)
ME (600)
MXO 1h down regulated DEGs
(2153)
EP (355)
LPC (1740)
ME (499)
AA treatment
MXO treatment
Number of up-regulated genes
Number of down-regulated genes
C4
C8
C7
C1
C0
C12
C9
C6
C2
C14
C10
C20
C19
ME
LPC
XYL
EP

Figure 9A

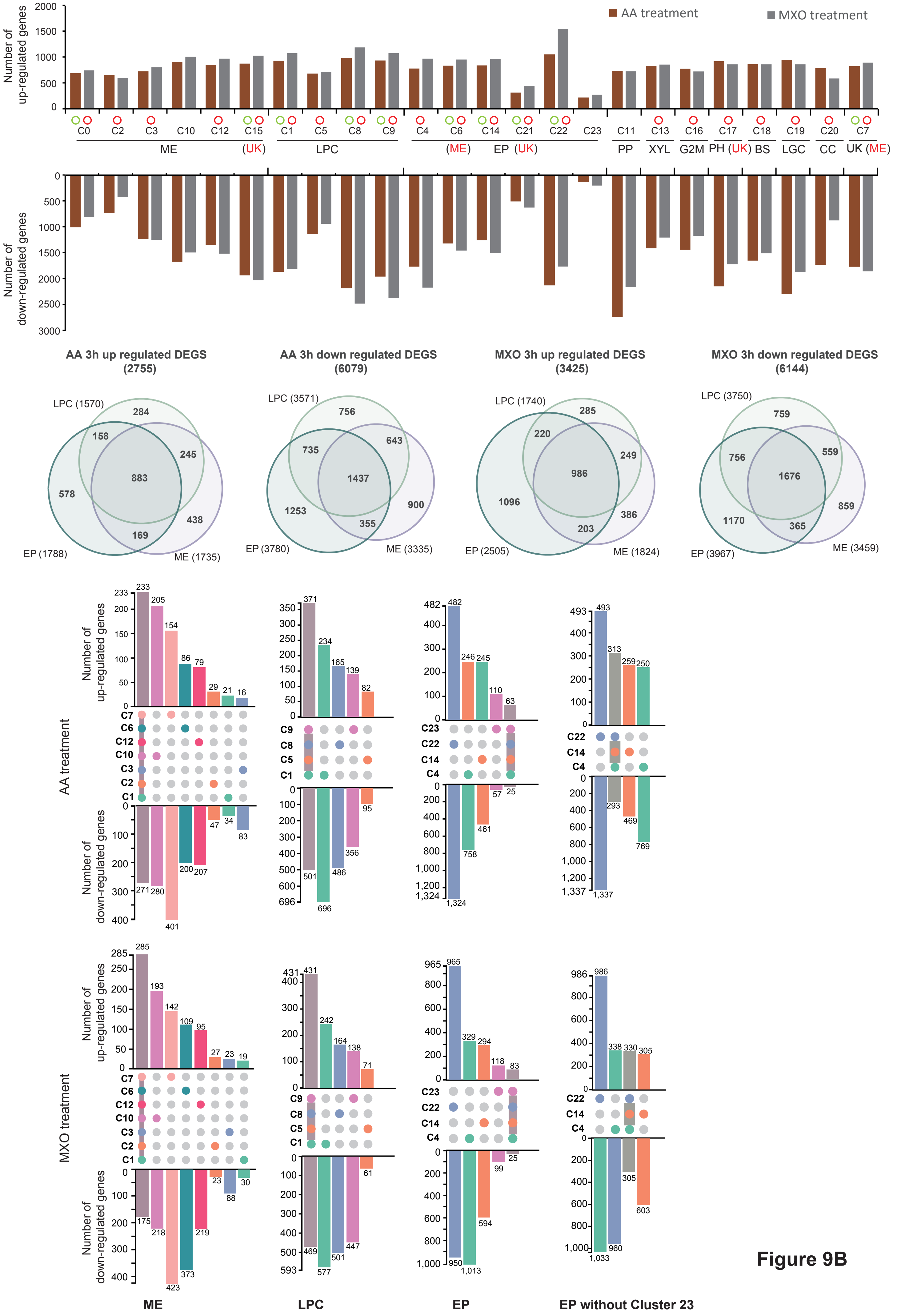

AA treatment
MXO treatment
Number of up-regulated genes
Number of down-regulated genes
C0 C2 C3 C10 C12 C15 C1 C5 C8 C9 C4 C6 C14 C21 C22 C23 C11 C13 C16 C17 C18 C19 C20 C7
ME (UK) LPC (ME) EP (UK) PP XYL G2M PH (UK) BS LGC CC UK (ME)
AA 3h up regulated DEGS (2755)
AA 3h down regulated DEGS (6079)
MXO 3h up regulated DEGS (3425)
MXO 3h down regulated DEGS (6144)
LPC (1570) EP (1788) ME (1735)
LPC (3571) EP (3780) ME (3335)
LPC (1740) EP (2505) ME (1824)
LPC (3750) EP (3967) ME (3459)
AA treatment
MXO treatment
ME
LPC
EP
EP without Cluster 23


**Figure 9B**

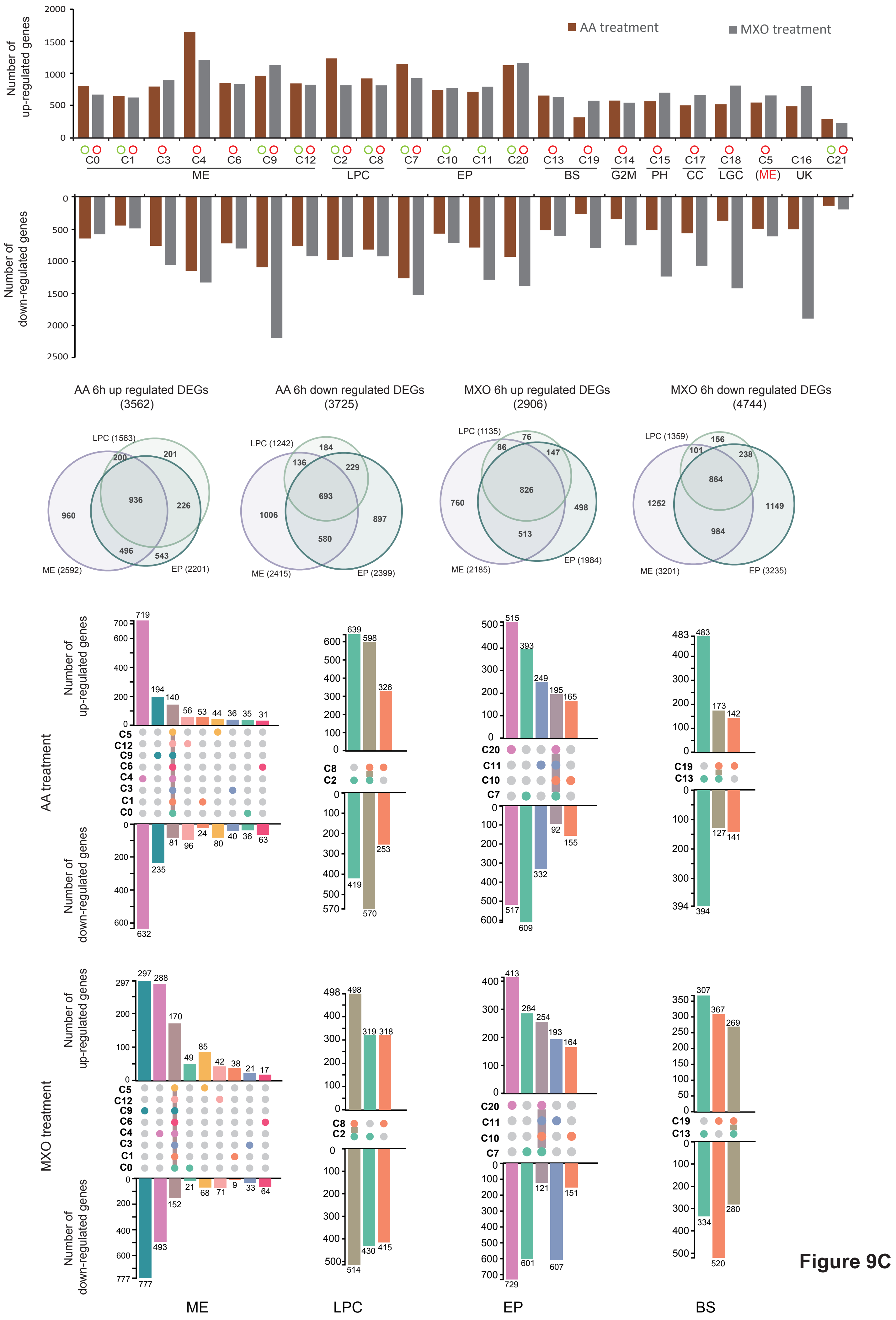
AA treatment
MXO treatment
Number of up-regulated genes
Number of down-regulated genes
C0 C1 C3 C4 C6 C9 C12 C2 C8 C7 C10 C11 C20 C13 C19 C14 C15 C17 C18 C5 C16 C21
ME
LPC
EP
BS
G2M
PH
CC
LGC
(ME)
UK
AA 6h up regulated DEGs (3562)
AA 6h down regulated DEGs (3725)
MXO 6h up regulated DEGs (2906)
MXO 6h down regulated DEGs (4744)
LPC (1563)
ME (2592)
EP (2201)
LPC (1242)
ME (2415)
EP (2399)
LPC (1135)
ME (2185)
EP (1984)
LPC (1359)
ME (3201)
EP (3235)
AA treatment
MXO treatment
ME
LPC
EP
BS

Figure 9C

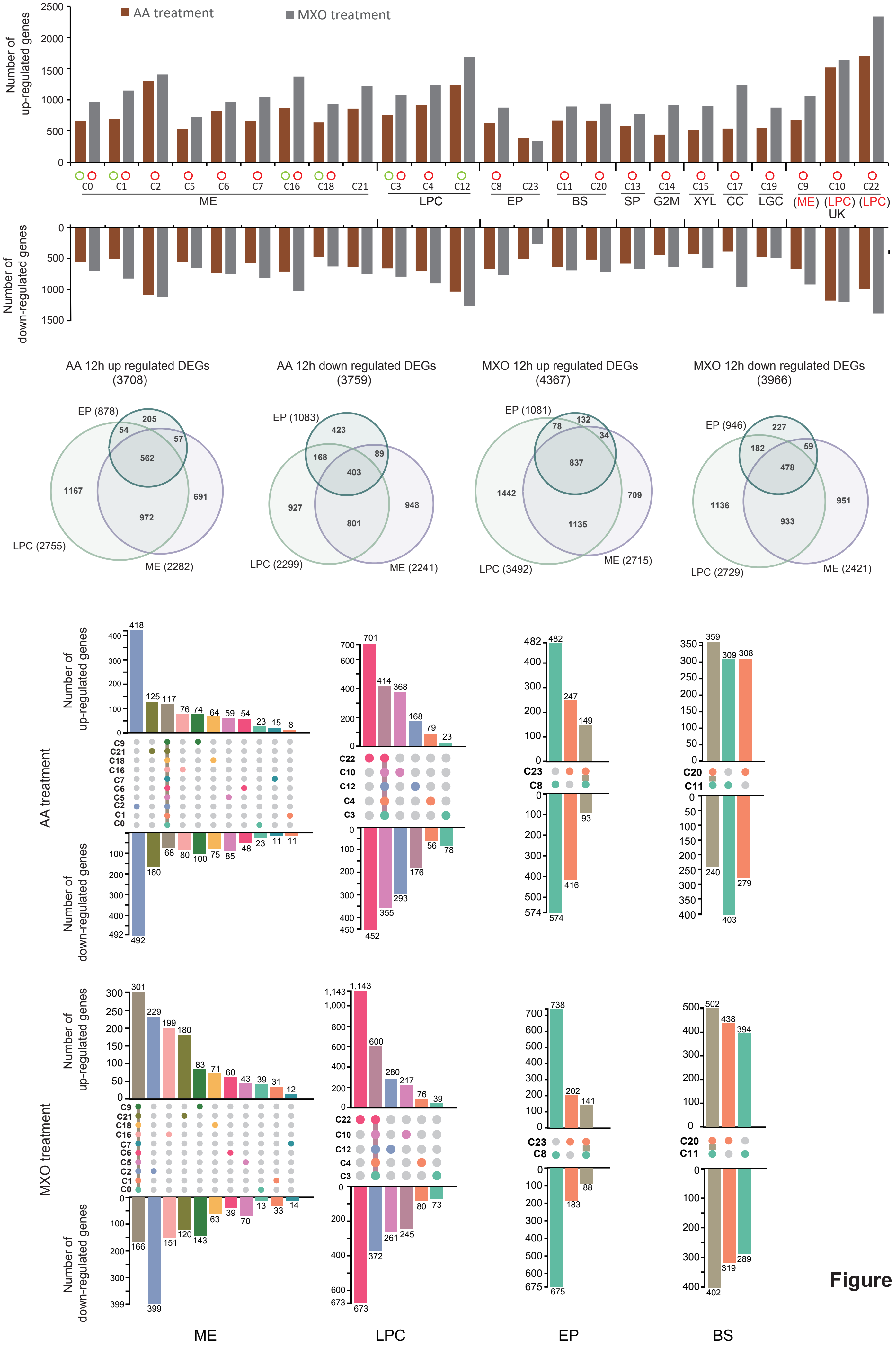
AA treatment
MXO treatment
Number of up-regulated genes
Number of down-regulated genes
C0 C1 C2 C5 C6 C7 C16 C18 C21
ME
C3 C4 C12
LPC
C8 C23
EP
C11 C20
BS
C13
SP
C14
G2M
C15
XYL
C17
CC
C19
LGC
C9 C10 C22
(ME) (LPC) (LPC)
UK
AA 12h up regulated DEGs (3708)
AA 12h down regulated DEGs (3759)
MXO 12h up regulated DEGs (4367)
MXO 12h down regulated DEGs (3966)
EP (878)
LPC (2755)
ME (2282)
EP (1083)
LPC (2299)
ME (2241)
EP (1081)
LPC (3492)
ME (2715)
EP (946)
LPC (2729)
ME (2421)
AA treatment
MXO treatment
ME
LPC
EP
BS

Figure 9D

**Supplemental Figure 9. A gene level view of the mitochondrial stress response at a cellular level at A) 1, B) 3, C) 6 and D 12 h. Top Panel -** The number of genes that changed in transcript abundance after 1, 3, 6 and 12 hours of antimycin A (AA) and myxothiazol (MXO) treatment (see Supplemental Table 6 for full details). mesophyll (ME), epidermis (EP), leaf pavement cells (LPC), phloem parenchyma (PP), xylem (XYL), bundle sheath (BS), phloem (PH), companion cells (CC), leaf guard cells (LGC), G2/M phase (G2M), S phase (SP) and unknown (UK). **Venn Diagrams** - The number of genes that were increased or decreased in transcript abundance after 1-, 3-, 6- and 12-hours treatment with antimycin A or myxothiazol and how they overlap in epidermal (EP), leaf pavement cells (LPC) and mesophyll cells (ME). **Bottom Panel -** The common and unique genes that increased or decreased in the epidermal, leaf pavement cell, mesophyll and bundle sheath clusters after 1-, 3-, 6- and 12-hours treatment with antimycin A or myxothiazol if there are two or more clusters in the same cell type.

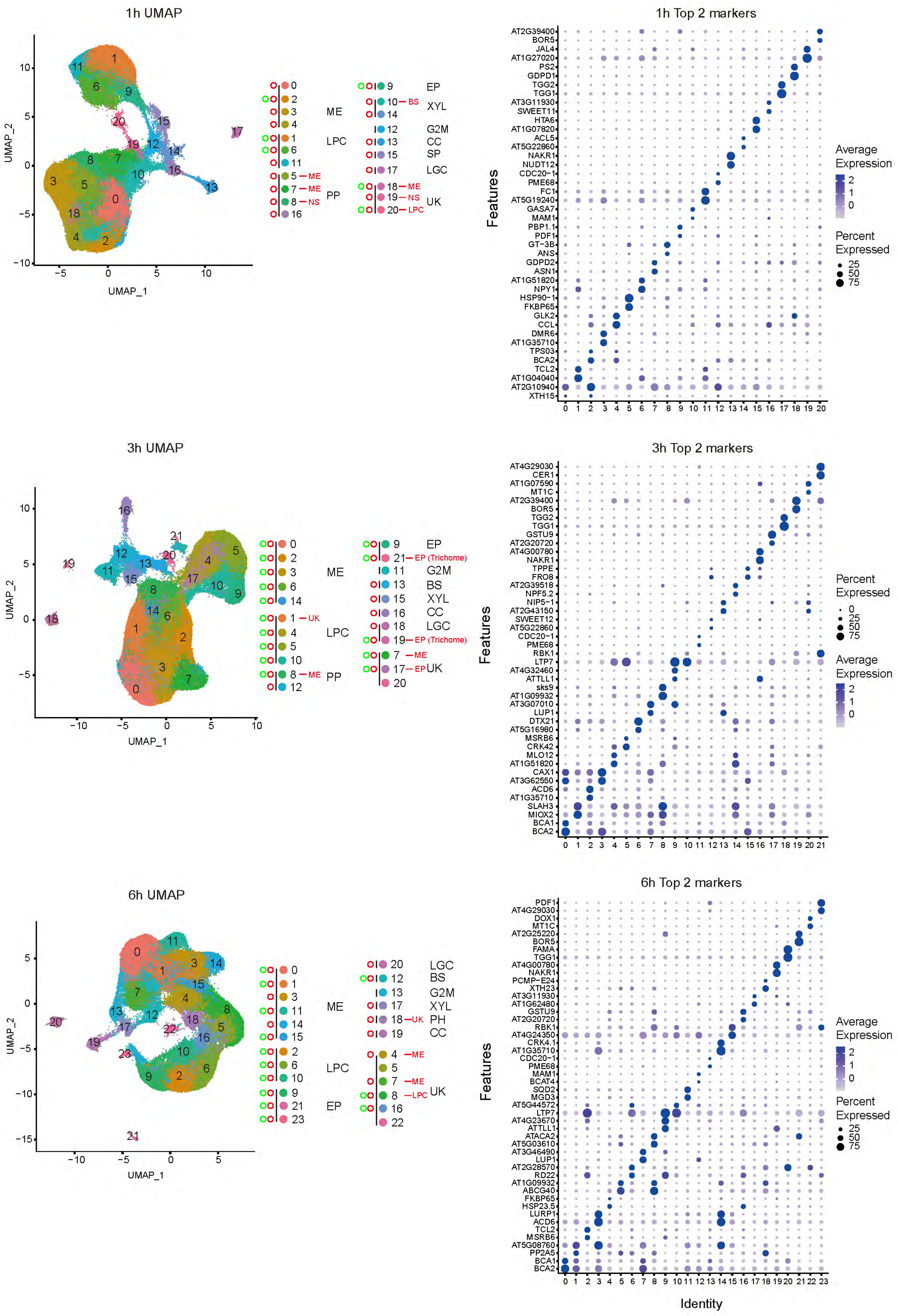
1h UMAP
UMAP_1
UMAP_2
ME
LPC
PP
EP
XYL
G2M
CC
SP
LGC
UK
10 — BS
5 — ME
7 — ME
8 — NS
18 — ME
19 — NS
20 — LPC
1h Top 2 markers
Features
AT2G39400
BOR5
JAL4
AT1G27020
PS2
GDPD1
TGG2
TGG1
AT3G11930
SWEET11
HTA6
AT1G07820
ACL5
AT5G22860
NAKR1
NUDT12
CDC20−1
PME68
FC1
AT5G19240
GASA7
MAM1
PBP1.1
PDF1
GT−3B
ANS
GDPD2
ASN1
AT1G51820
NPY1
HSP90−1
FKBP65
GLK2
CCL
DMR6
AT1G35710
TPS03
BCA2
TCL2
AT1G04040
AT2G10940
XTH15
Average Expression
Percent Expressed
3h UMAP
UMAP_1
UMAP_2
ME
LPC
PP
EP
G2M
BS
XYL
CC
LGC
UK
21 — EP (Trichome)
1 — UK
8 — ME
19 — EP (Trichome)
7 — ME
17 — EP
3h Top 2 markers
Features
AT4G29030
CER1
AT1G07590
MT1C
AT2G39400
BOR5
TGG2
TGG1
GSTU9
AT2G20720
AT4G00780
NAKR1
TPPE
FRO8
AT2G39518
NPF5.2
NIP5−1
AT2G43150
SWEET12
AT5G22860
CDC20−1
PME68
RBK1
LTP7
AT4G32460
ATTLL1
sks9
AT1G09932
AT3G07010
LUP1
DTX21
AT5G16980
MSRB6
CRK42
MLO12
AT1G51820
CAX1
AT3G62550
ACD6
AT1G35710
SLAH3
MIOX2
BCA1
BCA2
Percent Expressed
Average Expression
6h UMAP
UMAP_1
UMAP_2
ME
LPC
EP
LGC
BS
G2M
XYL
PH
CC
UK
18 — UK
4 — ME
7 — ME
8 — LPC
6h Top 2 markers
Features
PDF1
AT4G29030
DOX1
MT1C
AT2G25220
BOR5
FAMA
TGG1
AT4G00780
NAKR1
PCMP−E24
XTH23
AT3G11930
AT1G62480
GSTU9
AT2G20720
RBK1
AT4G24350
CRK4.1
AT1G35710
CDC20−1
PME68
MAM1
BCAT4
SQD2
MGD3
AT5G44572
LTP7
AT4G23670
ATTLL1
ATACA2
AT5G03610
AT3G46490
LUP1
AT2G28570
RD22
AT1G09932
ABCG40
FKBP65
HSP23.5
LURP1
ACD6
TCL2
MSRB6
AT5G08760
PP2A5
BCA1
BCA2
Average Expression
Percent Expressed
Identity

**Supplemental Figure 8. Unsupervised cell cluster of Col-0 and *anac017* at A) 1, B) 3, and C) 6 h treatment with antimycin A.** The samples were clustered using Sauret. The cell clusters were initially annotated using Single Cell Plant DataBase (scPlantDB) (Ryu *et al.*, 2021) and subsequently the cell types were confirmed with using 10x Xenium Chip cellular identification (red dot) and promoter GFP localization (green dot). For the purposes of reproducibility the clusters initially annotated as unknown are left annotated as unknown in this diagram in black text, with subsequent identification by 10x Xenium Chip and promoter GFP tagging identified the cell type, which is indicated in red (confirmed using 10x Xenium Spatial Transcriptome) or green (using promoter GFP-tagging) (see Supplemental Table 4 for cell annotation). mesophyll (ME), epidermis (EP), leaf pavement cells (LPC), bundle sheath (BS), phloem (PH), phloem parenchyma (PP), companion cells (CC), leaf guard cells (LGC), xylem (XYL), G2/M phase (G2M), S phase (SP) and unknown (UK).

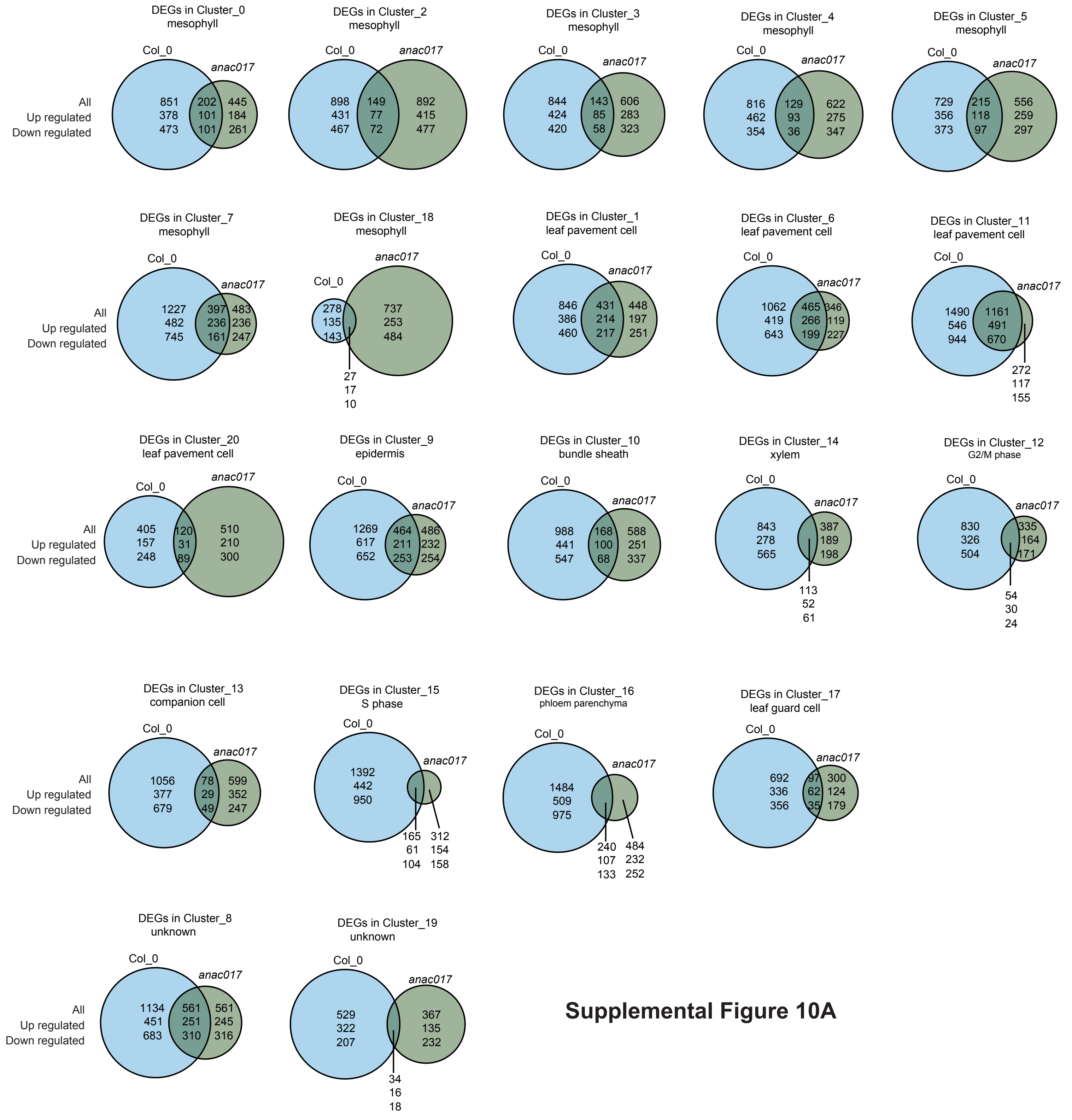


**Supplemental Figure 10A**

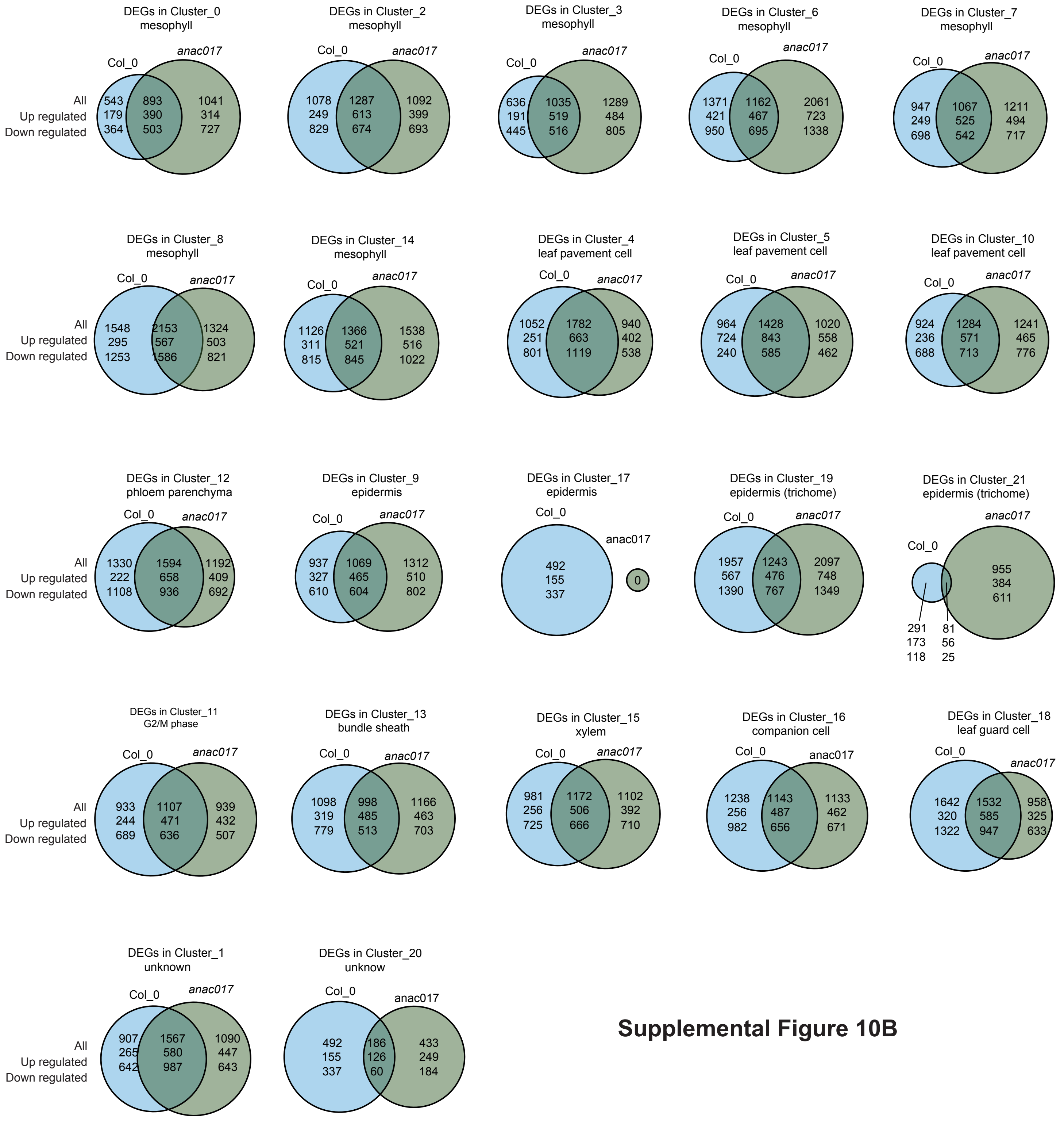


**Supplemental Figure 10B**

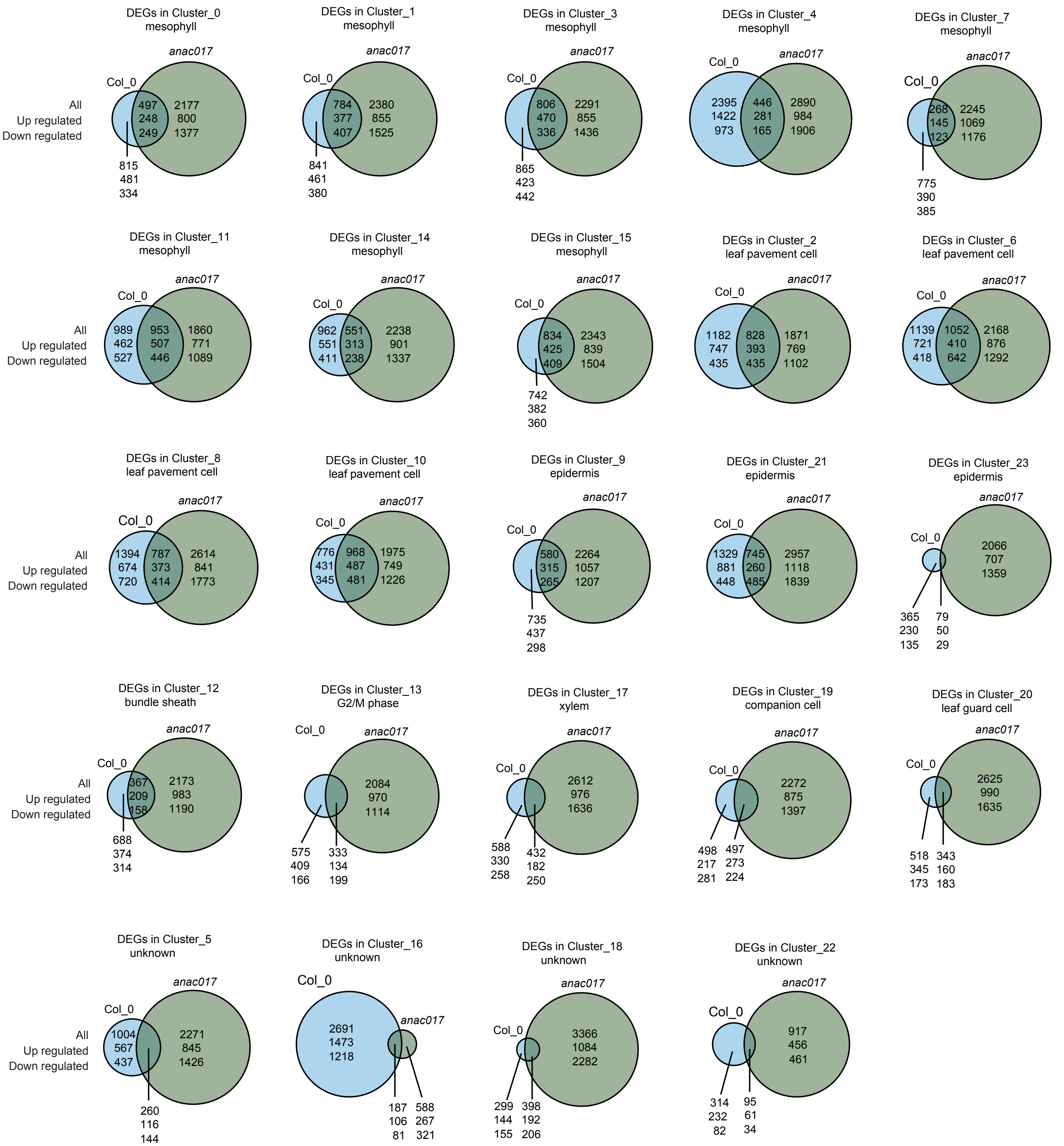


**Supplemental Figure 10C**

**Supplemental Figure 10. The number and overlap in differentially expressed genes (DEGs) in Col-0 and anac017 after A) 1, B) 3, and C) 6 hours of antimycin A treatment.** Venn diagrams show differentially expressed genes (DEGs) in Col-0 (blue) and anac017 (green) after antimycin A treatment for (A) 1 h, (B) 3 h, and (C) 6 h, resolved by snRNA-seq cluster (cluster ID and cell type indicated). Numbers denote total and shared DEGs, with up- and downregulated genes indicated where shown.

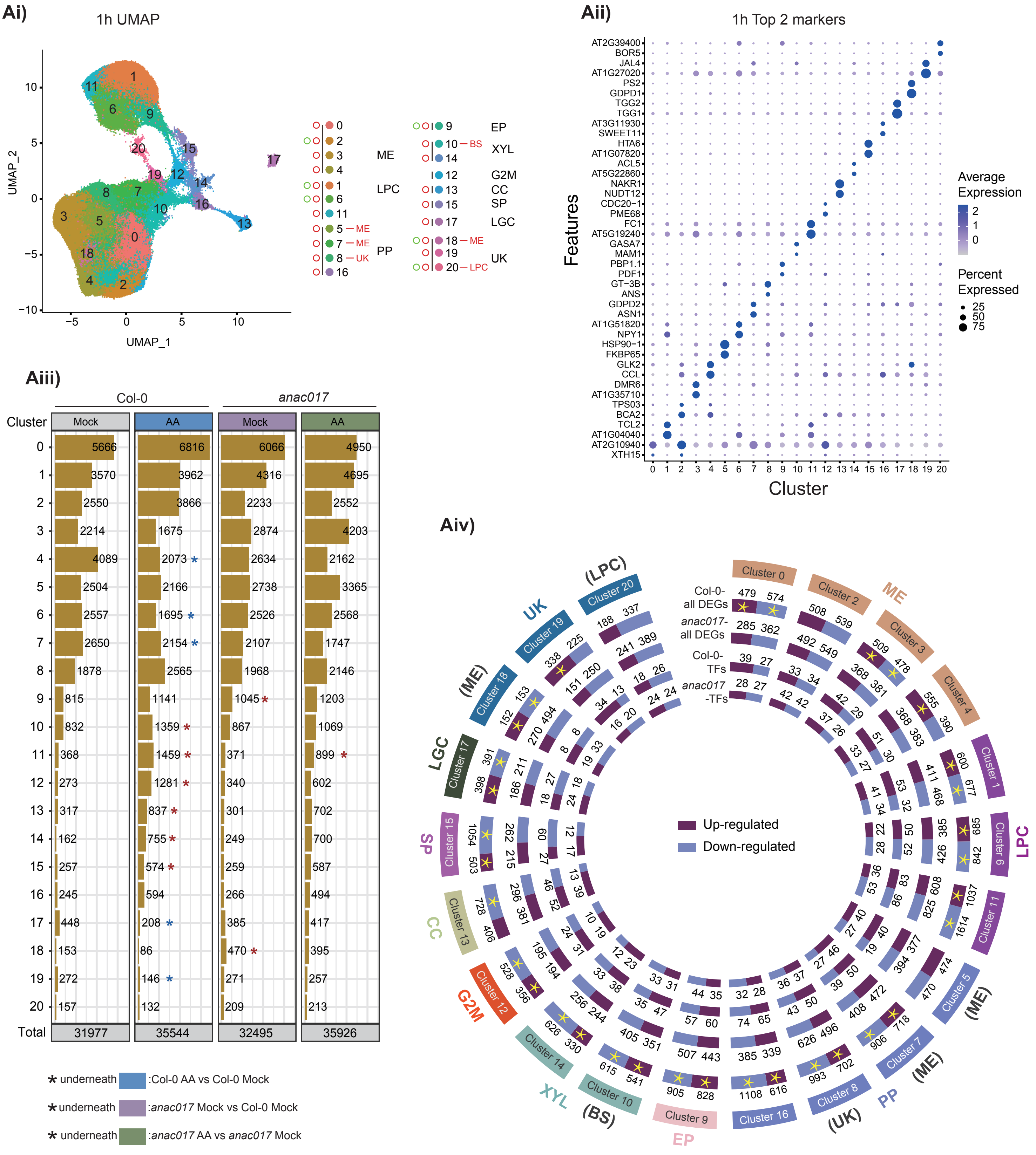


### Supplemental Figure 11A

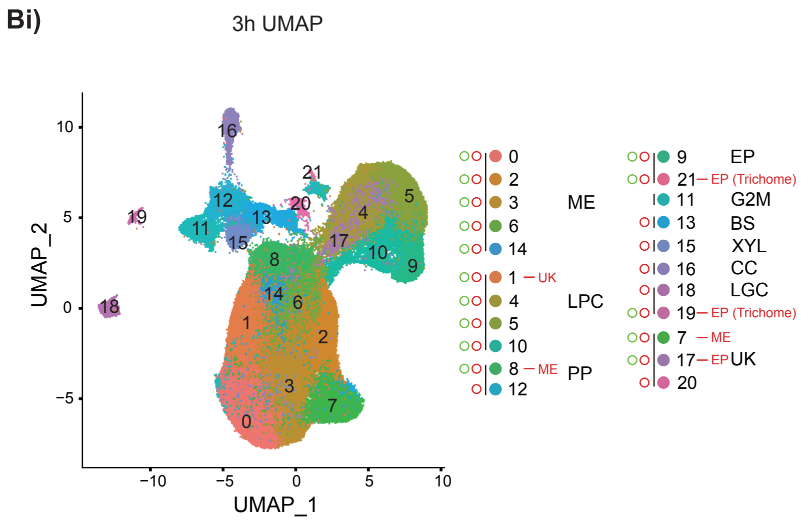


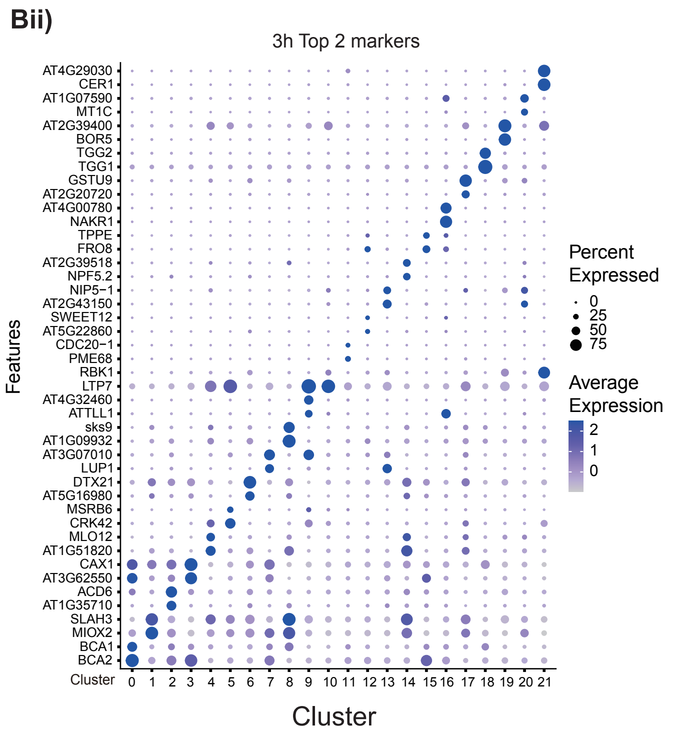


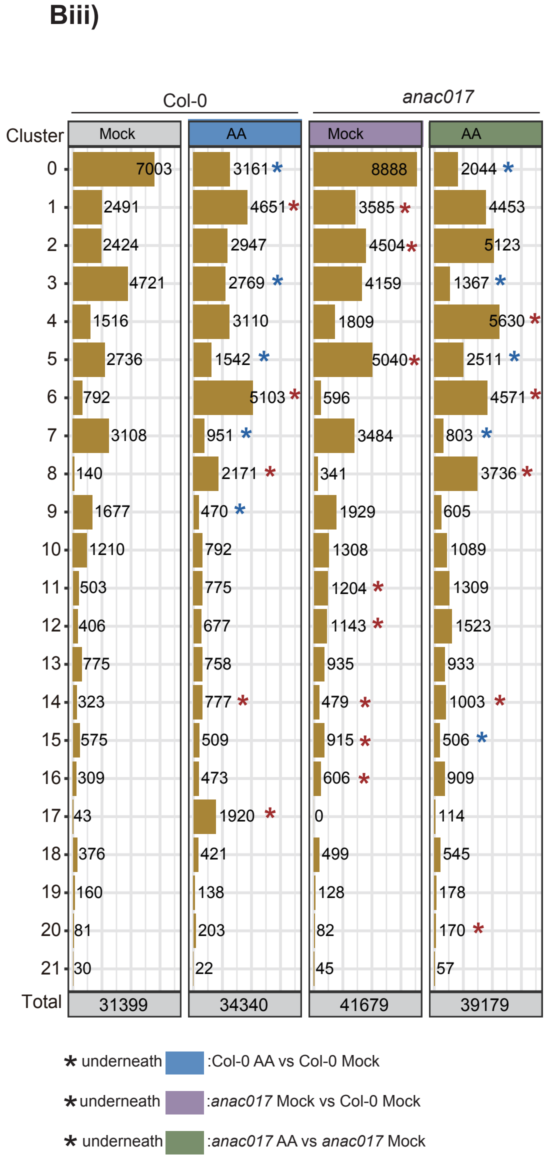


### Supplemental Figure 11B

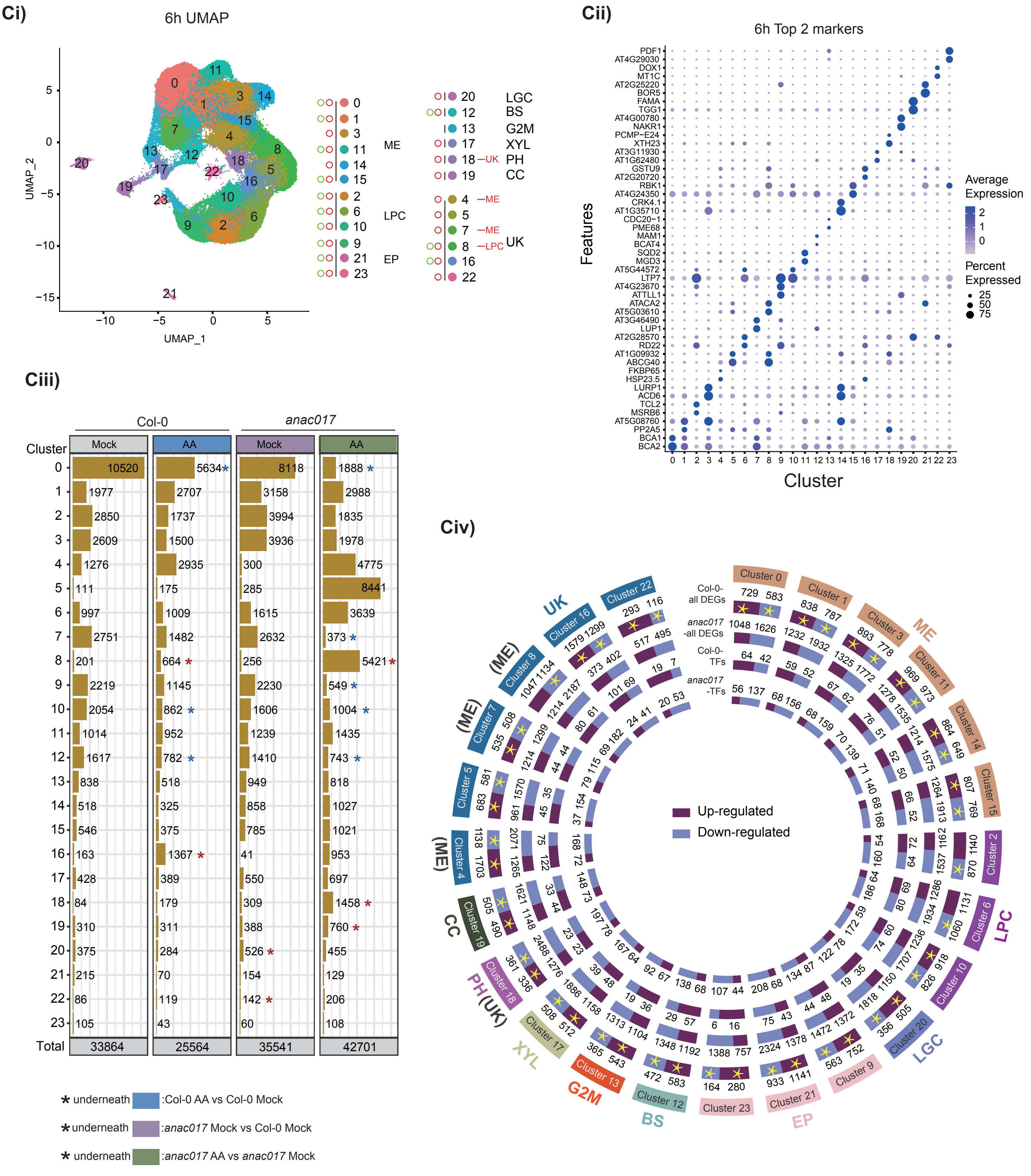


**Supplemental Figure 11C**

**Supplemental Figure11. Analysis of differentially expressed genes in Col-0 and *anac017* after A) 1, B) 3, and C) 6 hours of treatment with antimycin A. i)** UMAP cluster visualization of Col-0 and *anac017* after 1, 3 and 6 h treatment showing cell type that were predicted and re-annotated with experimental verification as outlined in Figure 2**. ii)** The top two marker genes in each cluster (see Supplemental Table 4). **iii)** The number of nuclei in each cluster in control (mock) and antimycin A (AA) treated samples after different times of treatment. Changes in the number of nuclei compared to control are indicated. Student's t-test with $p \leq 0.05$, n = 3. Three different comparison were carried out i) The response of Col-0 to AA, indicated with blue, ii) Col-0 mock treated compared to *anac017* mock treated, indicated in purple, and iii) the response *of anac017* to antimycin A treatment, indicated in green. A red asterisk indicated a significant increase in number and a blue asterisk indicates a significant decrease in numbers. **iv)** The total number of differentially expressed genes (DEGs) and genes encoding transcription factors in each cluster in Col-0 and *anac017* after **A)** 1, **B)** 3, and **C)** 6 hours of treatment with AA. The cluster number and cell type are indicated in the outer circle with clusters experimentally confirmed as in Figure 3. The yellow asterisks indicate a significance ($p < 0.001$) in the proportion of genes that changed in transcript abundance in Col-0 compared to *anac017* for each cluster based on all the total number of genes detected. The number of transcription factors is too small to carry out a proportionality test. mesophyll (ME), epidermis (EP), leaf pavement cells (LPC), phloem parenchyma (PP), xylem (XYL), bundle sheath (BS), phloem (PH), companion cells (CC), leaf guard cells (LGC), G2/M phase (G2M), S phase (SP) and unknown (UK). Clusters re-designated using experimentally based markers are indicated in brackets.

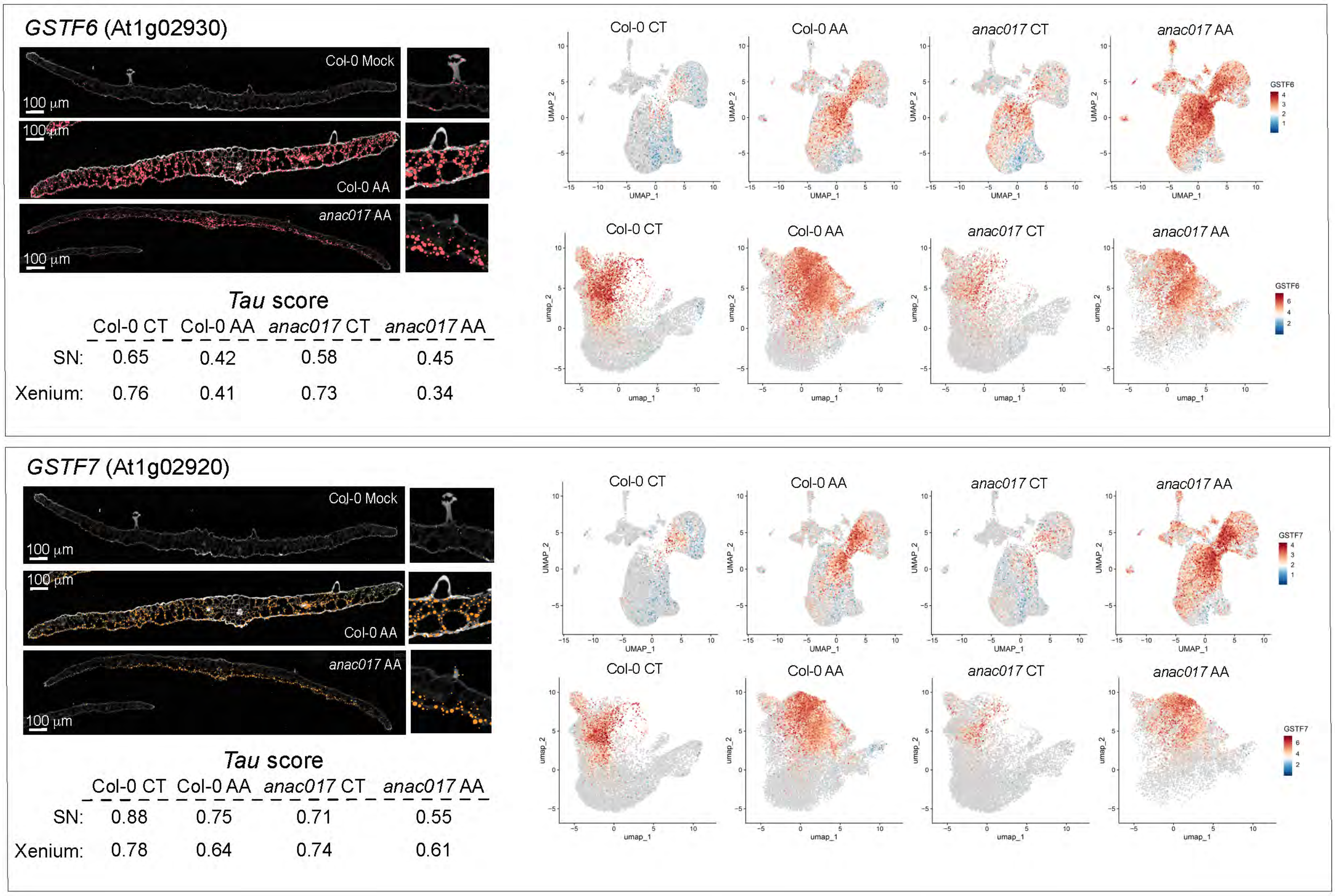


**Supplemental Figure 12. Single nuclei and spatial transcriptome generated UMAP projections of cell clusters after treatment with antimycin A for 3 h in Col-0 and *anac017*.** Each panel shows an individual gene with abbreviation and gene identity. The section for each treatment represents a single section of a total of 20-24 section for each treatment. The feature plots depict the UMAP projection for the snRNA-seq (top row in each panel) and spRNA-seq (bottom row in each panel). The calculated tau value, as a measure of expression specificity, where 1 is most specific and 0 is non-specific is shown. The feature plot clusters and cell types are designated in Figure 4. GSTF6 = Glutathione s-transferase 6, GSTF7 = Glutathione S-transferase 7, mock = control, AA = Antimycin A, SN = single nuclei.

*FAMA* (At3g24140)

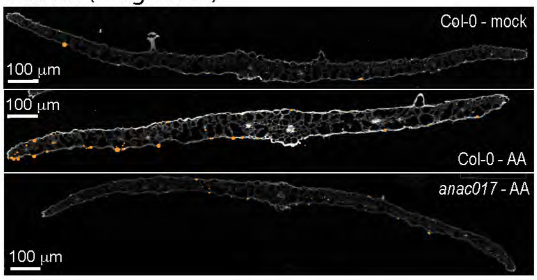


### *Tau* score

| | Col-0 CT | Col-0 AA | *anac017* CT | *anac017* AA |
|---|---|---|---|---|
| SN: | 0.99 | 0.99 | 0.99 | 0.99 |
| Xenium: | 0.99 | 0.99 | 0.99 | 0.99 |

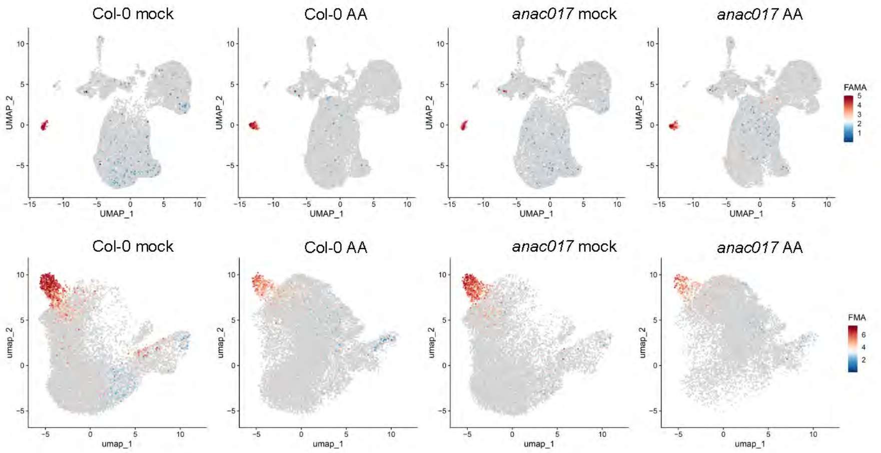


*HB-8* (At4g32880)

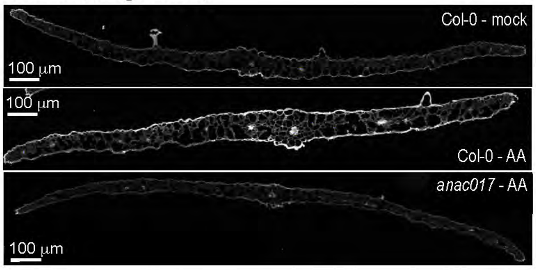


### *Tau* score

| | Col-0 CT | Col-0 AA | *anac017* CT | *anac017* AA |
|---|---|---|---|---|
| SN: | 0.90 | 0.92 | 0.92 | 0.94 |
| Xenium: | 0.88 | 0.89 | 0.94 | 0.87 |

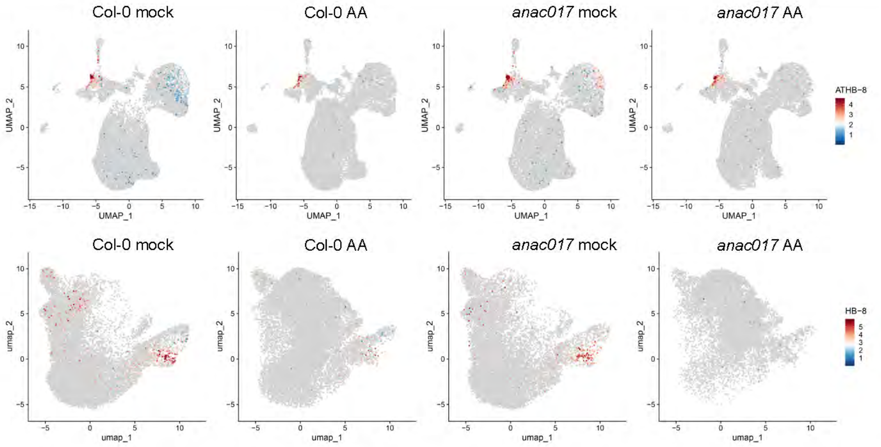


*MYB29* (At5g07690)

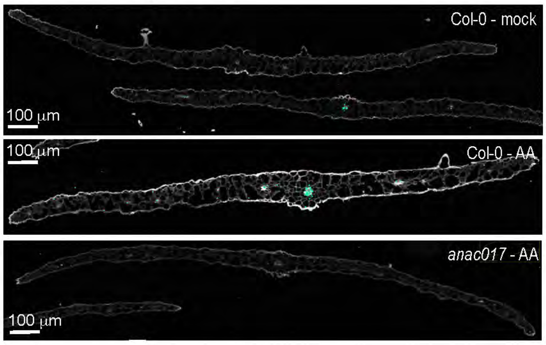


### *Tau* score

| | Col-0 CT | Col-0 AA | *anac017* CT | *anac017* AA |
|---|---|---|---|---|
| SN: | 0.86 | 0.84 | 0.82 | 0.86 |
| Xenium: | 0.96 | 0.68 | 0.92 | 0.78 |

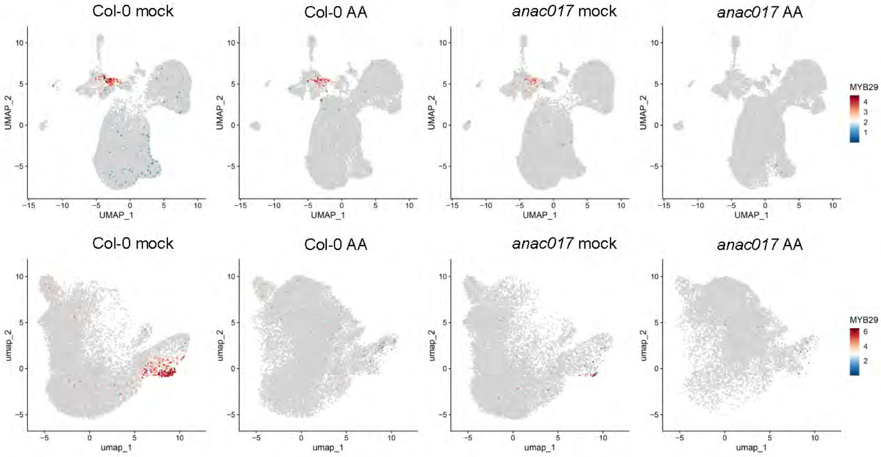


*MYB28* (At5g61420)

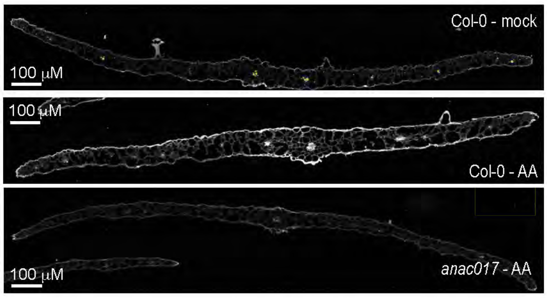


### *Tau* score

| | Col-0 CT | Col-0 AA | *anac017* CT | *anac017* AA |
|---|---|---|---|---|
| SN: | 0.86 | 0.84 | 0.82 | 0.86 |
| Xenium: | 0.97 | 0.85 | 0.97 | 0.76 |

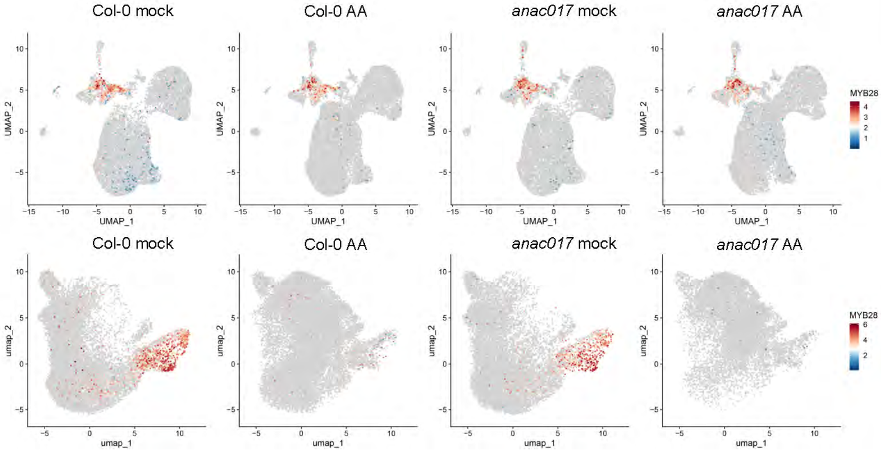

**Supplemental Figure 13. Spatial transcriptome patterns of genes encoding transcription factors.** Each panel shows an individual gene with abbreviation and gene identity. The section for each treatment represents a single section of a total of 24 sections for each treatment (12 after 3 h and 12 after 6 h of treatment with AA). The feature plots depict the UMAP projection for the snRNA-seq (top row in each panel) and spRNA-seq (bottom row in each panel). The calculated *tau* value, as a measure of expression specificity, where 1 is most specific and 0 is non-specific is shown. The feature plot clusters and cell types are designated in Figure 5. *FAMA* = bHLH transcription factor, *HB-8* = Homeobox gene 8, MYB28 and MYB29 = MYB domain protein 28 and 29, mock = control, AA = Antimycin A, SN = single nuclei.

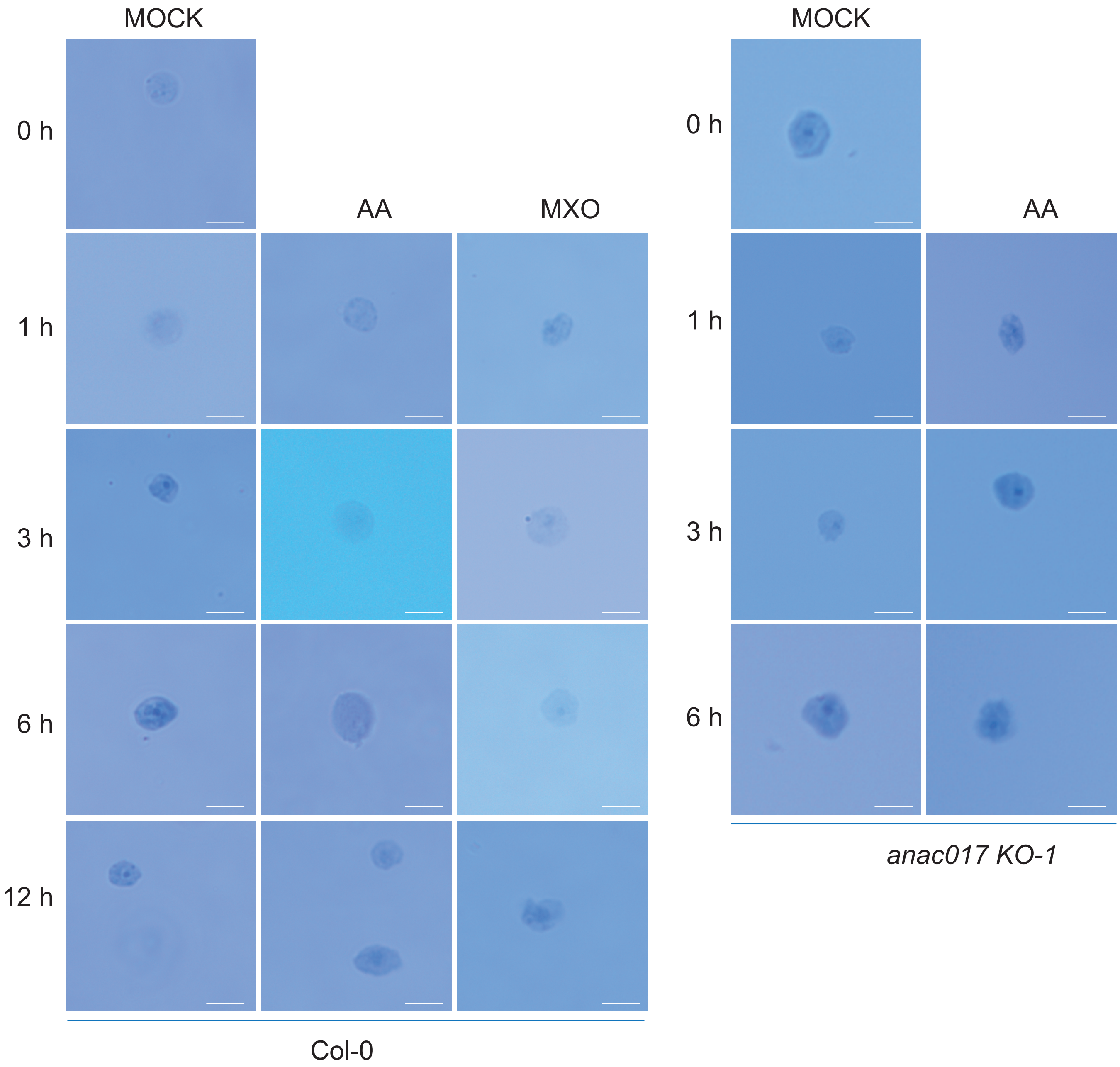


**Supplemental Figure 14. Images of nuclei after purification by flow cytometry.** The nuclei were free of debris and were observed to be intact, free of doublets and thus judged to be suitable for sequencing using microfluidics-based lysis and library preparation. Scale bar = 10 μm.
